# Biosynthesis of the α-D-Mannosidase Inhibitor (–)-Swainsonine

**DOI:** 10.1101/2024.09.26.615303

**Authors:** Shaonan Liu, Zainab Batool, Yang Hai

## Abstract

(–)-Swainsonine is a polyhydroxylated indolizidine alkaloid with potent inhibitory activity against α-D-mannosidases. In this work, we successfully reconstituted swainsonine biosynthetic pathway both in vivo and in vitro. Our study unveiled an unexpected epimerization mechanism involving two α-ketoglutarate-dependent non-heme iron dioxygenases (SwnH2 and SwnH1), and an unusual imine reductase (SwnN), which displays substrate-dependent stereospecificity such that the stereochemical outcome of SwnN-catalyzed iminium reduction is ultimately dictated by SwnH1-catalyzed C8-hydroxylation. We also serendipitously discovered that an *O*-acetyl group can serve as a detachable protecting/directing group, altering the site-selectivity of SwnH2-catalyzed hydroxylation while maintaining the stereoselectivity. Insights gained from the biochemical characterization of these tailoring enzymes enabled biocatalytic synthesis of a new polyhydroxylated indolizidine alkaloid, opening doors to the biosynthesis of diverse natural product-based glycosidase inhibitors.

## Introduction

Polyhydroxylated alkaloids, also known as iminosugar alkaloids, are a class of natural products recognized for their potent and selective glycosidase inhibitory activity (**Figure 1a**).^1^ These compounds mimic the transition states of glycoside-cleavage reactions, enabling them to bind tightly to glycosidase active sites and effectively inhibit enzymatic function.^2^ This distinctive mode of action makes polyhydroxylated alkaloids valuable chemical tools for studying glycosidase-mediated cellular processes and promising candidates for developing therapeutics targeting diseases such as cancer, diabetes, viral infections, and metabolic disorders, where glycosidases play a critical role.^3^

**Figure 1.**
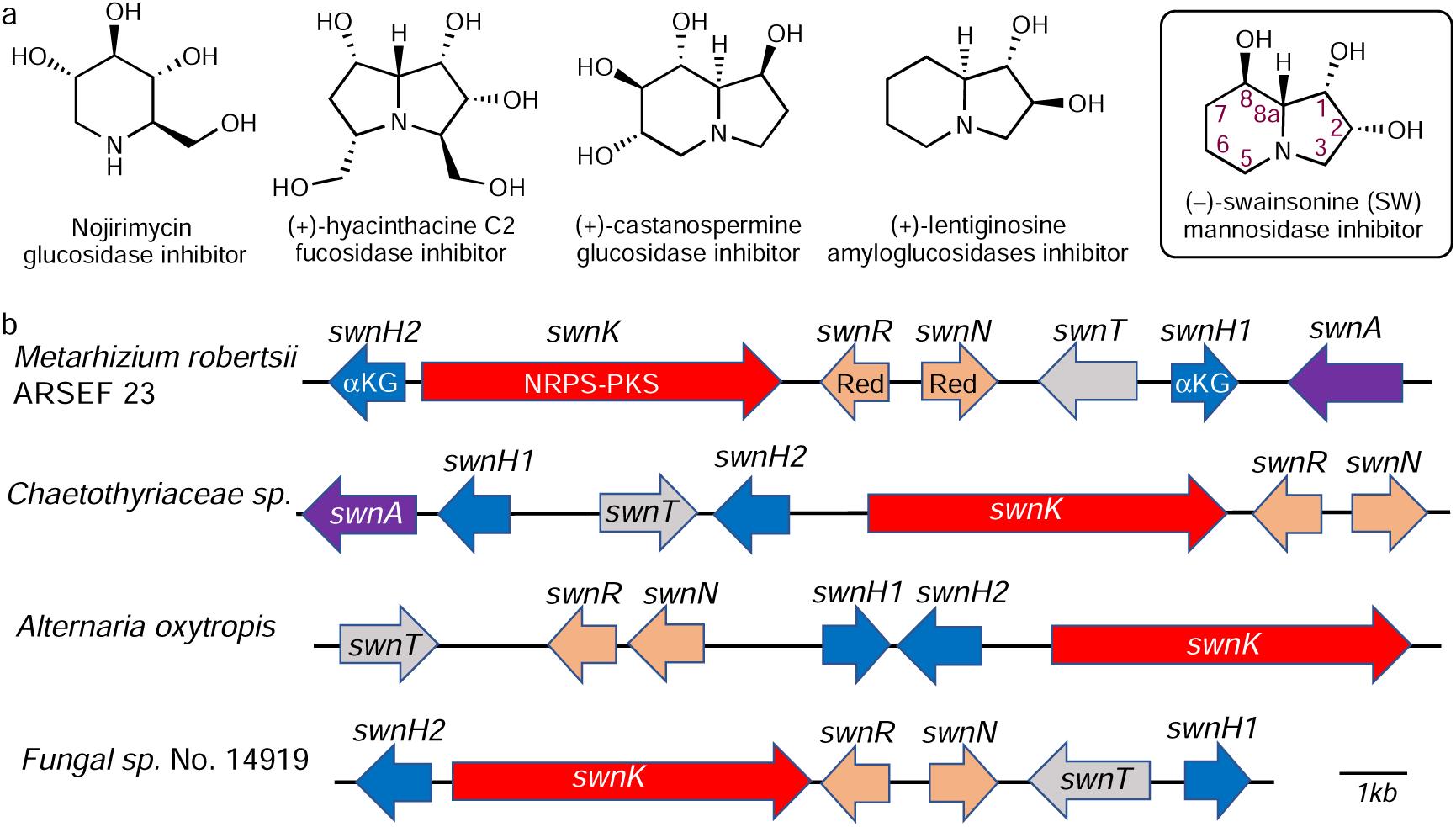
(–)-Swainsonine and other polyhydroxylated alkaloids. (A) Examples of polyhydroxylated alkaloids. (B) Swainsonine biosynthetic gene clusters from selected fungal species.

(–)-Swainsonine (SW) is a polyhydroxylated indolizidine alkaloid initially isolated from Darling pea (*Swainsona canescens*)^4^ and the fungus *Rhizoctonia leguminicola* (**Figure 1a**).^5^ It is also commonly found in spotted locoweeds (e.g. *Astragalus* and *Oxytropis*) and identified as the causative agent of “locoism” in grazing animals.^6^ The toxicity of SW arises from its inhibition of lysosomal α-mannosidase, leading to the accumulation of mannose-rich oligosaccharides, a hallmark symptom of the genetic disorder α-mannosidosis.^7^ SW is also shown as a potent inhibitor of Golgi α-mannosidase II, an enzyme essential for the maturation of *N*-linked oligosaccharides in newly synthesized glycoproteins.^8,9^ As a result, SW was the first glycoprotein processing inhibitor to undergo clinical trials for cancer treatment.^10^ Additionally, SW has been explored in the development of therapeutics for viral infections and immunological disorders.^11,12^

Due to its exceptional biological activities and unique structure, SW has garnered significant interest as a synthetic target, spurring the development of diverse synthetic strategies for its total synthesis and that of its analogues.^13–19^ However, despite decades of investigations, the mechanical details of SW biosynthesis remains elusive. Harris et al. first demonstrated that SW is derived from _L_-pipecolic acid (PA) and malonate, and identified an enigmatic epimerization evet at the ring fusion (C8a position) through feeding studies (**Figure S1**).^20–22^ The inverted stereo-configuration of C8a is important to SW’s potent α-D-mannosidase inhibition, as the inhibitory activity of (–)-8a-*epi*-swainsonine decreases by 7%.^23,24^

In 2017, Schardl and colleagues identified the conserved biosynthetic gene clusters (BGCs) for SW in fungi (**Figure 1b**). The SW BGC features a nonribosomal peptide-polyketide synthase (NRPS-PKS) hybrid (SwnK), two non-heme iron and α-ketoglutarate (Fe/αKG)-dependent oxygenases (SwnH1 and SwnH2), two NAD(P)H-dependent oxidoreductases (SwnN and SwnR), and occasionally a transporter (SwnT) and a pyridoxal 5’-phosphate (PLP)-dependent aminotransferase (SwnA).^25^ Based on predicted gene functions and Harris’s feeding study, Schardl et al. proposed a plausible biosynthetic pathway for SW, where SwnK makes the indolizidine scaffold while other enzymes (e.g. oxygenase and reductase) are responsible for the tailoring steps (**Figure S1**). However, the exact function of each enzyme remains uncharacterized. Recently, Wang and coworkers conducted gene-deletion experiments and suggested a multibranched biosynthetic pathway for SW (**Figure S1**).^26^ While this study offered insights, the multibranched proposal is contentious, as it contradicts several key findings by Harris. For instance, early feeding experiments indicated no epimerization at C1 in SW biosynthesis,^21^ whereas the multibranched pathway involves multiple C1-epimer intermediates. Moreover, several critical questions remain unresolved, such as the mechanism underlying the epimerization process.

To address these long-standing questions about the biosynthetic mechanism of SW and reconcile discrepancies in the literature, we reconstituted the complete SW biosynthetic pathway both in vivo and in vitro. We established the biochemical basis for each enzyme involved in SW biosynthesis, demonstrating that SW biosynthesis is far more complex than previously assumed. Our study uncovers that the net redox-neutral epimerization is realized through an oxidation-reduction cascade, interrupted by a hydroxylation event that dictates the stereochemical outcome of the reduction step. Insights gained from the characterization of these tailoring enzymes facilitated the biocatalytic synthesis of a novel polyhydroxylated indolizidine alkaloid, opening avenues for the future biosynthesis of diverse natural product-based glycosidase inhibitors.

## Results

### Heterologous reconstitution of SW biosynthesis in vivo

We initiated our study by reconstructing the SW biosynthetic pathway in *Aspergillus nidulans* LO7890 (**Figure 2a**).^27^ Expression of the NRPS-PKS hybrid SwnK alone did not yield any detectable products. However, supplementing the medium with L-PA or co-expressing the aminotransferase SwnA yielded 1-hydroxyindolizidine (**1**), indicating that *A*. *nidulans* LO7890 lacks sufficient endogenous L-PA and that SwnA is responsible for L-PA biosynthesis (**Figure S2**). Introducing the oxidoreductases SwnN and SwnR further increased the titer of compound **1** from 14 mg/L to 40 mg/L and 114 mg/L, respectively. To determine whether this increase resulted from enhanced L-PA availability, we dropped out *swnA* and supplemented exogenous L-PA to bypass its biosynthesis. A similar titer increase of **1** upon SwnR expression was still observed (**Figure S2**). Moreover, expressing SwnA alone was sufficient to produce L-PA in *A. nidulans,* and co-expression with SwnR or SwnN did not further elevate L-PA levels (**Figure S3**). These findings together indicate that neither SwnR nor SwnN contributes to L-PA biosynthesis and that that increased production of **1** in their presence is not due to enhanced L-PA supply.

**Figure 2.**
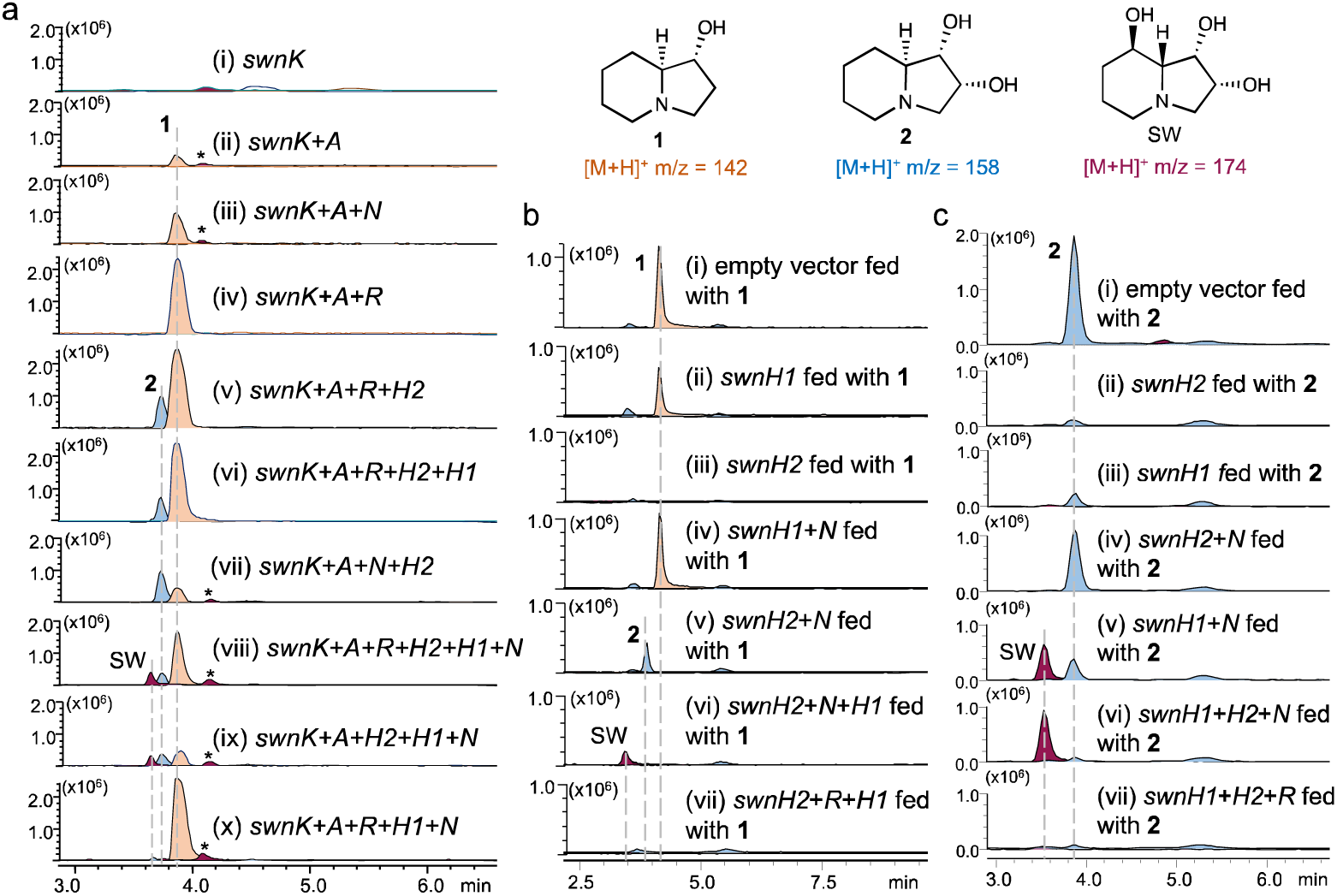
Reconstitution of swainsonine biosynthesis in *Aspergillus nidulans*. (a) LC-MS analysis of product profiles obtained from heterologous expression of different combinations of *swn* genes. The peak labeled with asterisk is a shunt product derived from SwnK product (**Figure S5**). (b) Feeding of intermediate **1** to *A. nidulans* transformed with different combination of *swn* genes. (c) Feeding of intermediate **2** to *A. nidulans* transformed with different combination of *swn* genes.

Incorporating the Fe/αKG-dependent oxygenase SwnH2 into the *swnAKRN* expression system resulted in the formation of intermediate 1,2-dihydroxyindolizidine (**2**). Subsequent introduction of oxygenase SwnH1 led to the production of the final product, SW (**Figure 2a**). All detectable compounds were isolated and characterized using NMR, high-resolution mass-spectrometry (MS), and optical rotation measurements (see **supplementary information**). Notably, we did not detect any epimers of **1** or **2** as previously reported,^26^ although conformational isomers were observed depending on the salt form (**Figure S4**).

Our results revealed that although SwnR enhances the production of **1**, it is not essential for SW biosynthesis. Removing *swnR* resulted in a reduced yield of **1**, but levels of **2** and SW were unaffected (compare trace ix with trace viii, **Figure 2a**). On the contrary, removal of *swnN* or *swnH1* completely abolished SW production, leaving only intermediates **1** and **2** detectable (compare trace vi with trace viii, and trace viii with trace ix, **Figure 2a**). Similarly, dropping out *swnH2* stalled the pathway at **1** and no advance intermediate was detected (compare trace x with trace viii, **Figure 2a**). These results indicate that SwnH2, SwnH1, and SwnN are indispensable for SW biosynthesis. The essential role of SwnN was further supported by feeding experiments. When L-PA was fed an *A. nidulans* strain co-expressing *swnKH1H2N*, SW production was observed. However, omission of *swnN* abolished SW biosynthesis (**Figure S5**). Instead, we detected a trace-level compound with the same mass-to-charge ratio (m/z = 174) as SW but a distinct retention time. Isolation and structural elucidation revealed this compound to be (*R*)-3-hydroxy-3-((*S*)-piperidin-2-yl)propanoic acid (**3’**), most likely derived from the intermediate (**3**) released from SwnK (**Figure S6**).

To gain further insights into the function of these enzymes, we fed purified intermediates (**1** and **2**) to *A. nidulans* strains expressing different combinations of *swn* genes (**Figure 2b, 2c**). In the strain expressing SwnH2 alone, compound **1** was completely consumed (trace iii, **Figure 2b**), but no downstream product (e.g. **2**) was detected unless SwnN was also present (trace v, **Figure 2b**). Similarly, feeding **2** to the strain expressing SwnH2 alone resulted in its consumption without formation of detectable new products (trace ii, **Figure 2c**). However, co-expression with SwnN rendered **2** unaltered (trace v, **Figure 2c**). Unexpectedly, SwnH1 could also transform **2**, albeit less efficiently than SwnH2 (trace iii, **Figure 2c**). Co-expression of SwnH1 and SwnN led to partial conversion of **2** into SW (trace v, **Figure 2c**), and full conversion was achieved upon addition of SwnH2 (trace vi, **Figure 2c**). These puzzling results are not readily explained by any existing biosynthetic proposals (**Figure S1**), underscoring the unexpected complexity of swainsonine biosynthesis.

### SwnA is a lysine 2-aminotransferase

To clarify the biosynthetic mechanism for SW and unambiguously assign enzyme functions, we next undertook in vitro characterization of each enzyme. We overexpressed SwnA, SwnR, SwnN, SwnH2, and SwnH1 in *E. coli.* For SwnK, we used *Saccharomyces cerevisiae* BJ5464-npgA strain as the expression system.^28^ Each protein was purified to homogeneity using metal-affinity chromatography followed by size-exclusion chromatography (**Figure S7**).

We first examined the function of SwnA, previously proposed as a lysine 6-aminotransferase.^26^ When SwnA was incubated with _L_-lysine and αKG, the expected lysine transamination reaction took place (**Figure S8a**). However, chemical reduction of the reaction product (i.e. dehydropipecolic acid) using NaBH_3_CN resulted in a racemic mixture of pipecolic acids (**Figure S8b**). The loss of the L-configuration at the Cα position indicated that the product of SwnA-catalyzed reaction is 1-piperideine-2-carboxylate (P2C), rather than the previously proposed 1-piperideine-6-carboxylate (P6C). If P6C was the true product, chemical reduction would have retained the L-configuration, yielding enantiomerically pure L-pipecolic acid.

To further confirm this result, we repeated the reaction using differently ^15^N-labeled _L_-lysine isotopologs. After Fmoc-derivatization and mass analysis, the resulting ketoacid product showed a +1 Da mass shift only when ^15^N_ε_-_L_-lysine was used as the substrate, but not with ^15^N_α_-_L_-lysine (**Figure S8c**). Similarly, feeding ^15^N_ε_-_L_-lysine to *A*. *nidulans* strains expressing the pathway led to partial isotope-incorporation into L-PA and **1**, whereas no mass shift was observed with ^15^N_α_-_L_-lysine (**Figure S9**). Together, these results confirm that the α-amino group of L-lysine is lost during the SwnA-catalyzed transamination, establishing that SwnA is a lysine 2-aminotransferase rather than a lysine 6-aminotransferase, as previously proposed.^26^

We next assessed the potential roles of SwnN and SwnR as P2C reductases using SwnA-coupled enzymatic assays (**Figure S10**). While increasing the concentrations of SwnN and SwnR to 50 µM did result in low levels of pipecolic acid production, the observed activities were insufficient to be considered physiologically relevant. Moreover, SwnR exhibited poor enantioselectivity, and SwnN primarily produced the undesired enantiomer, _D_-PA. These findings suggest that neither enzyme functions as a P2C. Instead, we propose that an endogenous imine reductase in the fungal host likely mediates the reduction of dehydropipecolic acid to L-PA during swainsonine biosynthesis.

### Reconstitution of indolizidine backbone biosynthesis of SW in vitro

Following the function characterization of SwnA, we next investigated SwnK, the enzyme responsible for constructing the indolizidine scaffold of SW. When provided with the necessary substrates and cofactors (i.e. malonyl-CoA, NADPH, and pipecolic acid), SwnK readily synthesized **1** in vitro, showing a clear preference for _L_-PA over _D_-PA as the amino acid substrate (**Figure S11**). This in vitro activity is consistent with our in vivo result that **1** was produced with only SwnK expressed and L-PA provided. These results also establish that SwnK, rather than the previously proposed SwnN,^26^ dictates the stereochemistry of the C1-OH group in **1**.

To investigate why SwnR significantly enhances the fermentation titer of **1**, we conducted metabolomics analysis using GC-MS. Alongside **1,** we identified a new metabolite, 5,6,7,8-tetrahydroindolizine (designated as **4’**), in cultures of *A. nidulans* co-expressing SwnK and SwnA (**Figure S12**). However, when either SwnR or SwnN was included, the levels of **4’** were drastically reduced. We propose that **4’** arises from intermediate **4**, an iminium precursor of **1** (**Figure S6**). In the absence of prompt reduction, the cyclic iminium moiety in **4** undergoes dehydration and isomerization, yielding the shunt product **4’**. Although SwnK alone is able to reduce **4** to generate **1**, efficient conversion in vivo likely depends on a dedicated imine reductase. Given that inclusion of *swnR* resulted in the highest production of **1**, we propose that SwnR serves as the primary imine reductase for **4**, enhancing the yield of **1** by preventing formation of the shunt product **4’**. In the absence of SwnR, SwnN can partially compensate for this activity, albet with reduced efficiency.

### Elucidating the epimerization process of SW biosynthesis

To unravel the enzymatic transformations in the tailoring phase of SW biosynthesis, we first investigated SwnH2, which was shown to act on both **1** and **2** in vivo (**Figure 2c** and **Figure 2d**). When supplied with the required cofactor and co-substrates (e.g. αKG, ferrous sulfate, sodium ascorbate), SwnH2 efficiently oxidized **1** to **2**, while concurrently generating a new compound, designated **5** (trace i, **Figure 3a**). Structural characterization identified **5** as the iminium derivative of **1** (**Table S6**), and the ratio between **2** to **5** was determined to be approximately 82:18 (**Figure S13**).

**Figure 3.**
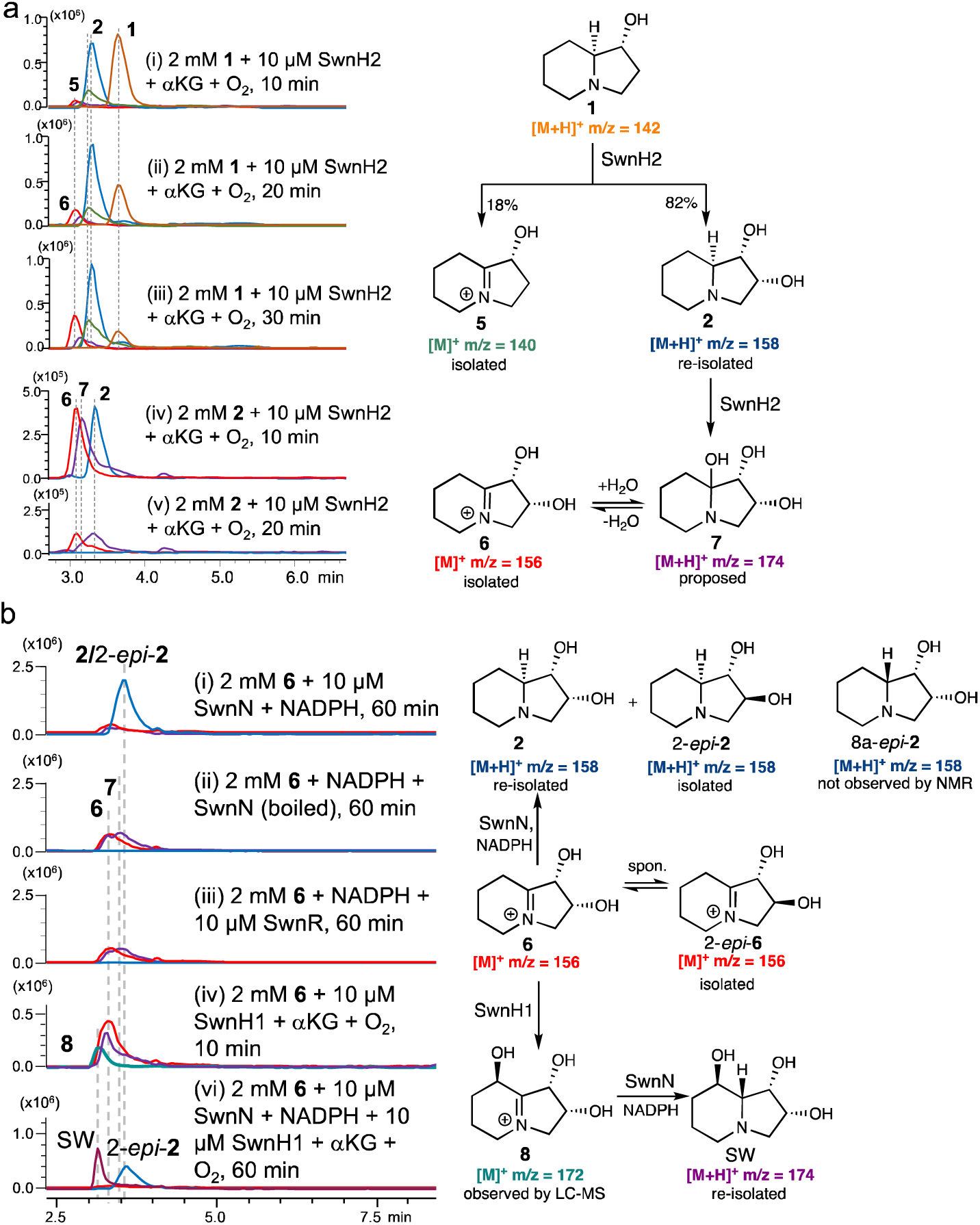
Reconstituting the biosynthesis of swainsonine in vitro. (a) SwnH2 is a bifunctional oxygenase catalyzing oxidation of both **1** and **2**. (b) Unexpected substrate-dependent stereoselecitivty exhibited by SwnN.

In agreement with in vivo observations, SwnH2 also catalyzed the oxidation of **2** (trace iv and v, **Figure 3a**), yielding two additional compounds, **6** and **7**, with m/z(+) values of 156 and 174, respectively. Isolation and structural analysis revealed that **6** is the iminium derivative of **2** (**Table S7**), while **7** was identified as the corresponding equilibrated hemiaminal, based on its reduction by NaBD_3_CN (**Figure S14**). These findings demonstrate that SwnH2 is a bifunctional Fe/αKG-dependent oxygenase capable of catalyzing both C2-hydroxylation and a formal amine desaturation reaction.

During the isolation process, we noticed that **6** spontaneously isomerized, resulting in the isolated product as a mixture of **6** and its epimer, 2-*epi*-**6** (**Figure S15**). The inherent instability likely accounts for the absence of **6** in in vivo experiments (e.g. trace v, **Figure 3a**). To asse whether **6** is a substrate for enzymatic reduction, an isolated sample of **6** (containing 33 mol% 2-*epi*-**6**) was incubated with SwnN and NADPH. This iminium reduction produced a product with the mass-to-charge ratio and retention time resembling **2** (trace i, **Figure 3b**). Isolation and structural analysis revealed this prodcut to be a ∼1:1 mixture of two diastereomers: **2** and its epimer, 2-*epi*-**2** (**Figure S16**). Control reaction using either SwnR or heat-inactivated SwnN failed to reduce **6,** confirming that NADPH alone is insufficient to reduce **6** under these assay conditions (trace ii and iii, **Figure 3b**). We hypothesize that 2-*epi*-**2** arose from 2-*epi*-**6**, an artifact generated during the isolation of **6**. Importantly, the ability of SwnN to reduce **6** back to **2** is consistent with our in vivo feeding studies, in which **2** was consumed by SwnH2 but accumulated when both SwnH2 and SwnN were present. Furthermore, the absence of 8a-*epi*-**2** in the isolated enzymatic products indicates that SwnN specifically reduces **6** via the α-face.

Recognizing that reduction of **6** is not the appropriate timing for epimerization, we explored an alternative biosynthetic sequence in which C8-hydroxylation precedes iminium reduction. When the same sample of isolated **6** was treated with SwnH1 and the requisite cofactor and co-substrates, rapid consumption of **6** was observed. A new species, designated as **8**, emerged at early time points (trace iv, **Figure 3b**). Chemical reduction of **8** with NaBH_4_ indicated that it is an iminium compound (**Figure S17**). When SwnN and NADPH were included in the reaction mixture, enzymatic reduction of **8** was achieved, yielding the final product SW (trace v, **Figure 3b**). Structural analysis of the reaction products also revealed the presence of 2-*epi*-**2**, which we attribute to reduction of 2-*epi*-**6**—an artifact carried over from the sample (**Figure S18**).

The above results support a revised swainsonine biosynthetic pathway in which SwnH2 first generates the iminium intermediate **6**, followed by C8-hydroxylation by SwnH1 to form **8**, which is then stereoselectively reduced by SwnN to complete SW biosynthesis, thereby resolving a previously unexplained stereoinversion mechanism. Importantly, since SwnN also reduces **6** to **2**, these findings also indicate that SwnN is a rare example of imine reductases capable of setting opposite stereochemistry depending on the substrate. This behavior stands in contrast to the non-stereospecific hydride transfer catalyzed by many sugar epimerases, such as UDP-hexose-4-epimerase, which reduces transient keto intermediates in a non-stereospecific manner to yield either UDP-galactose or UDP-glucose.^29^

### Timing of epimerization is determined by C8-hydroxylation

Having established the intermediacy of **6** in SW biosynthesis, we next focused on **5**, another iminium compound derived from **1**, to assess its potential role in SW biosynthesis. Compound **5** was neither further oxidized by SwnH2 (trace i, **Figure 4a**) nor reduced by SwnR (**Figure S19**). Instead, under these conditions, **5** spontaneously tautomerized to its ketone isomer, **9** (**Figure S19**). In contrast, SwnN successfully reduced **5** using NADPH (trace ii, **Figure 4a**), producing **1** as a single diastereomer product, with no detectable formation of 8a-*epi*-**1** in the isolated product.

**Figure 4.**
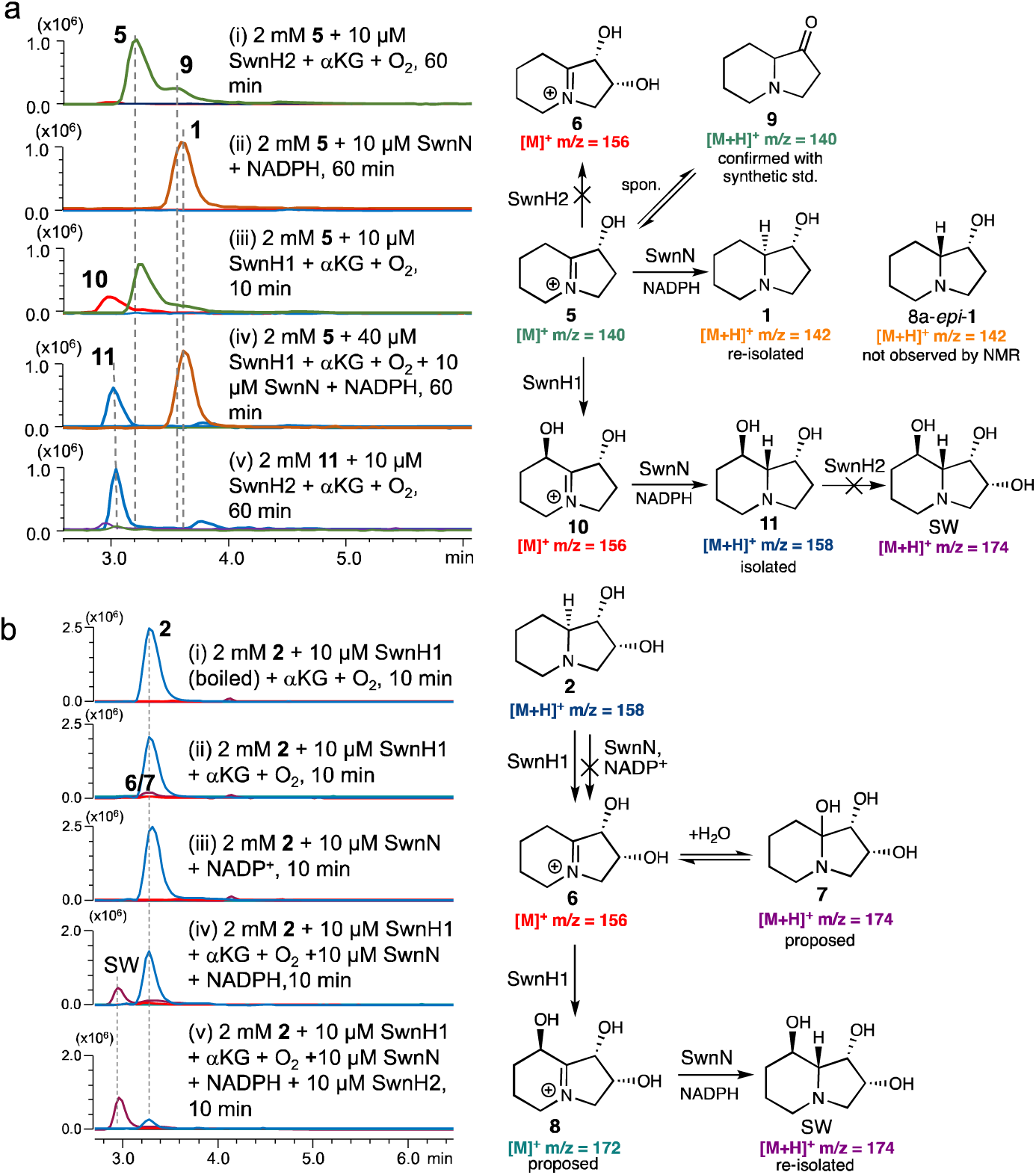
Additional insights gained from biochemical characterization of the tailoring enzymes. (a) Iminium compound **5** is not an on-pathway intermediate, but can be recycled by SwnN to regenerate **1**; (b) SwnH1 is also capable of catalyzing amine desaturation of **2.**

Interestingly, SwnH1 could further hydroxylate **5** to produce a new iminium compound **10**, albeit with low catalytic efficiency (trace iii, **Figure 4a**). When **10** was further reduced by SwnN, a new diol compound, **11**, was isolated and identified as 2-deoxy-swainsonine (trace iv, **Figure 4a**). However, SwnN-catalyzed reduction of **5** outcompeted SwnH1-catalyzed hydroxylation. Therefore, elevated enzyme concentrations (≥ 40 µM) of SwnH1 were required to enable successful isolation of **11**, confirming that **5** is not a native substrate of SwnH1. Furthermore, **11** was not a kinetically competent intermediate for SW biosynthesis, as incubation with SwnH2 produced only trace amounts of a triol product (trace v, **Figure 4a**). Overall, these results demonstrate that **5** is not an on-pathway intermediate for SW, but it can be recycled by SwnN to regenerate intermediate **1**.

A comparison of SwnN-catalyzed iminium reductions indicated that while SwnN establishes the stereochemistry at the ring fusion, the ultimate outcome is dictated by the C8-hydroxylation installed by SwnH1. When SwnH1 introduces the C8-OH group on the β-face, SwnN preferentially reduces the substrates from the same face, yielding C8a-epimerized products, such as **11** and SW. In contrast, in the absence of SwnH1-installed C8-OH group (as observed with **5** and **6**), SwnN reduces the iminium substrates from the α-face.

To understand the structural basis for this unusual substrate-dependent stereoselectivity, we generated an AlphaFold model of SwnN in complex with NADP^+^,^30^ and docked various products (e.g. **1**, **2**, **11**, and SW) into its active site pocket (**Figure S20**). The docking models suggest that SwnN recognizes the β-face oriented C8-OH group through a hydrogen bond, thereby positioning the substrate for β-face reduction. Without this anchoring interaction, substrates appear to be in a flipped-ring orientation, utilizing the C1-OH group to form similar hydrogen bonds, which in turn leads to α-face reduction. Although this model is supported by our mutagenesis experiments (**Figure S21**), further biochemical and structural characterization, including cocrystal structural determination of SwnN with a bound product, will be necessary to fully elucidate its unusual catalytic specificity.

### SwnH2 and SwnH1 are bifunctional enzymes

After ruling out the pathway branching from **5** to SW, we turned our attention to another unexpected finding from our in vivo studies. Specifically, SwnH1 was shown to utilize compound **2** in vivo (trace iii, **Figure 2c**), and in the presence of both SwnH1 and SwnN, **2** was partially converted to SW (trace v, **Figure 2c**). While SwnN alone was unable to oxidize **2** (trace iii, **Figure 4b**), SwnH1 successfully oxidized it to generate **6** and **7** (trace ii, **Figure 4b**), although the yield of **6** was insufficient for structural confirmation. In a one-pot reaction containing both SwnN and SwnH1, a clear transformation of **2** to SW was observed (trace iv, **Figure 4b**). With SwnH2 was also included, complete conversion of **2** to SW was achieved (trace v, **Figure 4b**), consistent with results from our in vivo feeding experiments. Notably, no 2-*epi*-**2** was detected in the isolated product (**Figure S18**), further confirming that 2-*epi*-**2** and its precursor, 2-*epi*-**6**, are artifacts that would not form when intermediate **6** is generated in situ.

These results establish that SwnH1, like SwnH2, is also a bifunctional Fe/αKG-dependent oxygenase. However, they must proceed through distinct mechanistic pathways. SwnH2 cleaves two C-H bonds at both C2 and C8a exclusively from the α-face, likely because its ferryl intermediate approaches the substrate from this face. The bifurcation between C2 and C8a is probably governed by the relative proximity of these hydrogens to the reactive ferryl species. In contrast, SwnH1 installs the C8-OH group on the β-face, suggesting that its ferryl intermediate approaches the substrate from the opposite face. Accordingly, hydrogen tom transfer (HAT) at C8a is feasible for SwnH2 but not SwnH1 (**Figure S22**).

This hypothesis is supported by enzymatic assays using C8a-deuterium-labeled substrate **2**. A primary kinetic isotope effect was observed with SwnH2, but not with SwnH1, indicating that C8a-H abstraction is specific to SwnH2 (**Figure S23**). To test whether SwnH1-catalyzed amine desaturaiton of **2** an enamine-imine tautomerization step, we performed the reaction in D_2_O. However, no deuterium incorporation was detected (**Figure S24**), arguing against this mechanism. To probe the possibility of an aminyl radical intermediate (**Figure S25**), we synthesized a radical clock substrate analogue, compound **2’**, and evaluated its reactivity with both SwnH1 and SwnH2. A ring-opened product–indicative of radical rearrangement–was observed with SwnH1, but not with SwnH2 (**Figure S26**). Taken together, these results indicate that although both enzymes catalyze the same formal amine desaturation reaction, they proceed via mechanistically distinct pathways: SwnH2 employs a classical oxygen-rebound mechanism involving initial α-face ferryl attack, while SwnH1 likely initiates desaturation via a β-face-directed aminyl radical pathway. Further structural and mechanistic studies will be necessary to fully elucidate the basis of their divergent catalytic strategies.

### Directing SwnH2 function by a protecting/directing group

Acetylation is a common protective strategy in natural product biosynthetic pathways, resulting in cryptic acetylated biosynthetic intermediates that are unforeseeable without biosynthetic investigation.^31,32^ Accordingly, we also hypothesized that a similar cryptic acetylation-deacetylation event may be involved in swainsonine biosynthesis. To test this hypothesis, we synthesized *O*-acetylated-**1** (**Ac**-**1**) and evaluated its relevance to SW biosynthesis using purified enzymes. No reaction was observed with SwnH1 (trace i, **Figure 5**). However, SwnH2 successfully converted **Ac**-**1** into an acetylated diol product (**Ac**-**12**) and the shunt product **5** (trace ii and iii, **Figure 5**). Structural characterization of **Ac**-**12** revealed an α-face-oriented hydroxyl group at C7. These results eliminate the possibility of **Ac-1** serving as a biosynthetic intermediate in SW biosynthesis. Nonetheless, the stereochemistry of the C7-OH group in **Ac**-**12** supports our proposed mechanism, where substrates approach the non-heme iron center in SwnH2 via the α-face (**Figure S19**).

**Figure 5.**
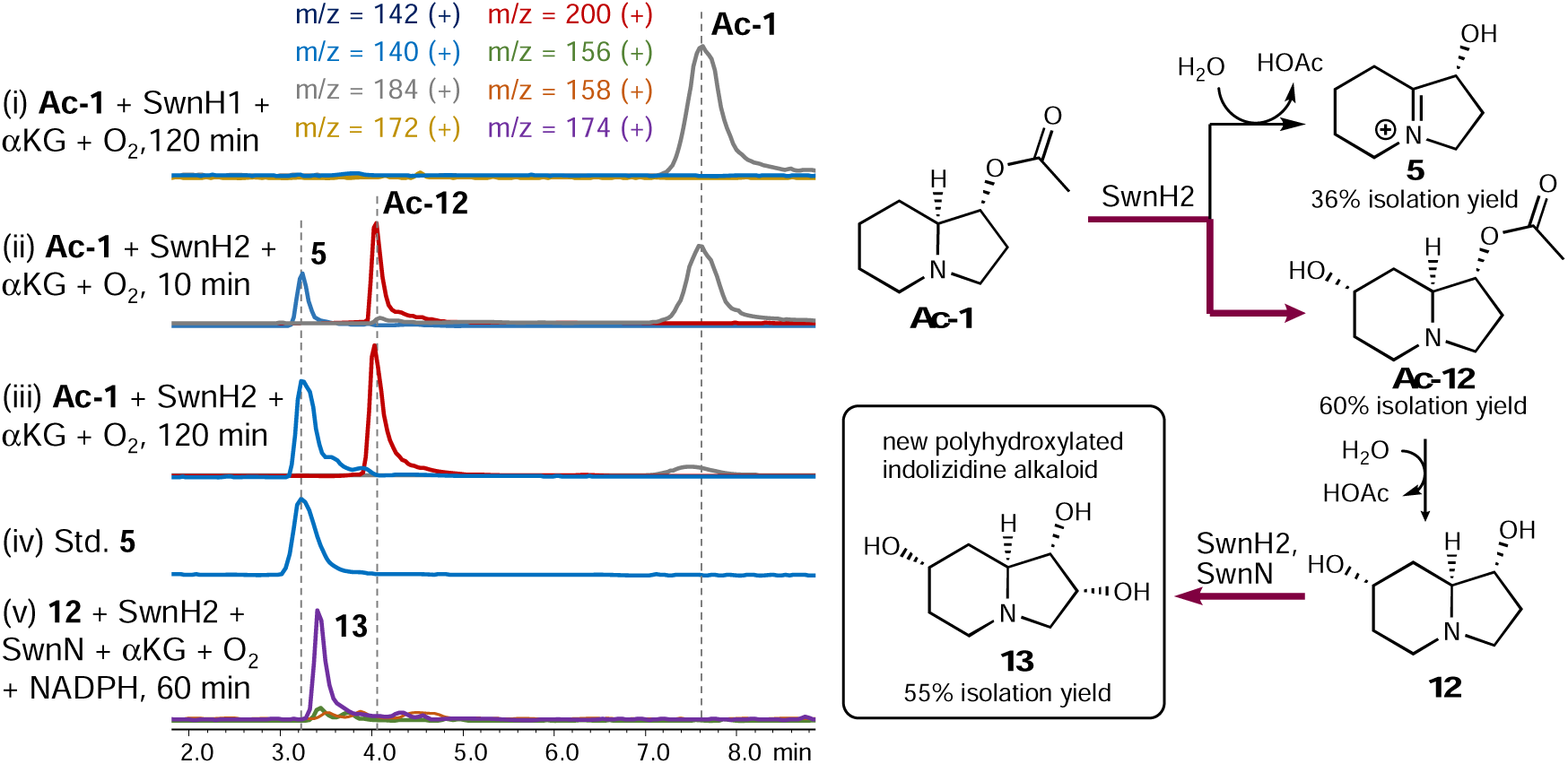
Biocatalytic synthesis of a novel polyhydroxylated indolizidine alkaloid enabled by a protective acetyl group-directed biosynthesis.

Given the removable nature of the *O*-acetyl group, we next subjected the deprotected diol, **12**, to further oxidative tailoring by SwnH2. SwnN and NAPDH were included in the reaction to quench any unstable iminium intermediates. From this biocatalytic cascade, we successfully isolated a triol compound, **13** (trace v, **Figure 5**). Structural analysis identified **13** as a novel polyhydroxylated indolizidine alkaloid, isomeric to SW, with the same C2-OH group but lacking C8a epimerization. The absence of epimerization supports our gatekeeping mechanism, which posits that epimerization occurs only after SwnH1 installs the C8-OH group.

## Discussion

Polyhydroxylated alkaloids are renowned for their potent glycosidase inhibitory activities and potential therapeutic applications, yet their biosynthetic pathways remains largely unexplored. Our study of the SW biosynthesis marks the first complete elucidation of the biosynthetic pathway of any polyhydroxylated alkaloid to date. Contrary to the multibranched pathway proposed by Wang et al.,^26^ our findings support a linear biosynthetic mechanism in agreement with feeding studies conducted four decades ago.^21^ Briefly, the NRPS-PKS hybrid SwnK constructs the indolizidine backbone from L-pipecolic acid, while the tailoring enzymes (SwnH2, SwnH1, and SwnK) catalyze late-stage modifications: C2-hydroxylation, amine desaturation, C8-hydroxylation, and imine reduction, culminating in the production of SW (**Figure 6**).

**Figure 6.**
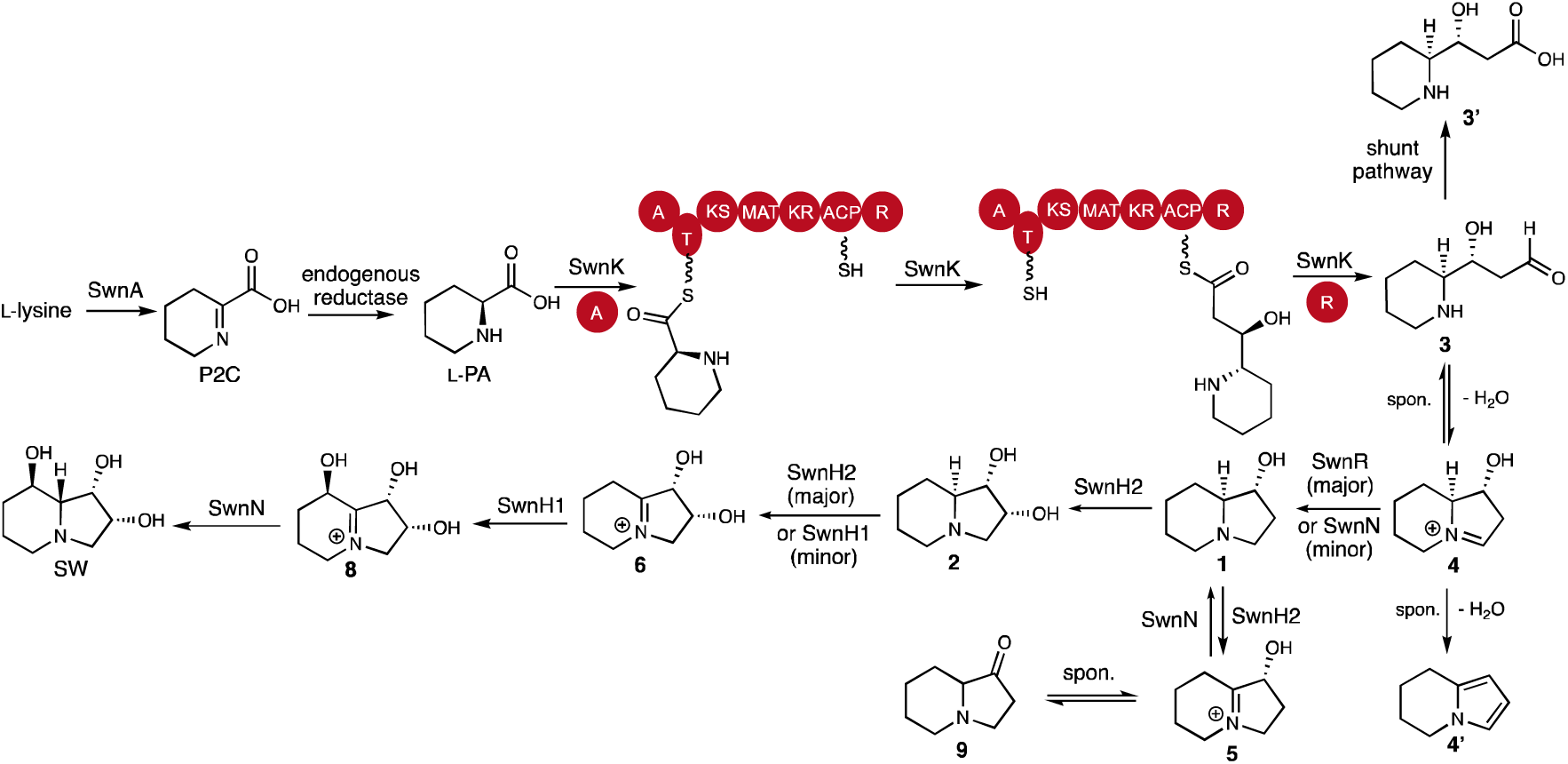
Revised swainsonine biosynthetic pathway based on all findings from this study.

The sequential oxidation-reduction process is a common chemical logic for stereoinversion in both chemical synthesis and biosynthesis.^33–35^ The epimerization process in SW biosynthesis elucidated here is reminiscent of the conversion of (*S*)-reticuline to (*R*)-reticuline via 1,2-dehydroreticuline in morphine biosynthesis.^36^ Unlike reticuline epimerase, a fusion protein consisting a cytochrome P450 domain and an aldo-keto reductase domain, SW biosynthesis recruits three separate enzymes to achieve stereoinversion. Notably, the stereochemical outcome of imine reduction is dictated by SwnH1-catalyzed hydroxylation, adding an additional layer of stereochemical control at the ring fusion. This complexity enables a single reductase, SwnN, to function in both epimerization and recycling of the byproduct **5**. Reductases exhibiting such substrate-dependent stereospecificity are exceedingly rare. While a few ketoreductases within PKS assembly lines show similar behaviors,^37–39^ SwnN is the first standalone imine reductase capable of substrate-dependent stereospecific reduction.

The serendipitous enzymatic synthesis of (Ac)-**12** and its derivative **13** highlights the role of an artificially attached *O*-acetyl group, which functions not only as a protecting group to prevent hydroxylation at C2, but also as a directing group, altering the site-selectivity of SwnH2 while maintaining stereoselectivity. We propose that the bulky acetyl group induces a subtle shift in substrate binding, rendering the C_2_-H bond inaccessible while positioning the C7 site closer to the ferryl intermediate.^40^ Although the detailed mechanism of SwnH2 remains to be fully elucidated, our ability to successfully alter the site-selectivity of SwnH2 using a detachable protecting group is noteworthy. This approach suggests a general strategy for biocatalysis, where deploying detachable protecting groups to direct enzyme activity may serve as a practical alternative to engineering the enzymes themselves.^40^

Finally, our study of SW biosynthesis sheds light on the biosynthetic pathway of slaframine, a structurally related indolizidine alkaloid (**Figure S27**). Like SW, slaframine is also derived from L-pipecolic acid, and the fungus *R*. *leguminiocola* produces both SW and slaframine. Harris et al. proposed that slaframine and SW share a common early biosynthetic pathway, with ketone indolizidine **9** serving as a shared precursor.^21^ However, our findings show that **9** is not a biosynthetic intermediate for SW, and the C1-OH stereochemistry in SW is determined by SwnK. These results suggest that the existence of a separate NRPS-PKS hybrid dedicated to slaframine biosynthesis in *R*. *leguminiocola*, likely similar to SwnK but with a distinct ketoreductase domain that determine the opposite stereochemistry of the C1-OH. Here we propose a revised biosynthetic pathway for slaframine, informed by our discoveries in SW biosynthesis (**Figure S27**).

## Conclusion

In conclusion, we have elucidated the complete biosynthesis of swainsonine, solving a biosynthetic puzzle that has persisted for decades. Our comprehensive characterization of SwnH2, SwnH1, and SwnN unveiled an unusual epimerization mechanism: it follows a classic oxidation-reduction sequence interrupted by a hydroxylation event that ultimately determines the stereochemical outcome of the reduction step. We demonstrated that reductase SwnN exhibits unusual substrate-dependent stereoselectivity, while SwnH1 and SwnH2 are bifunctional (Fe/αKG)-dependent oxygenases and provided mechanistic insights. Additionally, we directed the site-selectivity of SwnH2 by deploying a protective acetyl group on the substrate. This strategy enabled the biosynthesis of a new polyhydroxylated alkaloid. Our study highlights the potential of harnessing these biosynthetic enzymes for the discovery and development of new natural product-based glycosidase inhibitors.

## Data availability

All data supporting the findings of this study are presented in the main text and the Supplementary Information file. NCBI (https://www.ncbi.nlm.nih.gov/) accessions codes are referenced in the Supplementary Information file. All data is available from the corresponding author upon request.

## Acknowledgements

This work is supported by the University of California Cancer Research Coordinating Committee Grant C24CR7230 and NIH grant R35GM151205. The authors thank Dr. Christopher Schardl (University of Kentucky) for helpful discussions, and Prof. Berl R. Oakley (University of Kansas) for providing A. nidulans strain LO7890, and Prof. Yi Tang (University of California, Los Angeles) for providing the S. cerevisiae BJ5464-npgA strain.

## Author contributions

Y.H. and S.L. designed the overall research. S.L. and Z.B. performed all biochemical experiments. Y.H. and S.L. wrote the manuscript.

