## Supplementary Information for "Biosynthesis of the α-D-Mannosidase Inhibitor (–)-Swainsonine"

FIGURE S13. DETERMINING THE RATIO BETWEEN 2 AND 5 DURING THE INITIAL REACTION OF SWNH2-CATALYZED OXIDATION OF 1... 42

### 1. Method.

#### 1.1 Materials.

Glucose-6-phosphate, Glucose-6-phosphate dehydrogenase (G6PD), d7-D-glucose, NaBH3CN, FmocCl and Dowex® 1x4-400 resin were purchased from Sigma-Aldrich. d9-rac-pipecolic acid was sourced from C/D/N Isotopes. D-pipecolic acid, L-pipecolic acid, Salicylaldehyde, and Acetic acid-d were obtained from Combi-Blocks. NADPH, NADP, and NADH were acquired from AmBeed. Marfey's reagent [Nα-(5-Fluoro-2,4-dinitrophenyl)-L-alaninamide] was purchased from TCI. NaCD3CN, L-Lysine·2HCl (ε-$^{15}$N, 98%), and L-Lysine·2HCl (α-$^{15}$N, 98%) were purchased from Cambridge Isotope Laboratories. All other chemicals and solvents were obtained from Fisher Scientific.

#### 1.2 General Methods.

Enzymatic reactions were monitored by LC-MS using on a Shimadzu 2020 LC-MS-PDA (a Cosmosil C18 AR-II column (5.0 µm, 10 ID X 250 mm, Shodex)) using positive and negative mode electrospray ionization with an isocratic flow of 1% MeCN−H2O supplemented with 0.1% (v/v) formic acid in 6 min followed by 98% MeCN for 5 min with a flow rate of 0.5 mL/min. Compounds were monitored by thin layer chromatography (TLC) carried out on MilliporeSigma Aluminium TLC plates (silica gel 60 coated with $F_{254}$) using basic aqueous potassium permanganate as developing agent. NMR spectra were recorded on a Bruker Advance III 500 MHz NMR spectrometer. The spectra were calibrated by using residual undeuterated solvents (for $^{1}$H NMR) and deuterated solvents (for $^{13}$C NMR) as internal references: undeuterated methanol ($\delta_H$ = 3.31 ppm) and methanol-$d_4$ ($\delta_C$ = 49.00 ppm); The following abbreviations are used to designate multiplicities: s = singlet, d = doublet, t = triplet, q = quartet, m = multiplet, br = broad. Optical rotation was recorded on a Rudolph Research Laboratory AUTOPOL IV polarimeter. UV-Vis spectra were recorded on a Shimadzu 1601 spectrophotometer. Gas chromatography-mass spectrometry (GC-MS) analyses were carried out using a Waters GCT Premier GC TOF (EIHRMS), with an Agilent 7890 A gas chromatograph and a DB5-MS column (30 m × 0.25 mm I.D., 0.25 µm film). The oven temperature program started at 40°C, then ramped at 20°C/min to 300°C, and held at 300°C for 7 minutes. Accurate mass measurements and fragmentation patterns were used to confirm the identity of the analytes. HRMS were recorded on a Shimadzu LCMS-9030 UHPLC-QTOF (ESI-HRMS).

#### 1.3 Strains and culture conditions

*Aspergillus nidulans* strain LO7890 was a gift from Prof. Berl R. Oakley (University of Kansas) and was used for in vivo pathway reconstruction.[1] Escherichia coli strain XL10-Gold was used for cloning, while BL21 (DE3) were used for expression. The *Saccharomyces cerevisiae* strain JHY686 was used as the yeast host for in vivo homologous recombination to construct the *A. nidulans* expression plasmids,[2] while the *S. cerevisiae* BJ5464-NpgA strain was used for expression of SwnK.[3]

*Aspergillus nidulans* LO7890 was grown at 28 °C on CD medium (10 g/L of glucose, 1X nitrate salts, 0.1% v/v of trace elements, pH 6.5, and 20 g/L of agar for solid cultivation); or in CD-ST media (20 g/L of starch, 20 g/L of casamino acids, 1X nitrate salts, 0.1% v/v of trace elements, pH 6.5) for heterologous expression of the gene cluster, compound production, and RNA extraction.[4] For preparation of 20X nitrate salts, 120 g of $NaNO_3$, 10.4 g of KCl, 10.4 g of $MgSO_4{\bullet}7H_2O$, 30.4 g of $KH_2PO_4$ were dissolved in 1 L of double distilled water. For preparation of the trace element solution, 2.20 g of $ZnSO_4{\bullet}7H_2O$, 1.10 g of $H_3BO_3$, 0.50 g of $MnCl_2{\bullet}4H_2O$, 0.16 g of $FeSO_4{\bullet}7H_2O$, 0.16 g of $CoCl2{\bullet}5H_2O$, 0.16 g of $CuSO_4{\bullet}5H_2O$, and 0.11 g of $(NH_4)_6Mo_7O_{24}{\bullet}4H_2O$ were dissolved in 100 mL of double-distilled water, and the pH was adjusted to 6.5. All Escherichia coli strains were cultured in LB media at 37 °C. Yeast strains were cultured in YPD media (yeast extract 1%, peptone 2%, glucose 2%) at 28 °C.

##### 1.4 General DNA manipulation techniques

PCR was performed using Q5 High-Fidelity DNA Polymerase (NEB). The gene-specific primers are listed in Table S1. The sequences of PCR products were confirmed by DNA sequencing (Laragen Inc.). Escherichia coli XL10-Gold (Stratagene) was used for plasmid propagation. DNA restriction enzymes (New England Biolab) were used according to the protocol provided by the manufacturer. Genomic DNA of *Metarhizium robertsii* ARSEF 23 was extracted by using the *Quick*-DNA Fungal/Bacterial Miniprep Kit (ZYMO Research).

For isolation of RNA from *A. nidulans* transformants, the RNA extraction steps were performed using the PureLink™ RNA Mini Kit (Invitrogen) following the manufacturer's instructions. Residual genomic DNA in the extracts was digested by DNase I (2 U/µL) (Invitrogen) at 37 °C for 4 hours. SuperScript III FirstStrand Synthesis System (Invitrogen) was used for cDNA synthesis with Oligo-dT primers following directions from the user manual. The intron free ORFs of swnA, swnK, swnH1, and swnH2 were amplified by PCR and ligated to linear expression vector.

##### 1.5 Protein overexpression in *E. coli* and purification

*E. coli* BL21 (DE3) transformants harboring the corresponding plasmids were grown overnight in LB medium containing 50 µg/mL kanamycin at 37 °C. Each 1 litter fresh LB medium supplemented with 50 µg/mL kanamycin was inoculated with 5 mL of the overnight starting culture, and incubated at 37 °C and 230 rpm until optical density $OD_{600}$ reached 0.8. The expression was induced by 200 μM IPTG and the cell cultures were left grown at 16 °C for 20h. Cells were harvested by centrifugation and resuspended in cell lysis buffer [50 mM $K_2HPO_4$ (pH 7.5), 10 mM imidazole, 300 mM NaCl, 5% glycerol]. Cells were lysed by sonication on ice and the cell lysate was cleared by centrifugation at 26,000 g for 60 min at 4 °C. The supernatant was incubated with $Ni^{2+}$-NTA resin for 30 min at 4 °C and then the slurry was loaded onto a gravity column. The resin was washed and eluted with increasing concentrations of imidazole (from 10 mM to 500 mM) in cell lysis buffer. The fractions were examined by SDS-PAGE gels. Pure fractions were concentrated by Amicon concentrators (Millipore), supplemented with 10% glycerol and stored at -80 °C.

The Non-heme Iron(II)-dependent oxidase (SwnH1 or H2) was incubated with EDTA at a final concentration of 3 mM on ice for 30 minutes to chelate the active-site metal. The solution was then dialyzed at 4 °C using 20 kDa Slide-A-Lyzer Dialysis Cassettes against 2 L of dialysis buffer (50 mM potassium phosphate, pH 7.5, 1 mM DTT) for 12 hours to remove EDTA and dissociated metal ions. Following dialysis, 10% (v/v) glycerol was added to stabilize the protein, which was subsequently flash-frozen in liquid nitrogen and stored at -80 °C. Protein was supplemented with 10% glycerol and stored at -80 °C.

Protein concentration was determined by measuring the absorbance at 280 nm. All proteins were expressed and purified using the same procedure as described.

##### 1.6 Protein overexpression in yeast and purification

For protein expression in *S. cerevisiae* BJ5464-NpgA, the intron-less gene, was amplified by PCR and ligated to linear expression vector (XW55 or XW06 or XW02). SwnA was inserted into XW02 vector. SwnK was inserted into XW55 vector. SwnR gene and /or SwnN was inserted into XW06 vector. Each plasmid was introduced into *S. cerevisiae* BJ5464-NpgA strain with Frozen E-Z Yeast Transformation Kit (Zymo) selected by dropout media, uracil-dropout for XW55 vector, leucine-dropout for XW02 vector, tryptophandropout for XW06 vector selection. *S. cerevisiae* BJ5464-NpgA was transformed with corresponding plasmid combinations and was grown on select nutrient drop-out SD plates at 28 °C for 3 days. Single colony was picked up and grown in 2 mL SD media with selective nutrients dropped out for 24 h. 1 mL of SD media was then transferred to 100 mL SD media with selective nutrients dropped out for 48 h. 100 mL of SD media was then transferred to 1 L YPD (2% dextrose). The culture was shaken at 250 rpm at 28 °C for 6 hrs, Cells were harvested by centrifugation and resuspended in cell lysis buffer [50 mM $K_2HPO_4$ (pH 7.5), 10 mM imidazole, 300 mM NaCl, 5% glycerol]. Lysis of Cells using the

French Press, and the cell lysate was cleared by centrifugation at 26,000 g for 60 min at 4 °C. The supernatant was incubated with $Ni^{2+}$-NTA resin for 30 min at 4 °C and then the slurry was loaded onto a gravity column. The resin was washed and eluted with increasing concentrations of imidazole (from 10 mM to 500 mM) in cell lysis buffer. The fractions were examined by SDS-PAGE gels. Pure fractions were concentrated by Amicon concentrators (Millipore), supplemented with 10% glycerol and stored at -80 °C. Protein concentration was determined by quantifying the protein concentration measuring the absorbance at 280 nm.

**1.7 Pathway reconstitution in Aspergillus nidulans LO7890.**

To construct overexpression plasmids for *A. nidulans*, each gene was amplified from the *M. robertsii* ARSEF 23 gDNA by PCR and the corresponding overlapping fragments were ligated into double-digested pYTU, pYTR or pYTP vectors through yeast homologous recombination. The identity of resulting plasmid was identified by colony-PCR and enzymatic digestion. Protoplasts preparation, genetic transformation and heterologous production in *A. nidulans* were performed based on the reported protocol.[4] Plasmids were confirmed by DNA sequencing and plasmids maps were shown below.

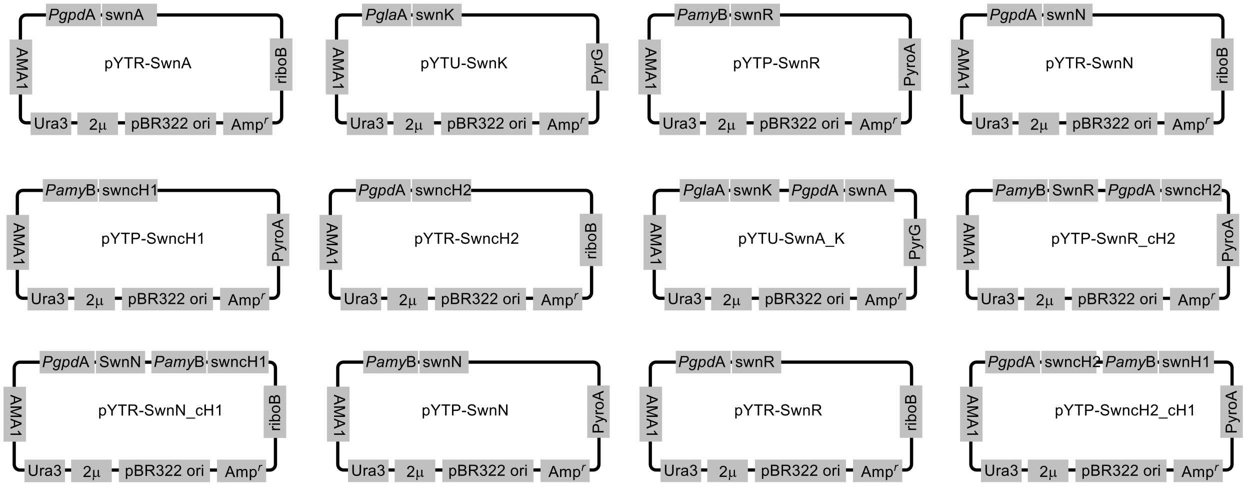

**1.8 LC-MS and GC-MS analysis of metabolites from small-scale fermentations.**

*A. nidulans* transformants were cultivated on 15 mL of CD-ST solid medium (10 g/L glucose, 50 mL/L 20× Nitrate Salts, 1 mL/L Trace Elements, 2% agar), supplemented with the necessary vitamins and precursors (either 2 mM L-PA, or 2 mM isotope-labeled lysine, or 0.5 mM of compound **1** or **2**) for 3 days at 28 °C. For LC-MS analysis, a 1 cm² section of the solid medium was extracted with 1 mL of acetone. The organic phase was dried under vacuum and the resulting residue was dissolved in methanol for LC-MS injection. To determine the absolute configuration of pipecolic acid generated by SwnA and reductase, the residue was dissolved in $CH_3CN$ and derivatized using Marfey's reagent. Briefly, 20 μL of Marfey's reagent [$N^{\alpha}$-(5-Fluoro-2,4-dinitrophenyl)-L-alaninamide] acetone solution (100 mM) was added to 20 μL of mixture, followed by the addition of 20 μL of 0.5M $NaHCO_3$ aqueous solution. The reaction mixture was incubated at 40 °C for 1 hr, and quenched with 50 μL of 1M HCl. The quenched reaction was subject to LC–MS analysis. For volatile compounds, a 0.5 $cm^2$ section of the solid medium was extracted with 1 mL of acetone. The organic phase was dried under vacuum, and the residue was dissolved in 400 μL of 1 M NaOH. The aqueous phase was then extracted with 1 mL of ethyl acetate. The organic layer was dried under vacuum, and the residue was treated with 100 μL of TMS-HT reagent (HMDS and TMSCl in anhydrous pyridine, obtained from TCI). After centrifugation, the supernatant was subject for GC-MS analysis.

**1.9 Isolation of natural products from either fermentation or in vitro enzymatic reactions.**

Isolation of **1** from fermentation: the corresponding *A. nidulans* transformants were cultured in liquid CD-ST medium (2% starch, 2% casamino acids, 50 mL/L nitrate salts, 1 mL/L trace elements solution), which was shaken at 28 °C for 4 days. After fermentation for 4 days, the cell culture was filtered by miracloth (GE Healthcare) to separate the medium and the mycelium. The pH of the medium was adjusted to 12 by NaOH. The Dichloromethane (DCM) extract from the supernatant and the acetone extract from the mycelium were combined and concentrated in vacuo to give an oily residue. The crude extract was dissolved in water and subjected to C18 (reversed-phase) flash chromatography and eluted with a linear gradient of 0-20% MeOH-$H_2O$ solvent system supplemented with 0.1% HCl (11.8 M) as additive to give compound **1** (456.7 mg, 114.2 mg/L).

Isolation of 8a-*d*-**1** from fermentation: the corresponding *A. nidulans* (coexpression of SwnK and SwnR) transformants were cultured in solid CD-ST medium [2% starch, 2% casamino acids, 50 mL/L nitrate salts, 1 mL/L trace elements solution, 2% agar, 2 mM 8a-*d*-*rac*-pipecolic acid (d, 80%)], which was shaken at 28 °C for 4 days. After fermentation for 4 days, the cell culture was extraced by acetone. The acetone extract from the cultured were combined and concentrated in vacuo to give an oily residue. The crude extract was dissolved in water and subjected to C18 (reversed-phase) flash chromatography and eluted with a linear gradient of 0-20% MeOH-$H_2O$ solvent system supplemented with 0.1% HCl (11.8 M) as additive to give 8a-*d*-**1** (70.0 mg, 70.0 mg/L).

Preparation of **1** from enzymatic reduction of **5**: A 5 mL scale reaction was performed at 30 °C in 100 mM KPi buffer (pH 6.5), containing 10 μM SwnN, 1 mM NADP, 10 μM glucose-6-

phosphate dehydrogenase (G6PD), 50 mM glucose-6-phosphate (G6P), and 5 mM **5**. After 4 h, the mixture reaction was directly subject to anion exchange chromatography using Dowex® 1x4-400 resin. The enzymatic product was eluted using $H_2O$. The corresponding water phase was subject to C18 (reversed-phase) flash chromatography and eluted with a linear gradient of 0-20% MeOH-$H_2O$ solvent system supplemented with 0.1% HCl (11.8 M) as additive to give compound **1** (3.4 mg, 80%).

Isolation of **2** from fermentation: the corresponding *A. nidulans* transformants were cultured in liquid CD-ST medium (2% starch, 2% casamino acids, 50 mL/L nitrate salts, 1ml/L trace elements), which was shaken at 28 °C for 4 days. After fermentation for 4 days, the cell culture was filtered by miracloth (GE Healthcare) to separate the supernatant and the mycelium. The supernatant was adjusted pH as 12 by NaOH. The Dichloromethane (DCM) extract from the supernatant and the acetone extract from the mycelium were combined and concentrated in vacuo to give an oily residue. The crude extract was dissolved in water and subjected to C18 (reversed-phase) flash chromatography and eluted with a linear gradient of 0-20% MeOH-$H_2O$ solvent system supplemented with 0.1% HCl (11.8 M) as additive to give compound **2** (18.2 mg, 4.6 mg/L).

Isolation of **3'** from fermentation: the corresponding *A. nidulans* transformants were cultured in solid CD-ST medium [2% starch, 2% casamino acids, 50 mL/L nitrate salts, 1 mL/L trace elements solution, 2% agar], which was shaken at 28 °C for 4 days. After fermentation for 4 days, the cell culture was extracted by acetone. The acetone extract from the cultured were combined and concentrated in vacuo to give an oily residue. The crude extract was dissolved in water and subjected to C18 (reversed-phase) flash chromatography and eluted with a linear gradient of 0-5% MeOH-$H_2O$ solvent system supplemented with 0.1% formic acid as additive to give **3'** (4.0 mg, 4.0 mg/L).

Preparation of **2** from enzymatic oxidation of **1**: A 20 mL scale reaction was performed at 30 °C in 100 mM KPi buffer (pH 6.5), containing 20 μM SwnH2, 2.5 mM ascorbate, 20 mM α-ketoglutarate, 100 μM $(NH_4)_2Fe(SO_4)_2$, 20 μM SwnN, 1 mM $NADP^+$, 10 μM G6PD, 50 mM G6P, and 5 mM **1**. After 4 hours incubation, the mixture reaction was directly subject to anion exchange chromatography using Dowex® 1x4-400 resin. The enzymatic product was eluted using $H_2O$. The corresponding water phase was subject to C18 (reversed-phase) flash chromatography and eluted with a linear gradient of 0-20% MeOH-$H_2O$ solvent system supplemented with 0.1% HCl (11.8 M) as additive to give compound **2** (13.6 mg, 70%).

Preparation of 8a-*d*-**2** from enzymatic oxidation of 8a-*d*-**1**: A 20 mL scale reaction was performed at 30 °C in 100 mM KPi buffer (pH 6.5), containing 20 μM SwnH2, 2.5 mM ascorbate,

20 mM α-ketoglutarate, 100 μM $(NH_4)_2Fe(SO_4)_2$ and 5 mM 8*a*-*d*-**1**. After 1 hours incubation, the mixture reaction was directly subject to anion exchange chromatography using Dowex® 1x4-400 resin. The enzymatic product was eluted using $H_2O$. The corresponding water phase was subject to C18 (reversed-phase) flash chromatography and eluted with a linear gradient of 0-20% MeOH-$H_2O$ solvent system supplemented with 0.1% HCl (11.8 M) as additive to give compound 8*a*-*d*-**2** (5.4 mg, 34%).

<u>Isolation of **6** from enzymatic oxidation of **1**</u>: A 20 mL scale reaction was performed at 30 °C in 100 mM KPi buffer (pH 6.5), containing 20 μM SwnH2, 2.5 mM ascorbate, 20 mM α-ketoglutarate, 100 μM $(NH_4)_2Fe(SO_4)_2$, and 5 mM **1**. After 2 hours incubation, the mixture reaction was directly subject to anion exchange chromatography using Dowex® 1x4-400 resin. The enzymatic product was eluted using $H_2O$. The corresponding water phase was subject to C18 (reversed-phase) flash chromatography and eluted with a linear gradient of 0-20% MeOH-$H_2O$ solvent system supplemented with 0.1% HCl (11.8 M) as additive to give compound **6** (7.5 mg, 39%).

<u>Isolation of **11** from in vitro enzymatic reaction with **5**</u>: A 20 mL scale reaction was performed in 100 mM KPi buffer (pH 6.5), containing 100 μM SwnH1, 2.5 mM ascorbate, 10 mM α-ketoglutarate, 200 μM $(NH_4)_2Fe(SO_4)_2$, 10 μM SwnN, 1 mM NADP, 10 μM G6PD, 20 mM G6P, and 2 mM **5**. The reaction is incubated at 30 ºC. After 4 hours incubation, the mixture reaction was directly subject to anion exchange chromatography using Dowex® 1x4-400 resin. The enzymatic product was eluted using $H_2O$. The corresponding water phase was subject to C18 (reversed-phase) flash chromatography and eluted with a linear gradient of 0-20% MeOH-$H_2O$ solvent system supplemented with 0.1% HCl (11.8 M) as additive to give compound **11** (2.5 mg, 32%).

<u>Enzymatic synthesis of Ac-**12** and **5**</u>: A 100 mL scale reaction was performed in 100 mM KPi buffer (pH 6.5), containing 20 μM SwnH2, 2.5 mM ascorbate, 20 mM α-ketoglutarate, 200 μM $(NH_4)_2Fe(SO_4)_2$, and 3 mM **Ac-1**. The reaction was incubated at 30 ºC. After 4 h, the reaction mixture was subject to anion exchange chromatography using Dowex® 1x4-400 resin. The enzymatic product was eluted using $H_2O$ and was subject to reversed-phase C18 flash chromatography and purified with a linear gradient of 0-20% MeOH-$H_2O$ solvent system supplemented with 0.1% HCl (11.8 M) as additive to give compound **5** (19.0 mg, 36%) and compound **Ac-12** (42.2 mg, 60%). To deprotect, 2 mL of saturated $NH_3•H_2O$ was added to **Ac-12** 36.3 mg, and the resulting mixture was stirred at 22 °C overnight. Solvent was removed give **12** quantitatively.

Enzymatic synthesis of **13**: 20 mL scale in vitro assay was performed at 30 °C in 100 mM KPi buffer (pH 6.5), containing 10 μM SwnH2, 2.5 mM ascorbate, 10 mM α-ketoglutarate, 200 μM $(NH_4)_2Fe(SO_4)_2$, 10 μM SwnN, 2 mM NADP, 10 μM G6PD, 20 mM G6P, and 2 mM **12**. The reaction is incubated at 30 ºC. After 4 h, the mixture reaction was directly subject to anion exchange chromatography using Dowex® 1x4-400 resin. The enzymatic product was eluted using $H_2O$. The corresponding water phase was subject to C18 (reversed-phase) flash chromatography and eluted with a linear gradient of 0-20% MeOH-$H_2O$ solvent system supplemented with 0.1% HCl (11.8 M) as additive to give compound **13** (4.6 mg, 55%).

Enzymatic synthesis of SW from **1**: A 10 mL scale reaction was performed at 30 °C in 100 mM KPi buffer (pH 6.5), containing 20 μM SwnH2, 20 μM SwnH1, 2.5 mM ascorbate, 20 mM α-ketoglutarate, 100 μM $(NH_4)_2Fe(SO_4)_2$, 20 μM SwnN, 10 mM NADP, 1 mg/mL GDH, 100 mM D-glucose, and 5 mM **1**. After 4 hours incubation, the mixture reaction was directly subject to anion exchange chromatography using Dowex® 1x4-400 resin. The enzymatic product was eluted using $H_2O$. The corresponding water phase was subject to C18 (reversed-phase) flash chromatography and eluted with a linear gradient of 0-20% MeOH-$H_2O$ solvent system supplemented with 0.1% HCl (11.8 M) as additive to give compound SW (5.7 mg, 66%).

Enzymatic synthesis of SW from **6**: A 10 mL scale reaction was performed at 30 °C in 100 mM KPi buffer (pH 6.5), containing 20 μM SwnH1, 2.5 mM ascorbate, 20 mM α-ketoglutarate, 100 μM $(NH_4)_2Fe(SO_4)_2$, 20 μM SwnN, 10 mM NADPH, and 2 mM **6**. After 4 hours incubation, the mixture reaction was directly subject to anion exchange chromatography using Dowex® 1x4-400 resin. The enzymatic product was eluted using $H_2O$. The corresponding water phase was subject to C18 (reversed-phase) flash chromatography and eluted with a linear gradient of 0-20% MeOH-$H_2O$ solvent system supplemented with 0.1% HCl (11.8 M) as additive to give compound SW (2.5 mg, 72%).

Enzymatic synthesis of SW from **2**: A 10 mL scale reaction was performed at 30 °C in 100 mM KPi buffer (pH 6.5), containing 20 μM SwnH1, 2.5 mM ascorbate, 20 mM α-ketoglutarate, 100 μM $(NH_4)_2Fe(SO_4)_2$, 20 μM SwnN, 10 mM NADPH, and 2 mM **2**. After 4 hours incubation, the mixture reaction was directly subject to anion exchange chromatography using Dowex® 1x4-400 resin. The enzymatic product was eluted using $H_2O$. The corresponding water phase was subject to C18 (reversed-phase) flash chromatography and eluted with a linear gradient of 0-20% MeOH-$H_2O$ solvent system supplemented with 0.1% HCl (11.8 M) as additive to give compound SW (2.7 mg, 78%).

**1.10 Chemical synthesis of substrate and standard.**

Synthesis of Ac-**1**: To a stirred solution of **1** (125.5 mg, 1 mmol) in 6 mL pyridine at 22 °C, $Ac_2O$ (4 mL, 1 mmol) and DMAP (10.0 mg, 0.08 mmol) were sequentially added. The resulting mixture was stirred at 22 °C for 4 h. 1 mL MeOH was used to quench this reaction. Solvent was removed under in vacuo and the residue was subject to gel filtration chromatography by using Sephadex LH20 resin with methanol as the mobile phase to quantitatively give **Ac-1** (165.6 mg, 90%).

Synthesis of **9**: To a stirring solution of oxalyl chloride (110 μL, 1.3 mmol) in $CH_2Cl_2$ (4 mL), cooled to -78 °C, DMSO (185 μL, 2.6 mmol) was added dropwise. After 30 min, a solution of **1** (20 mg, 0.14 mmol) in $CH_2Cl_2$ (1 mL) was added. After an additional 30 min, $Et_3N$ (1 mL, 7.2 mmol) was added, and the mixture was allowed to stir at room temperature. The mixture was quenched with saturated aq. $NaHCO_3$ (25 mL), and extracted with $CH_2Cl_2$ (3 × 25 mL). The combined organic layers were washed with brine (50 mL), dried over anhydrous $MgSO_4$, and filtered. The solvent was removed under vacuum to give compound **9** (8.9 mg, 45%) as a yellow oil.

Synthesis of radical clock substrate **2'**: To a stirring solution of *cis*-pyrrolidine-3,4-diol hydrochloride (60.0 mg, 0.43 mmol) in MeOH (10 mL) were added acetic acid (0.2 mL,3.3 mmol), sodium cyanoborohydride (81.5 mg, 1.29 mmol, 3.0 equiv), and (1-ethoxycyclopropoxy)trimethylsilane (0.2 mL, 1.0 mmol). The reaction mixture was stirred at 60 °C for 24 h. Completion was confirmed by TLC analysis ($CH_2Cl_2$/MeOH/$NH_4OH$ 10:10:1). The mixture was concentrated in vacuo, the residue dissolved in water and subject to C18 (reversed-phase) flash chromatography and eluted with a linear gradient of 0-10% MeOH-$H_2O$ solvent system supplemented with 0.1% HCl (11.8 M) as additive to give compound **2'** (33.0 mg, 43% yield) as a pale-yellow oil.

**1.11 Biochemical characterization of swnA**

Each reaction was performed at 100 µL scale in 100 mM KPi buffer (pH 8.0), containing 10 µM SwnA, 10 mM α-ketoglutarate, and 1 mM L-Lysine ($^{15}N$ labeling). To analyse the product by LC–MS, the quenched reaction mixture was derivatized by Fmoc-Cl. Briefly, 15 µl of reaction mixture was mixed with 15 µl of sodium borate solution (1 M), and 15 µl of Fmoc-Cl stock solution (in acetonitrile) was added. The resulting mixture was incubated at 37 °C for 30 min and then injected into the LC–MS system. For coupled reaction, 10 µM reductase (SwnR or SwnN) and 5 mM NADPH were included. The reaction was quenched with 1 equal volume of methanol after 2 hours incubation at 30 °C and centrifuged at 14,000 g for 5 min to remove protein pellet.

For non-coupled SwnA reaction, $NaBH_3CN$ aqueous solution (50 mM final concentration) was added and the resulting mixture was incubated at 37°C for 30 mins to chemically reduce the dehydropipecolic acid product. The enantiopurity of the reduced pipecolic acid products were derivatized by Marfey's reagents. Briefly, 20 μL of Marfey's reagent [$N^{\alpha}$-(5-Fluoro-2,4-dinitrophenyl)-L-alaninamide] acetone solution (100 mM) was added to 20 μL of reaction mixture, followed by the addition of 20 μL of 0.5M $NaHCO_3$ aqueous solution. The reaction mixture was incubated at 40 °C for 1 hr, and quenched with 50 μL of 1M HCl. The quenched reaction was subject to LC–MS analysis.

**1.12 Biochemical characterization of SwnK**

A 50 μL scale reaction was performed at 22 °C in 100 mM KPi buffer (pH 7.0), containing 20 μM SwnK, 10 μM reductase (SwnR or SwnN), 5mM Malonyl-CoA, 10 mM $MgCl_2$, 10 mM NADPH, 5 mM ATP, and 1 mM L-pipecolic acid or deuterium labeled (d9)-D/L-pipecolic acid. The reaction was quenched with 4 equal volumes of acetonitrile after 2 hours incubation at 22 °C and centrifuged at 14,000 g for 5 min. The reaction product was analyzed by LC–MS.

**1.13 Biochemical characterization of tailoring enzymes**

All analytical tests were performed at 100 μL scale in assay buffer [100 mM KPi buffer (pH 6.5)] containing 10 μM enzyme (SwnH2 or SwnH1) and the necessary cofactors [2.5 mM ascorbate, 100 μM $(NH_4)_2Fe(SO_4)_2$], 10 mM α-ketoglutarate, and 2 mM substrates. Each reaction was quenched with 1 equal volumes of methanol after 10 min to 2 hours incubation at 30 °C and centrifuged at 14,000 g for 5 min before LC–MS analysis. When SwnN and SwnR were included in the assays, 10 μM enzyme and 10 mM NADPH were used.

**1.14 Determining the ratio between 1 and 5 formed in SwnH2-catalyzed oxidation of 1.**

A 2 mL scale reaction was performed at 30 °C in 100 mM KPi buffer (pH 6.5), containing 5 μM SwnH2, 2.5 mM ascorbate, 5 mM α-ketoglutarate, 100 μM $(NH_4)_2Fe(SO_4)_2$, and 5 mM **1**. The mixture reaction was quenched with 1 equal volumes of methanol after 30 min incubation at 30 °C and centrifuged at 14,000 g for 5 min before analyzed by LC–MS. The LC-MS recorded the ratio of ions with m/z 142 and 143. Subsequently, 50 μL of 1 M $NaCD_3CN$ was added to the reaction mixture, which was then incubated at 37 °C for 30 minutes. After a second centrifugation at 14,000 g for 5 minutes, LC-MS analysis was again performed to record the ratio of ions with m/z 142 and 143. Comparison of the data before and after the addition of NaCD3CN allowed the measurement of the recovery ratio of the starting material **1** and the formation of compound **5**. Following this, all solvents were removed, and the residue was subjected to gel filtration chromatography using Sephadex LH20 resin with methanol as the mobile phase to quantitatively isolate the mixture of **1** (including both nondeuterated leftover **1** and deuterated **1** resulted from the reduction of **5** and **2** for NMR analysis to determine the relative ratio.

**1.15 Determining the kinetic isotope effect of SwnH2 and SwnH1-catalyzed oxidation of 2.**

A 1:1 mixture of 8a-deuterated and non-deuterated **2** was prepared as 8a-*d*-**2** (*d*, 50%). A 100 uL analytical reaction was performed at 30 °C in 100 mM KPi buffer (pH 6.5), containing 10 μM enzyme (SwnH2 or SwnH1), 2.5 mM ascorbate, 10 mM α-ketoglutarate, 100 μM $(NH_4)_2Fe(SO_4)_2$, and 2 mM 8a-*d*-**2** (*d*, 50%). Each reaction was quenched with 1 equal volumes of methanol after 10 min to 2 hours incubation at 30 °C and centrifuged at 14,000 g for 5 min before LC–MS analysis. In order to confirm the mechanism of SwnH1, 10 μM SwnN and 10 mM NADPH were used in a cascade assay of SwnH1 and SwnN.

**1.16 Assay for SwnH1-catalyzed oxidation of the radical clock substrate 2'.**

A 100 uL analytical reaction was performed at 30 °C in 100 mM KPi buffer (pH 6.5), containing 40 or 80 μM enzyme (SwnH2 or SwnH1), 10 mM ascorbate, 20 mM α-ketoglutarate, 100 μM $(NH_4)_2Fe(SO_4)_2$, and 5 mM **2'**. Reactions were incubated for 20 minutes at 30 °C and quenched by the addition of 100 μL methanol. The resulting mixtures were centrifuged at 14,000 × g for 5 minutes to remove precipitated proteins. To derivatize the aldehyde product generated via radical clock ring opening, 50 μL of the supernatant was mixed with 50 μL of a 50 mM phenylhydrazine solution prepared in a 4:1 (v/v) mixture of ethanol and 1 M sodium acetate–acetic acid buffer (pH 5.5). The mixture was incubated at 37 °C for 1 hour, then centrifuged at 14,000 × g for 5 minutes before LC–MS analysis.

**1.17 Determination of the absolute configuration of 1 by Mosher's Ester Method.**

**1** was dissolved in anhydrous pyridine to prepare a 100 mM stock solution. An aliquot of 20 μL of this stock solution was reacted with 2 μL of either (R)-MTPA-Cl or (S)-MTPA-Cl in a total volume of 50 μL dry pyridine. The reactions were carried out at 37 °C for 1 hour. Following the reaction, solvents were evaporated under reduced pressure. The resulting Mosher esters were redissolved in deuterated pyridine ($C_5D_5N$) for NMR analysis.

#### 2. Supplementary Table

**Table S1. Primers used in this study.**

| Primers | Sequences ( 5' → 3') |
|---|---|
| xw55_SwnK_For | TGGCTAGCCATCACCATCACCATCACCATCACACTAGTGGCATCTCAGATCACGAC |
| xw55_SwnK_Rev1 | CTGCTAGCTTCCACTGATGCCATCTTGCCC |
| xw55_SwnK_For1 | TGCCGGCTTGTCATGATGCGCGGTC |
| xw55_SwnK_Rev | TCGTGAAGGCATCGGTCCGCACAAATTTGTCATTTTCACGTACGCATTTTGGCCC |
| xw06_SwnN_For | TATCAACTATTAACTATATCGTAATACCATATGGTCGTCGTTGCCGTTGC |
| xw06_SwnN_rev | ATAAATCGTGAAGGCATGTTTAAACCTAGGCTACGGTGCTCTGCTCTGCT |
| xw02_SwnR_For | ATCAACTATTAACTATATCGTAATACCATATGCGTGTGGCAATTGCT |
| xw02_SwnR_Rev | ACTATAAATCGTGAAGGCATGTTTCTACAGTATCGAGTCCGGATGC |
| pYTU_SwnK_For | TCCCCAGCATCATTACACCTCAGCAATGAGTGGCATCTCAGATCACG |
| pYTU_SwnK_Rev | TTCAACACAGTGGAGGACATACCCGTAATTTTCTGTTCGAAGTTTGCGAGACGGAC |
| pYTP_SwnR_For | TGAACAATAAACCCCACAGAAGGCATTTTTAATTAATGCGTGTGGCAATTGCTGG |
| pYTP_SwnR_Rev | GCTTGATATCGAATTCCTGCAGCCCGGGGGATCCTGGCAAGGGCCAAAATGCGTA |
| pYTR_SwnN_For | TACCCCGCCACATAGACACATCTAAACATTAATTAATGGTCGTCGTTGCCGTTGC |
| pYTR_SwnN_Rev | AGGGTATCATCGAAAGGGAGTCATCCAATTTAAATTGCTCTGGACCATCTTTATAGCATC CAC |
| pYTP_SwnH1_For | TGAACAATAAACCCCACAGAAGGCATTTTTAATTAATGGGCGCCTTTCCTCCCCC |
| pYTP_SwnH1_Rev | GCTTGATATCGAATTCCTGCAGCCCGGGGGATCCTCGCGGGACCAGGACCGG |
| pYTR_SwnH2_For | TACCCCGCCACATAGACACATCTAAACATTAATTAATGATCAATTCAGACGCACAGTCGG |
| pYTR_SwnH2_Rev | AGGGTATCATCGAAAGGGAGTCATCCAATTTAAATATGATTTTGGCAGCCTGTTAGTCTT |
| pYTR_SwnA_For | TACCCCGCCACATAGACACATCTAAACATTAATTAATGCACTTGGAGCGGGAC |
| pYTR_SwnA_Re | AGGGTATCATCGAAAGGGAGTCATCCAATTTAAATTGGCGCTGGGCATAACTC |
| pet_SWnH1_For | TGTACTTCCAATCCAATGGCGCCTTTCCTCCCCCC |
| pet_SWnH1_Rev1 | GCTCCATCTCCGCGAAGCAAACCTCGTC |
| pet_SWnH1_For1 | TGCTTCGCGGAGATGGAGCCGGGCTCGG |
| pet_SWnH1_Rev | GTTATCCACTTCCAATTCACGAGTTAGCCATGCCCA |
| pet_SWnH2_For | TGTACTTCCAATCCAATATCAATTCAGACGCACAGTC |
| pet_SWnH2_Rev1 | GCGCTCCATTTCAAGAAAGGAAACTTCATCGGGCC |
| pet_SWnH2_For1 | CTTTCTTGAAATGGAGCGCGGGTCTG |
| pet_SWnH2_Rev | CCGTTATCCACTTCCAATTTACTTGAGCATCTTCTCATAAATAGCC |
| pet_SwnN_For | TGTACTTCCAATCCAATATGGTCGTCGTTGCCGTTG |
| pet_swnN_Re | TTATCCACTTCCAATCTACGGTGCTCTGCTCTGCT |
| pet_SwnR_For | TGTACTTCCAATCCAATATGCGTGTGGCAATTGCTGG |
| pet_SwnR_Re | TTATCCACTTCCAATCTACAGTATCGAGTCCGGATGC |

**Table S2. Amino acid sequence of characterized proteins in this study.**

| Name | NCBI Accession number | Sequence |
|---|---|---|
| SwnA | XP_007824806.2 | MHLERDKVYDAPEGEVWSTVKPASTHNSAAPKRLAQRWN<br>HRWSDESLTQGVSPLKDSSKTVKASTTIPLGTGRPTALYYP<br>WQSVSMAGTEASRQPRGLKPTLAGNMTCSKGEAAFDLSS<br>ALNYGEPSGWSQLVAFFRETTGRVHRPPYADWDTTLTCG<br>STSAVDLVLRMFCNRGDCVLAERFTYPGTLMASRAQGLRT<br>VGIAMDADGLVPEALDAALRGWDAASRGRKPFVLYTIPSG<br>HNPTGVTQSAARKRAVYQVAERHDLLIVEDDPYFFLRLGR<br>GGGGESLPTSYLSLDTAGRVVRVESTSKILAPGLRCGWLT<br>ASRQVVDMFGNFAEVGPSSPAGPSQAMLYKLLVESWGPE<br>GFAGWLDYLSGEYGRRRDVMVAACERHLPREVCAWTAPT<br>HGMFLWVGAALERHPRYQESGARDQDRDADADAELCRA<br>VEDGICARAEANGVLVARGSWCRVGGGADRAFFRMTFVA<br>TAEAADLERGVAQFGRAVRDEFGLG |
| SwnK | XP_007824811.1 | MSGISDHDYQVLVNEFNATEDISLLGSRLDQLFEQVADRFP<br>DNTAVIHNESEVTFKQLNSSANILARCLAKRGLQQGDVVGL<br>AVSRSIDLIAAMLAVLKLGAAYVPIDPSFPAERINQMVEDAG<br>LRLILLSGRPTKGLGRWASLCLSVSEARDGSVTDGTNLETE<br>IQTRDLAYVIYTSGSTGRPKGVEISHGAAANFLSSLRKHEP<br>GCSERDRLLAITTISFDMSALELLLPLVSGSAMIVANTSAVK<br>DPRELISLMARHKVTILQATPATWTMLLESGWKGNPRLSKV<br>ICGGEPLSRQLADRLLAAADSVWNVYGPSETTYGSVGRVG<br>QGDIVVGNPVANGRIYVLDDNMSPVPIGSEGEVYIGGGSVS<br>NGYRNKAELTRSRFLVNPFHGGVFFRTGDLARFIAPGKLQ<br>VVGRIDGVVKIRGHRIDVGDIEAVLVDHANVSEAVVVSRDD<br>RLVAYCVLHAPCHDAASLDGILRPWVAERLPAYMLPTFFV<br>QMDALPLSPSEKVNRKALPDPIEAMQHQTMIQPTSELEQRI<br>QAIWSDILGHDRFGIEDNFFRIGGDSVRIIRMQAALEKQLHR<br>PVPTPKLFEHYTIKALAAYLAGTGRENNNESQAVSQTFAGS<br>HEDIAIVSMACRLPGGVATPEALWQLLQSGGDTIIDVPKDR<br>WDADKLYNADANIDGTSYCRRGGFLDAIYSYDASFFGISPR<br>EAQAMDPTHNLMLELCWEGFERSGYTRDQLSGSATGVFL<br>GVSNNATTNGTPPDLKGYSITGSASATMCGRLSHTLGLRG<br>PSLAVDTACSSSLVATHLACNALRQGECSMALAGGVSLLT<br>TPGIHIEFSKLGGLSADGRCRAFSDDTDGTGFSEGAAIIVLK<br>RLSDARRDGDDIHAVLRGTAVMHGGSSAGLTAPSGPGQV<br>ALLRNALARAALEPGDIDYVEAHGTATKLGDPIEATALAEVF<br>GTERSGSDPLRIGSAKSNLGHTQAAAGVVGLLKVVLSMNH<br>DTLPRTLHVREPMAAVDWKRTNMELVLQNRPWLPNNNRL<br>RRAGVSAFGIGGTNAHVIVEESPPPAVEETGNITLPSLPATL<br>PFVLSGQGDSALRAQAEKLRLHIESGAGKDSPLRDVAYSL<br>ATCRSHLHRRLVVMAGDKAETLEKLASVSSGPTQPLSVNE<br>VGSPTVAMLFTGQGSQLPGMGKDLYAVYPVFRDALDEIAA<br>KFTDLERPLLDIMWAESGSENAALLSRTDFAQPALFALEVS<br>LWKLWQSWGVKPDFLLGHSVGELAAAHAAGVLDLSDACR<br>LVMMRGRLMQAIPRQGKMASVEASSAEVSAAIQELGQHD<br>KVEIAGYNTPLQTVISGHEEAVEATSVYMSKLGRKTKLLDT<br>SHAFHSFHMNGMLDDLRALAQNIRFSPPKMRIISSMTGRLA<br>GAGELERPEYWAQQARNAVRFSDAFQTLAGQGANVFLEL<br>GPSAMLCGLGAACLADPGQVGAALWLPSLKPNMNGPLVI<br>QSSLSELHVRHVPVDWAAYFKPFDCKRVMLPTYAFQREDF<br>RPANKASWFDVASLSTGTNDAAPRVQDMMFEINWRRVET<br>KSIQPRGAWGLLCPSGETAWTTEAQRALLATGIQLVAVSK<br>PHEADQLDGLLSLWDSDADTVQMAHGVTALALAQLQEAIR<br>TGLGAPIVWVTRHAVGAGADDRPVNIGAGPLWGLMRAAR |

| | | |
|---|---|---|
| | | SEHPELRLRLIDVDEETDRASLSQAIMLADQTEIAVRREQLL MPHMERAGLAAPLPVGQPFVRTDGAVLVTGGLGDLGSRV ARRLATAHGVCDLVLVSRQGTNSPGADALVAELAELGAKA TIVGCDLADLDSLGSVMQLFTPDRPLRGVVHAAGVVDSGT LSSLTPRKCAAVFAPKVDGLWNLHQLTKHMDLDLFMMFSS ISGIIGLPGLGNYAAANSFIDSLAHLRRAQGLPASSVAYGTW AGDGMATTLVPTTRAHLSQLGLGFLPPEAGLEIFEHAVYQG RALTVAAVLDLQRLRAYYEEQGGVPPLLNSMLGGTKVQKP ADEVVNLRDLLADAAPEQHSSIVLHMVQATIAKALGYTRAE DVDASRPMQELGIDSLTAILTRNHLATLTGMALPPNIALLYP NLKSLSEFLLCRLMDDVESSTSSPSDTDGATPSTPTSAASC VDMADIRRGVLDSTFQFNNPVSPGVPGTVFVTGATGFVGT FMIHEFLQRRISVYCLVRASGSQEAQQRMITTLKQYGLWR PEHEPLLVSVAGDLSQPLLGLGEVVFDDLASRVDAILHSGA LVDWMRPLDDYIGPNVLGTHEVLRLASCGRAKAIHFISTIST LPIHAGYGLAEHDGEYGYGTSKYLAEKMVVAARFRGAKAS SYRLPFVAASAASGRFRLDRGDFLNNLVTGSLDLGAFPLIN ADLSSVLPVDYLCGTIAAIMTEDQERVGEDYDFVNPQALTF KHFFEMMCSVSGGKEMVSFGEWHRRALEYAAAHPTSSLA RIATVLDGYNDKTVGSLVSGSPVGKHVLGLDAYRAPPLDE EYVRRYVHCIEAARAKMRT |
| SwnR | XP_007824810.1 | MRVAIAGYGDLTRYICEEFVKAGHVLVILTRSYKPQLESQG VAQAITDYSPSSLRAPLADCEVLISTISDISSAYTNVHRNLIL ACQESPRCKRFIPAEFVADIEAYPDEPGFYYAPHEPIREML RGQTDLEWTLVCIGWLSDYFVPSKNRHIKDIGEFHPMNWA GNKIVIPGTGNEPVDFTWARDVVRGLASLIEAPRGSWEPYT FMSGERSCWNDATRLAVQKYRPGIPIQHVSLHSVAGMIKT AKDENTEVLADYYLLSISQACAIPTDKVEAHREKYFSGVTF RSLRDGLCQVDEHPDSIL |
| SwnN | EXU97980.1 | MVVVAVAGGTGGVGRTVLDAIAKSGQHQAIVLSRTTSVPT AVDEPKRFAVDYNSVEQMKKILQENNVQVVVSALLLVDEA VAQSQINLIRAAAQSGTVTKFIPSEYYIDFHAPIPGADLFTNF QLEAEAELSRHAQLTWTLIRVGIFLDHLTMPHNPKTTYITPF WVFVDIDHEQCVFPGDGSQPLVLTHSQDLAAYIERLVGLPA ENWPRESVVASNKLLVKDLESLVNKVTGKKFKVAYDSVECI HKGRITQLPSNTAVFQDPAKGEMFRDVEHQVMLSMLSRA HDLPGKNLAELFPEVETTDIEDFFRSGWTLKQSRAP |
| SwnH1 | XP_007824807.1 | MGAFPPPIDPSYIPSVSLTRLPATAPTADIIATLERDGALILV DLVSPQDVAAINAEIEPYIQKARAESHEAYDLIPKQTIMVPG VVGKSPTMARMAELDVIDTLRTRVLQRKCTATWEDRTEDF SIDPLLNSSLTYHISYGGPRQRLHRDDMIHGIYHRGGEYRL SDETMLGFMIAGSKTTRENGATMAIPGSHKWDHARVPRVD EVCFAEMEPGSALVFLGTVYHGAGHNSVPDQVRKIYGLFFI PGTLRPEENQFLAIPRSKVLGMSDKMLSLLGYKKPGTWLGI VNNGDPAENLAEVLGMANS |
| SwnH2 | XP_007824812.1 | MINSDAQSAQKQVEVEKPDEKYSAPRLLPPIPDSYQPAKAI TKIPATSSLEDILAILERDGGVILTDFVSLQELDKIDEELEPYT KSSIADDDSYNNFIGKKTLVIPGLVGKSDTIANILDTNETIDKL LKVILEERYPAVFEQHTEELVVDPLLSICMGFHVGHGSPRQ ALHRDDMIFSSKHRPDMKINEVDGFSCFLAGTRITRENGGT MVILGSHKWEHDRRGRPDEVSFLEMERGSAFIFLSTLAHG AGYNTIPGEVRKITNLVFCRGTLRTEENQFLCVPRSKVLKM SPKMQTLLGFKKPAGSWLGMVENEDPAKDLEAIYEKMLK |

**Table S3. Structural characterization of 1.**

<table>
<tr><td></td><td colspan="2">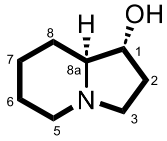

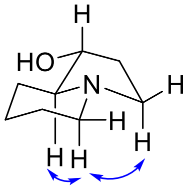

Key NOESY signals are shown<br>$J_{1,8a}$ = 7.7 Hz, the *trans* configuration between H-1 and H-8a<br>$[\alpha]_D^{20}$ = -33.2 ($c$ = 0.5 in ethanol), literature report: $[\alpha]_D^{27}$ = -51.0 ($c$ = 0.7 in ethanol)[5,6]<br>HRMS m/z calculated for $C_8H_{16}NO^+$ $[M+H]^+$ 142.1232, found 142.1227.</td></tr>
<tr><td>Pos.</td><td>$\delta_C$ ($CD_3OD$)</td><td>$\delta_H$ ($CD_3OD$), multi, $J$, integration</td></tr>
<tr><td>1</td><td>76.3, CH</td><td>3.82, ddd, $J$ = 9.2, 7.7, 4.8 Hz, 1H</td></tr>
<tr><td>2</td><td>32.0, $CH_2$</td><td>$H_a$: 2.23-2.12, m, 1H<br>$H_b$: 1.58-1.45, m, 1H</td></tr>
<tr><td>3</td><td>53.4, $CH_2$</td><td>$H_a$: 2.94, td, $J$ = 9.0, 2.3 Hz, 1H<br>$H_b$: 2.36, q, $J$ = 9.2 Hz, 1H</td></tr>
<tr><td>4</td><td>n.a.</td><td>n.a.</td></tr>
<tr><td>5</td><td>54.4, $CH_2$</td><td>$H_a$: 3.01, ddd, $J$ = 11.0, 3.7, 3.7 Hz, 1H<br>$H_b$: 2.12-2.04, m, 1H</td></tr>
<tr><td>6</td><td>25.8, $CH_2$</td><td>$H_a$: 1.65, dddd, $J$ = 11.3, 4.2, 2.0, 2.0 Hz, 1H<br>$H_b$: 1.58-1.45, m, 1H</td></tr>
<tr><td>7</td><td>25.0 $CH_2$</td><td>$H_a$: 1.83, ddd, $J$ = 12.9, 3.3, 3.3 Hz, 1H<br>$H_b$: 1.30, dddd, $J$ = 12.8, 12.8, 3.8, 3.8 Hz, 1H</td></tr>
<tr><td>8</td><td>29.3, $CH_2$</td><td>$H_a$: 2.00, ddd, $J$ = 12.2, 5.2, 2.8 Hz, 1H<br>$H_b$: 1.26-1.16, m, 1H</td></tr>
<tr><td>8a</td><td>72.1, CH</td><td>1.78, ddd, $J$ = 10.5, 7.8, 2.7 Hz, 1H</td></tr>
</table>

**Table S4. Structural characterization of 1•HCl.**

| | 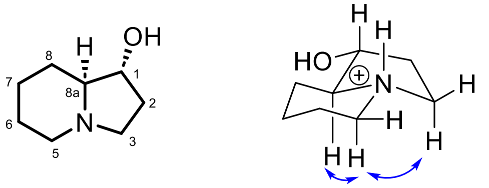 Key NOESY signals are shown | | 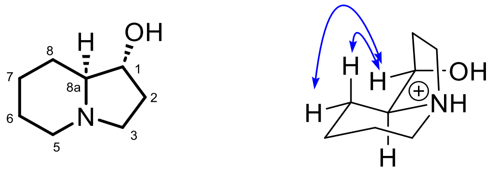 Key NOESY signals are shown | |
|---|---|---|---|---|
| Pos. | $\delta_C$ ($CD_3OD$) | $\delta_H$ ($CD_3OD$), multi, *J*, integration | $\delta_C$ ($CD_3OD$) | $\delta_H$ ($CD_3OD$), multi, *J*, integration |
| 1 | 73.1, CH | 4.13, ddd, J = 9.0, 9.0, 5.8 Hz, 1H) | 74.3 CH | 4.29-4.24, m, 1H |
| 2 | 30.3, $CH_2$ | $H_a$: 2.45-2.33, m, 1H<br>$H_b$: 1.89-1.70, m, 1H | 31.5, $CH_2$ | $H_a$: 2.59-2.48, m, 1H<br>$H_b$: 2.06-1.89, m, 1H |
| 3 | 52.1, $CH_2$ | $H_a$: 3.63-3.51, m, 1H<br>$H_b$: 3.28-3.17, m, 1H | 48.6, $CH_2$ | $H_a$: 3.63-3.51, m, 1H<br>$H_b$: 3.51-3.40, m, 1H |
| 4 | n.a. | n.a. | n.a. | n.a. |
| 5 | 53.4, $CH_2$ | $H_a$: 3.63-3.51, m, 1H<br>$H_b$: 3.04, ddd, J = 12.6, 12.6, 3.2 Hz, 1H | 48.3, $CH_2$ | 3.34-3.28, m, 2H |
| 6 | 24.0, $CH_2$ | $H_a$: 2.06-1.89, m, 1H<br>$H_b$: 1.89-1.70, m, 1H | 19.5, $CH_2$ | 1.89-1.70, m, 2H |
| 7 | 23.1 $CH_2$ | $H_a$: 2.06-1.89, m, 1H<br>$H_b$: 1.69-1.49, m, 1H | 21.0 $CH_2$ | $H_a$: 1.89-1.70, m, 1H<br>$H_b$: 1.69-1.49, m, 1H |
| 8 | 27.0, $CH_2$ | $H_a$: 2.06-1.89, m, 1H<br>$H_b$: 1.69-1.49, m, 1H | 23.3, $CH_2$ | $H_a$: 1.89-1.70, m, 1H<br>$H_b$: 1.49-1.41, m, 1H |
| 8a | 71.4, CH | 2.99-2.88, m, 1H | 48.6, CH | 3.63-3.51, m, 1H |

**Table S5. Structural characterization of 2.**

| | | | |
|---|---|---|---|
| | 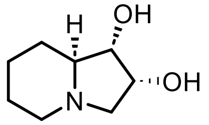<br>$J_{1,8a}$ = 9.0 Hz, the *trans* configuration between H-1 and H-8a<br>$J_{1,2}$ = 7.0 Hz, the *cis* configuration between H-1 and H-2<br>$[\alpha]^{20}_{D}$ = -21.0 (*c* = 0.2 in $H_2O$), literature report: $[\alpha]^{20}_{D}$ = -33.6 (*c* = 0.25 in $H_2O$)[6]<br>HRMS m/z calculated for $C_8H_{16}NO_2^+$ $[M+H]^+$ 158.1181, found 158.1176. | | |
| Pos. | $\delta_C$ ($CD_3OD$) | $\delta_H$ ($CD_3OD$), multi, *J*, integration | $\delta_H$ ($D_2O$), multi, *J*, integration |
| 1 | 76.6, CH | 3.52, dd, *J* = 8.7, 7.1 Hz, 1H | 3.54, dd, J = 9.0, 7.0 Hz, 1H |
| 2 | 28.2, CH | 4.10, ddd, *J* = 6.9, 6.9, 5.2 Hz, 1H | 4.12. ddd, J = 7.0, 7.0, 5.2 Hz, 1H |
| 3 | 58.2, $CH_2$ | $H_a$: 3.40, dd, *J* = 10.2, 6.9 Hz, 1H<br>$H_b$: 2.13, dd, *J* = 10.2, 5.3 Hz, 1H | $H_a$: 3.36, dd, *J* = 10.7, 6.9 Hz, 1H<br>$H_b$: 2.11, dd, *J* = 10.7, 5.2 Hz, 1H |
| 4 | n.a. | n.a. | n.a. |
| 5 | 48.5, $CH_2$ | $H_a$: 2.98, ddd, *J* = 10.8, 3.2, 3.2 Hz, 1H<br>$H_b$: 2.07, ddd, *J* = 11.6, 11.6, 3.0 Hz, 1H | $H_a$: 2.92, ddd, J = 11.3, 3.4, 3.4 Hz, 1H<br>$H_b$: 2.06, ddd, J = 11.8, 11.8, 2.9 Hz, 1H |
| 6 | 19.8, $CH_2$ | $H_a$: 1.70-1.60, m, 1H<br>$H_b$: 1.57-1.45, m, 1H | $H_a$: 1.64-1.58, m, 1H<br>$H_b$: 1.45-1.32, m, 1H |
| 7 | 15.8, $CH_2$ | $H_a$: 1.87-1.79, m, 1H<br>$H_b$: 1.37-1.25, m, 1H | $H_a$: 1.81-1.73, m, 1H<br>$H_b$: 1.31-1.19, m, 1H |
| 8 | 24.7, $CH_2$ | $H_a$: 2.02-1.95, m, 1H<br>$H_b$: 1.25-1.15, m, 1H | $H_a$: 1.92, dd, *J* = 13.0, 3.1 Hz, 1H<br>$H_b$: 1.18-1.07, m, 1H |
| 8a | 189.6, C | 1.92, ddd, *J* = 11.1, 8.6, 2.8 Hz, 1H | 1.98, ddd, *J* = 11.4, 9.1, 2.8 Hz, 1H |

**Table S6. Structural characterization of 5•HCl.**

| | 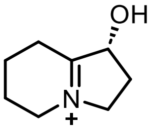 HRMS m/z calculated for $C_8H_{14}NO^+$ $[M+H]^+$ 140.1075, found 140.1071. | |
|---|---|---|
| Pos. | $\delta_C$ ($D_2O$) | $\delta_H$ ($D_2O$), multi, *J*, integration |
| 1 | 76.6, CH | 5.14-5.06, m, 1H |
| 2 | 28.2, $CH_2$ | $H_a$: 2.63, dddd, *J* = 13.6, 8.2, 8.2, 2.9 Hz, 1H<br>$H_b$: 2.03-1.94, m, 1H |
| 3 | 58.2, $CH_2$ | $H_a$: 4.12-4.04, m, 1H<br>$H_b$: 4.04-3.92, m, 1H |
| 4 | n.a. | n.a. |
| 5 | 48.5, $CH_2$ | $H_a$: 3.73, ddd, *J* = 13.0, 6.0 Hz, 1H<br>$H_b$: 3.68-3.57, m, 1H |
| 6 | 19.8, $CH_2$ | 1.94-1.87, m, 2H |
| 7 | 15.8, $CH_2$ | 1.85-1.74, m, 2H |
| 8 | 24.7, $CH_2$ | $H_a$: 2.96-2.84, m, 1H<br>$H_b$: 2.80-2.69, m, 1H |
| 8a | 189.6, C | n.a. |

**Table S7. Structural characterization of 6•HCl.**

| | 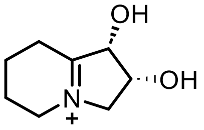 | | 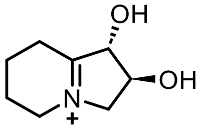<br>Isolated **8** contains C2-epimer spontaneously formed during the ioslation process | |
|---|---|---|---|---|
| | HRMS m/z calculated for $C_8H_{14}NO_2^+$ $[M+H]^+$ 156.1025, found 156.1020. | | | |
| Pos. | $\delta_C$ ($D_2O$) | $\delta_H$ ($D_2O$), multi, *J*, integration | $\delta_C$ ($D_2O$) | $\delta_H$ ($D_2O$), multi, *J*, integration |
| 1 | 78.1, CH | 5.00-4.97, m, 1H | 81.9, CH | 4.90-4.85, m, 1H |
| 2 | 67.6, CH | 4.58, dd, *J* = 4.8, 4.8 Hz, 1H | 67.6, CH | 4.49, ddd, *J* = 7.3, 6.2, 6.2 Hz, 1H |
| 3 | 65.3, $CH_2$ | $H_a$: 4.26-4.17, m, 1H<br>$H_b$: 3.96-3.88, m, 1H | 62.5, $CH_2$ | $H_a$: 4.26-4.17, m, 1H<br>$H_b$: 3.88-3.80, m, 1H |
| 4 | n.a. | n.a. | n.a. | |
| 5 | 48.7, $CH_2$ | 3.68-3.60, m, 2H | 48.7, $CH_2$ | 3.74-3.71 m, 2H |
| 6 | 19.8, $CH_2$ | 2.05-1.83, m, 2H | 19.6, $CH_2$ | 2.05-1.83, m, 2H |
| 7 | 15.5, $CH_2$ | 1.83-1.72, m, 2H | 15.5, $CH_2$ | 1.83-1.72, m, 2H |
| 8 | 25.0, $CH_2$ | $H_a$: 2.95-2.83, m, 1H<br>$H_b$: 2.83-2.68, m, 1H | 25.0, $CH_2$ | $H_a$: 2.95-2.83, m, 1H<br>$H_b$: 2.83-2.68, m, 1H |
| 8a | 190.3, C | n.a. | 190.3, C | n.a. |

**Table S8. Structural characterization of 9.**

| | |
|---|---|
| | 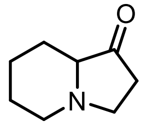 HRMS m/z calculated for $C_8H_{14}NO^+$ $[M+H]^+$ 140.1075, found 140.1071. |

| Pos. | $\delta_C$ ($CD_3OD$) | $\delta_H$ ($CD_3OD$), multi, *J*, integration |
|---|---|---|
| 1 | 215.16, C | n.a. |
| 2 | 36.5, $CH_2$ | 2.40 – 2.30, m, 2H |
| 3 | 51.01, $CH_2$ | $H_a$: 3.30 – 3.24, m, 1H<br>$H_b$: 2.57 – 2.50, m, 1H |
| 4 | n.a. | n.a. |
| 5 | 54.77, $CH_2$ | $H_a$: 3.17 – 3.09, m, 1H<br>$H_b$: 2.29 – 2.25, m, 1H |
| 6 | 25.88, $CH_2$ | $H_a$: 1.72 – 1.65, m, 1H<br>$H_b$: 1.61 – 1.52, m, 1H |
| 7 | 24.63, $CH_2$ | $H_a$: 1.89 – 1.81, m, 1H<br>$H_b$: 1.38 – 1.27, m, 1H |
| 8 | 25.94, $CH_2$ | $H_a$: 1.98 – 1.89, m, 1H<br>$H_b$: 1.72 – 1.65, m, 1H |
| 8a | 69.87, CD | n.a. |

**Table S9. Structural characterization of 11.**

<table>
<tr><td></td><td colspan="2">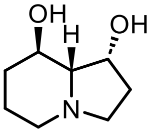<br>$J_{1,8a}$ = 4.5 Hz, the *cis* configuration between H-1 and H-8a<br>$J_{8,8a}$ = 9.6 Hz, the *trans* configuration between H-8 and H-8a<br>$[\alpha]_D^{20}$ = -38.0 (*c* = 0.2 in $H_2O$).<br>HRMS m/z calculated for $C_8H_{16}NO_2{}^+$ $[M+H]^+$ 158.1181, found 158.1178.</td></tr>
<tr><td>Pos.</td><td>$\delta_C$ ($CD_3OD$)</td><td>$\delta_H$ ($CD_3OD$), multi, *J*, integration</td></tr>
<tr><td>1</td><td>69.9, CH</td><td>4.38, ddd, *J* = 6.9, 4.5, 1.9 Hz, 1H</td></tr>
<tr><td>2</td><td>31.6, $CH_2$</td><td>$H_a$: 2.20, dddd, *J* = 14.0, 8.3, 8.3, 2.1 Hz, 1H<br>$H_b$: 1.61-1.51, m, 1H</td></tr>
<tr><td>3</td><td>52.2, $CH_2$</td><td>$H_a$: 3.02, ddd, *J* = 9.2, 2.2 Hz, 1H<br>$H_b$: 2.08, q, *J* = 9.2 Hz, 1H</td></tr>
<tr><td>4</td><td>n.a.</td><td>n.a.</td></tr>
<tr><td>5</td><td>51.6, $CH_2$</td><td>$H_a$: 2.97-2.85, m, 1H<br>$H_b$: 1.88, ddd, *J* = 13.3, 11.5, 2.9 Hz, 1H</td></tr>
<tr><td>6</td><td>23.2, $CH_2$</td><td>$H_a$: 1.70-1.64, m, 1H<br>$H_b$: 1.51-1.40, m, 1H</td></tr>
<tr><td>7</td><td>32.4, $CH_2$</td><td>$H_a$: 2.04-1.97, m, 1H<br>$H_b$: 1.26-1.14, m, 1H</td></tr>
<tr><td>8</td><td>66.4, CH</td><td>3.70, ddd, J = 10.4, 10.4, 4.7 Hz, 1H</td></tr>
<tr><td>8a</td><td>73.4, CH</td><td>1.70, dd, J = 9.6, 4.4 Hz, 1H</td></tr>
</table>

**Table S10. Structural characterization of 12.**

| | | |
|---|---|---|
| | 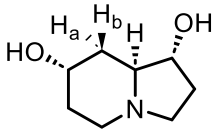<br>$J_{1,8a}$ = 7.9 Hz, the *trans* configuration between H-1 and H-8a<br>$J_{8\text{-}a,8a}$ = 3.0 Hz, the *cis* configuration between $H_a$-8 and H-8a<br>$J_{7,8\text{-}a}$ = 11.9 Hz, the *trans* configuration between H-7 and $H_a$-8<br>$[\alpha]_D^{20}$ = -27.6 (*c* = 0.5 in methanol).<br>HRMS m/z calculated for $C_8H_{16}NO_2^+$ $[M+H]^+$ 158.1181, found 158.1176. | |
| Pos. | $\delta_C$ ($D_2O$) | $\delta_H$ ($D_2O$, multi, *J*, integration |
| 1 | 74.8, CH | 3.80, ddd, *J* = 9.6, 7.9, 4.8 Hz, 1H) |
| 2 | 30.1, $CH_2$ | $H_a$: 2.16-2.07, m, 1H<br>$H_b$: 1.50, dddd, *J* = 14.1, 9.5, 4.8, 2.2 Hz, 1H |
| 3 | 51.4, $CH_2$ | $H_a$: 2.86, ddd, *J* = 9.2, 9.2, 2.3 Hz, 1H)<br>$H_b$: 2.43-2.29, m, 1H |
| 4 | n.a. | n.a. |
| 5 | 48.9, $CH_2$ | $H_a$: 2.79-2.71, m, 1H<br>$H_b$: 2.43-2.29, m, 1H |
| 6 | 30.5, $CH_2$ | 1.71-1.59, m, 2H |
| 7 | 63.9, CH | 4.16-4.09, m, 1H |
| 8 | 34.3, $CH_2$ | $H_a$: 2.03-1.94, m, 1H<br>$H_b$: 1.41, ddd, *J* = 14.3, 11.9, 2.8 Hz, 1H |
| 8a | 63.6, CH | 2.19, ddd, *J* = 11.6, 8.0, 3.0 Hz, 1H |

**Table S11. Structural characterization of 13.**

| | 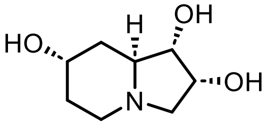<br>$J_{1,8a}$ = 9.1 Hz, the *trans* configuration between H-1 and H-8a<br>$J_{1,2}$ = 7.0 Hz, the *cis* configuration between H-1 and H-2<br>$[\alpha]_D^{20}$ = -24.0 ($c$ = 0.3 in $H_2O$).<br>HRMS m/z calculated for $C_8H_{16}NO_3^+$ $[M+H]^+$ 174.1130, found 174.1126. | |
|---|---|---|
| Pos. | $\delta_C$ ($D_2O$) | $\delta_H$ ($D_2O$), multi, $J$, integration |
| 1 | 74.5, CH | 3.56, dd, $J$ = 9.1, 7.0 Hz, 1H |
| 2 | 66.8, CH | 4.20-4.11, m, 1H |
| 3 | 60.1, $CH_2$ | $H_a$: 3.40, dd, $J$ = 10.8, 6.9 Hz, 1H<br>$H_b$: 2.18, dd, $J$ = 10.7, 5.2 Hz, 1H |
| 4 | n.a. | n.a. |
| 5 | 46.6, $CH_2$ | $H_a$: 2.81-2.74, m, 1H<br>$H_b$: 2.44-2.32, m, 1H |
| 6 | 30.7, $CH_2$ | 1.77-1.63, m, 2H |
| 7 | 63.8, CH | 4.20-4.11, m, 1H |
| 8 | 34.4, $CH_2$ | $H_a$: 2.02, q, $J$ = 13.9, 2.7 Hz, 1H)<br>$H_b$: 1.44, ddd, $J$ = 14.3, 11.8, 2.8 Hz, 1H |
| 8a | 60.1, CH | 2.44-2.32, m, 1H |

**Table S12. Structural characterization of 3'.**

| | | |
|---|---|---|
| | 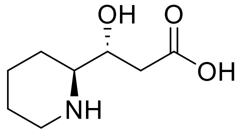<br>$J_{1,8a}$ = 3.2 Hz, the *erythro* configuration between N-4 and O-2<br>HRMS m/z calculated for $C_8H_{16}NO_3^+$ $[M+H]^+$ 174.1130, found 174.1132. | |
| Pos. | $\delta_C$ ($D_2O$) | $\delta_H$ ($D_2O$), multi, *J*, integration |
| 1 | 67.9, CH | 4.14, ddd, *J* = 6.9, 6.9, 3.2 Hz, 1H |
| 2 | 39.4, $CH_2$ | 2.38, d, *J* = 7.0 Hz, 2H |
| 3 | 177.4, C | n.a. |
| 4 | n.a. | n.a. |
| 5 | 44.9, $CH_2$ | $H_a$: 3.35, d, *J* = 12.4 Hz, 1H<br>$H_b$: 2.95, dd, *J* = 13.0, 13.0 Hz, 1H |
| 6 | 21.8, $CH_2$ | $H_a$: 1.91-1.77, m, 1H<br>$H_b$: 1.62-1.50, m, 1H |
| 7 | 21.7, CH | $H_a$: 1.91-1.77, m, 1H<br>$H_b$: 1.51-1.39, m, 1H |
| 8 | 21.4, $CH_2$ | $H_a$: 1.91-1.77, m, 1H<br>$H_b$: 1.51-1.39, m, 1H |
| 8a | 60.0, CH | 3.15, d, *J* = 12.2 Hz, 1H |

**Table S13. Structural characterization of 2’.**

| | | |
|---|---|---|
| | 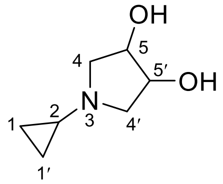 HRMS m/z calculated for $C_7H_{14}NO_2^+$ [M+H]$^+$ 144.1025, found 144.1026. | |
| Pos. | $\delta_C$ ($D_2O$) | $\delta_H$ ($D_2O$), multi, *J*, integration |
| 1, 1′ | 70.3, $CH_2$ | $H_a$: 0.48-0.39, m, 2H<br>$H_b$: 0.37-0.30, m, 2H |
| 2 | 36.8, CH | 1.96-1.88, m, 1H |
| 3 | n.a. | n.a. |
| 4, 4′ | 57.8, $CH_2$ | $H_a$: 3.10, dd, *J* = 11.2, 5.0 Hz, 2H<br>$H_b$: 2.64, dd, *J* = 11.1, 4.5 Hz, 2H |
| 5, 5′ | 4.3, CH | 4.21-4.07, m, 2H |

### 4.Figures

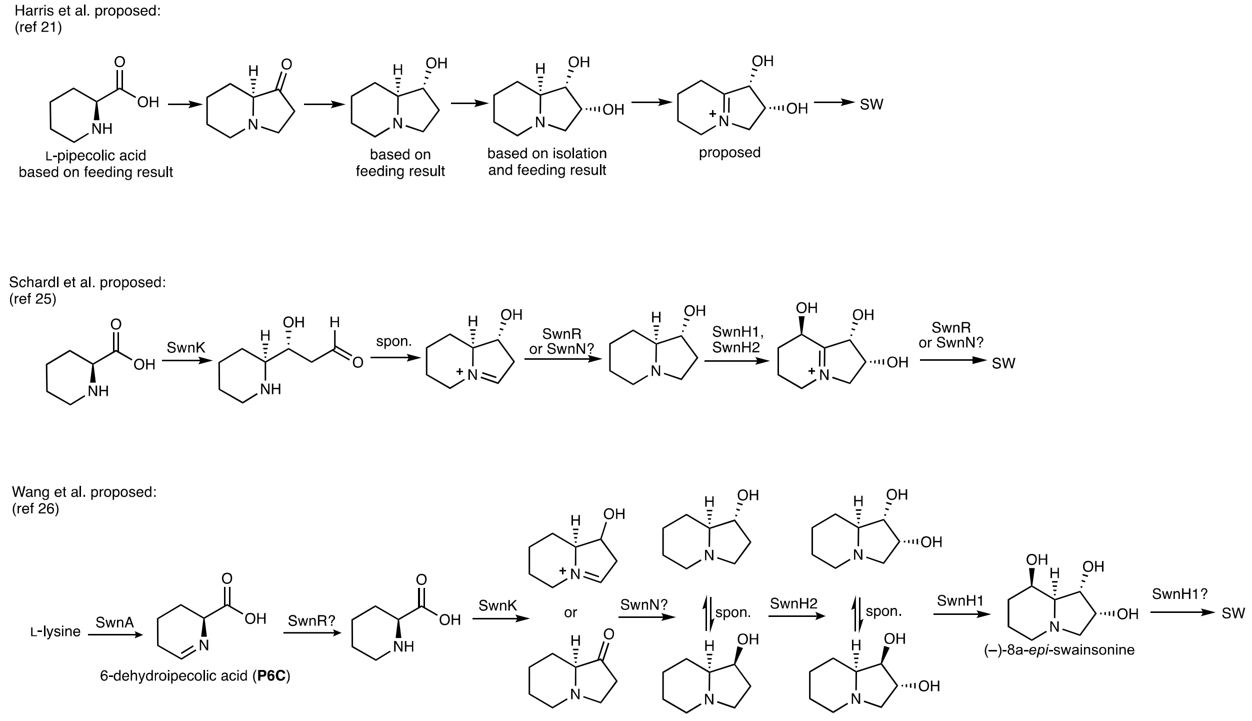

**Figure S1.** Different biosynthetic proposals for swainsonine.

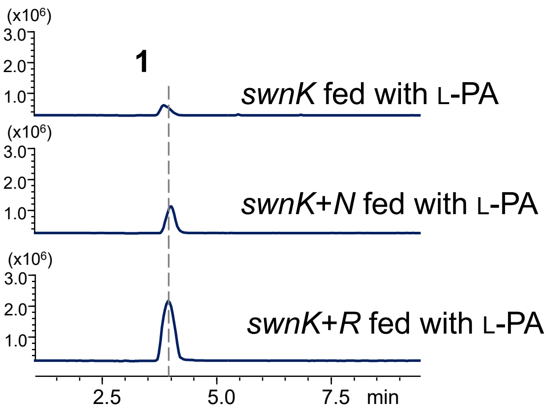

**Figure S2.** Extracted ion chromatograms (EIC) shows the production of **1** derived from fed L-pipecolic acid.

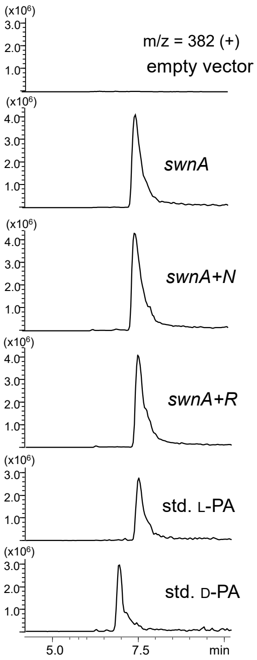

**Figure S3.** Extracted ion chromatograms show the heterologous formation of L-PA in *A. nidulans* is independent of either SwnN or SwnR.

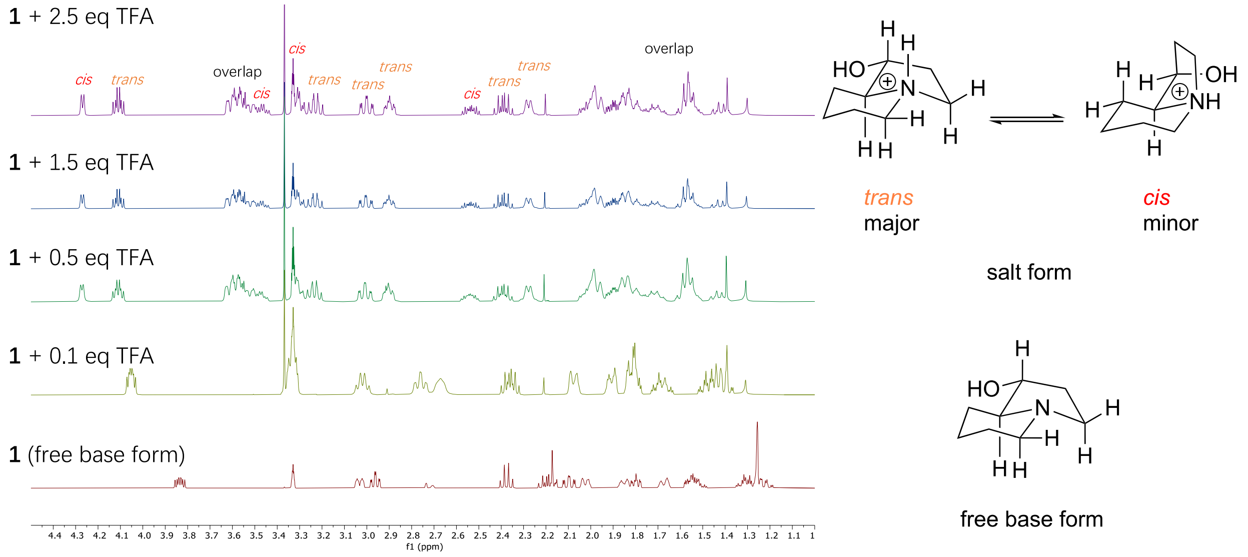

**Figure S4. Conformational isomers observed with 1 in its salt form**.
Indolizidine alkaloids are known to adopt different conformations in equilibria.[9] Two stable conformational isomers of **1** were resolved on NMR when **1** was in its salt form. Conformation assignments were made based on key NOE signals, which are summarized in Table S3 and S4.

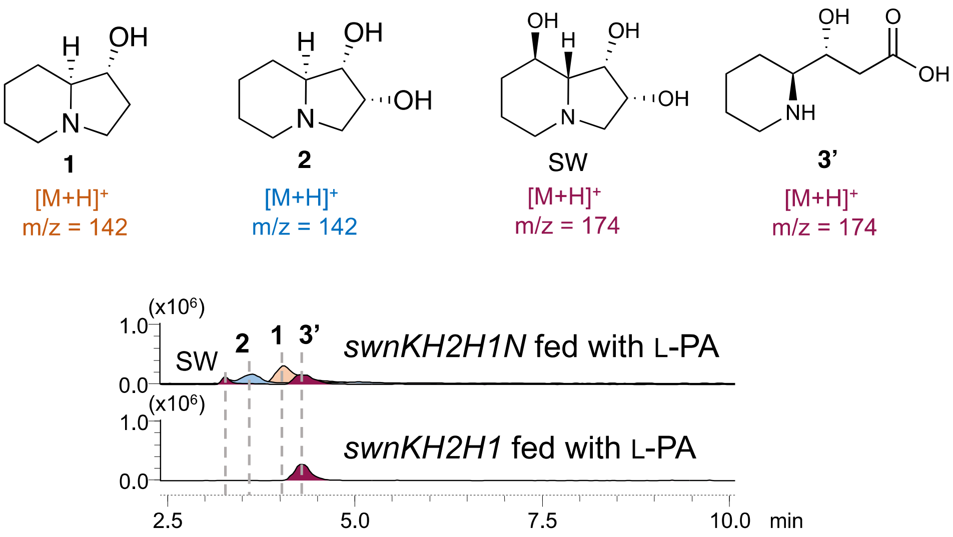

**Figure S5.** Feeding of L-PA to *A. nidulans* transformed with or without of *swnN* gene. Compound 3' was isolated and its structural determination is shown in Table S12.

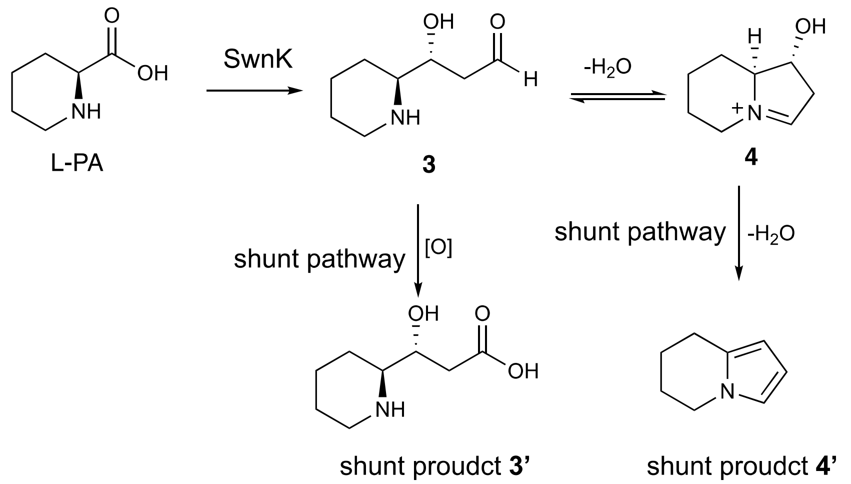

**Figure S6.** Shunt products observed in vivo.

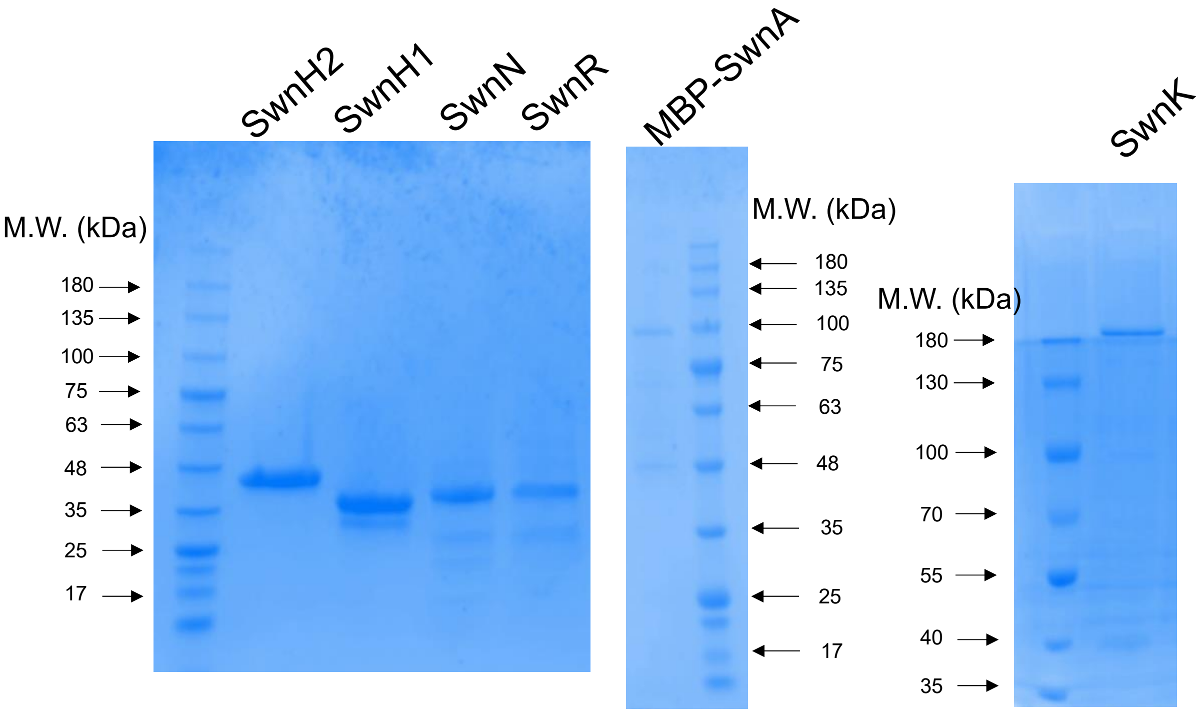

**Figure S7.** SDS-page gels of purified proteins studied in this work.

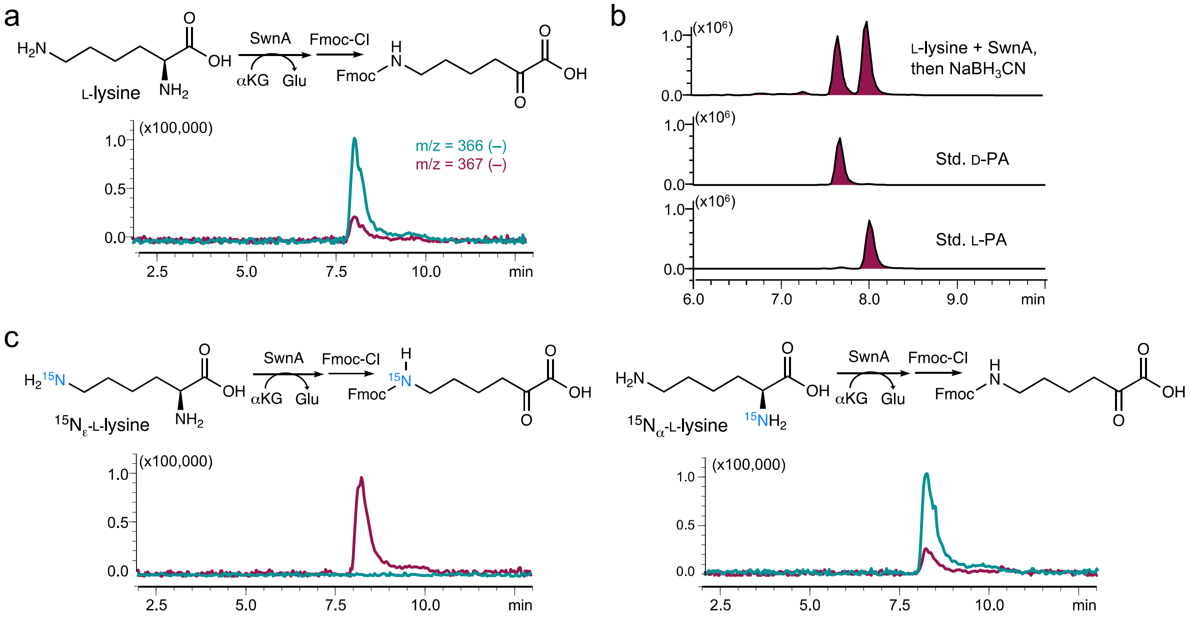

**Figure S8.** SwnA is a lysine 2-aminotransferase.

(a) Fmoc-derivatization of SwnA-catalyzed transamination product from L-lysine. (b) Chiral analysis of the chemically reduced SwnA enzymatic product after derivatization with Marfey reagents. (c) Isotope-labeling experiment confirms that SwnA is a lysine 2-aminotransferase.

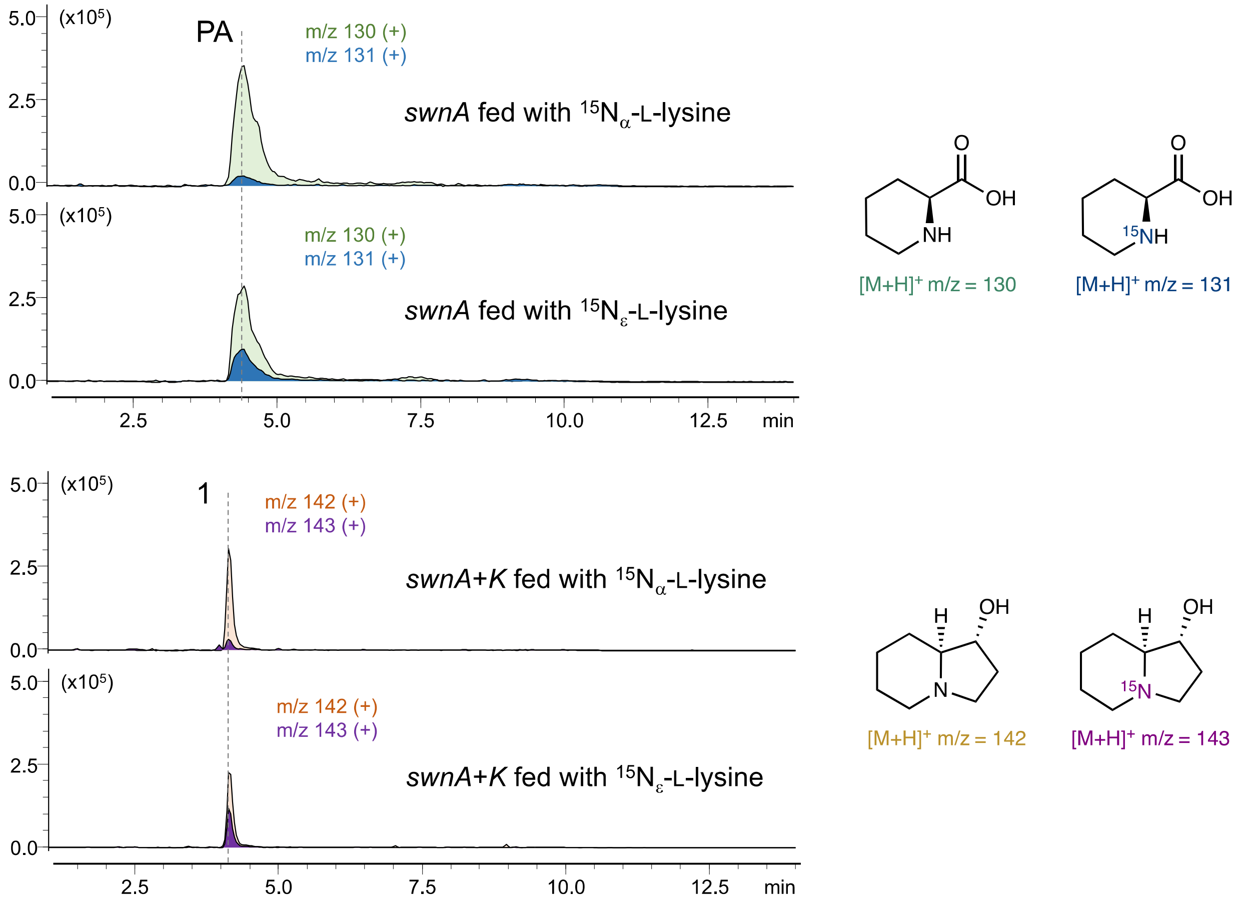

**Figure S9.** Feeding of isotope-labeled lysine to *A. nidulans* expressing SwnA and SwnK.

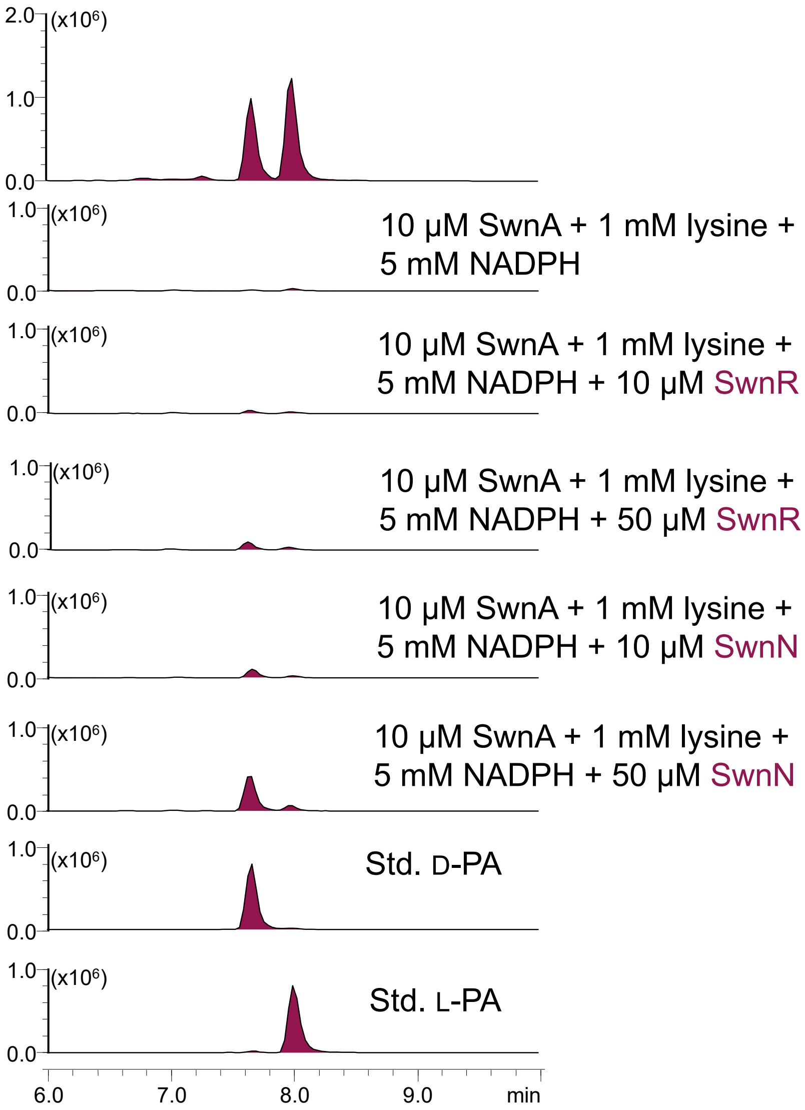

**Figure S10.** SwnA-Coupled enzymatic assays.

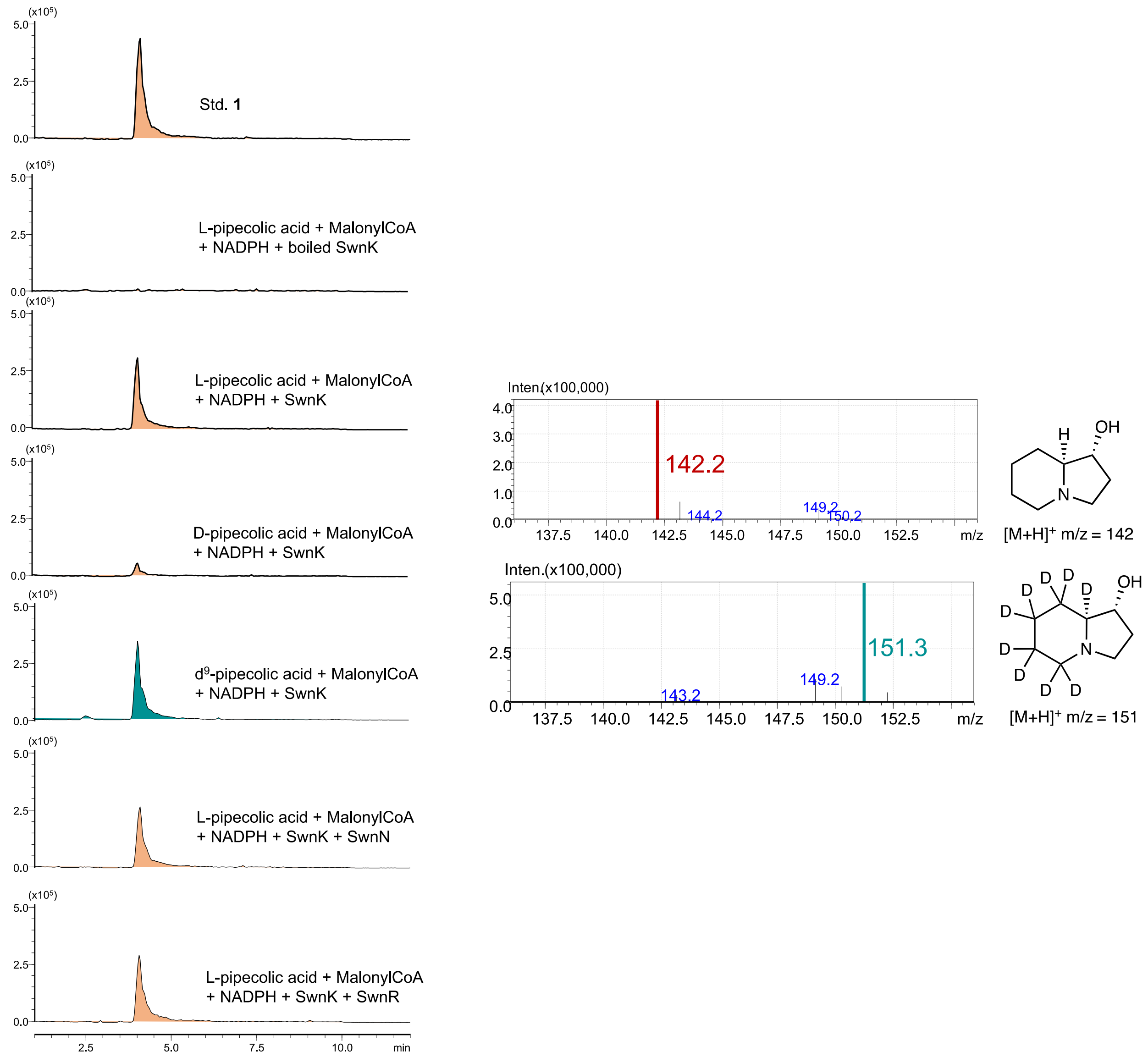

**Figure S11.** Biochemical characterization of SwnK in vitro.

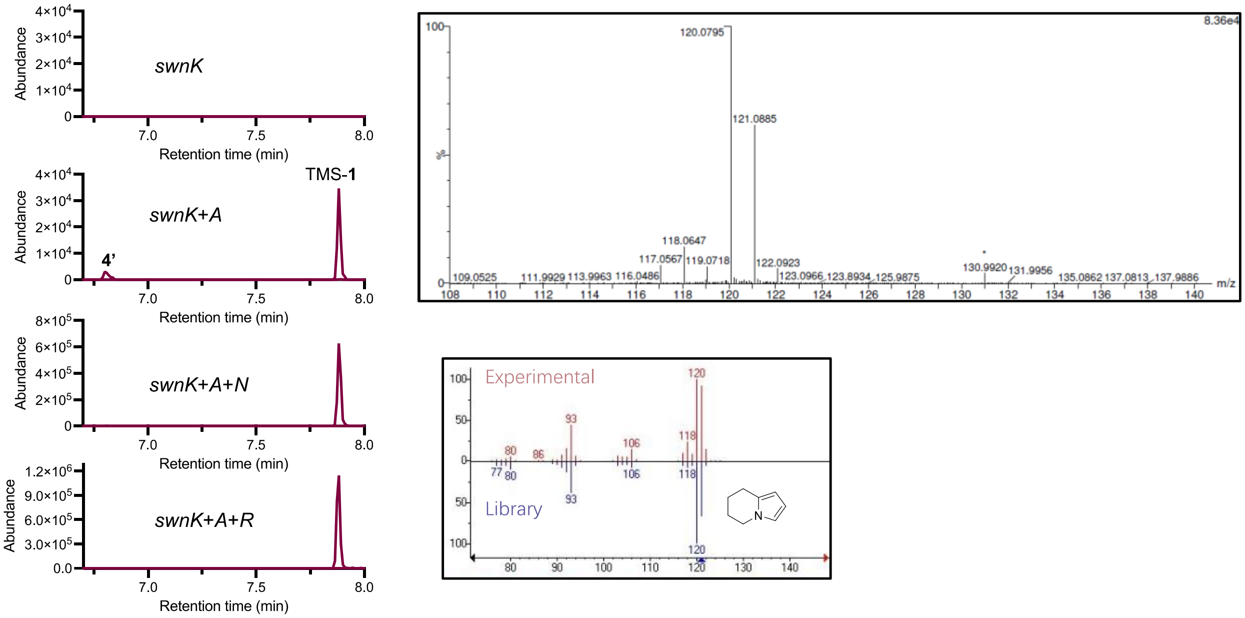

**Figure S12.** GC-MS analysis of the metabolite profile revealed the shunt-product **4**‘.

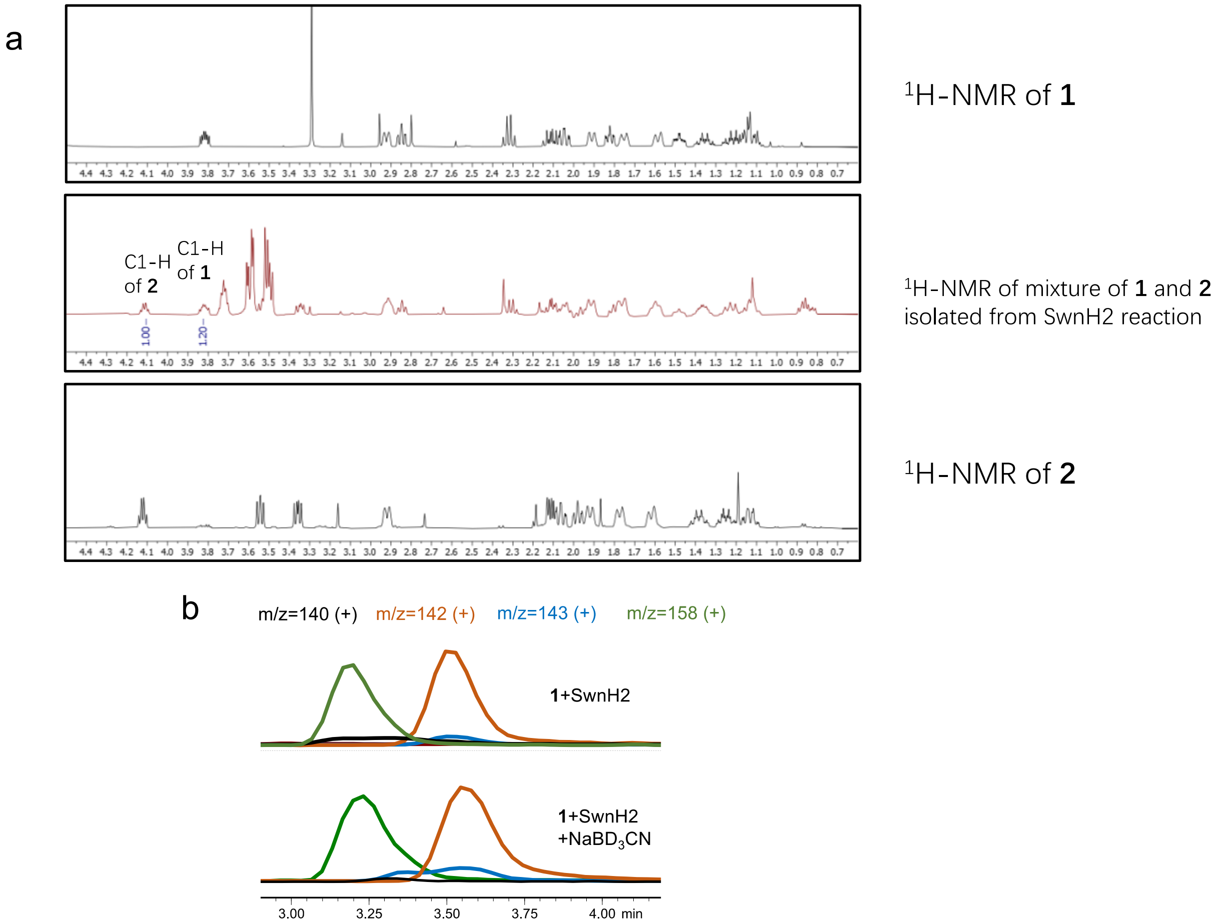

**Figure S13.** Determining the ratio between **2** and **5** during the initial reaction of SwnH2-catalyzed oxidation of **1**

Compound **5** was reduced by $NaBD_3CN$ in situ to make C8a-deuterated **1** in order to ease the purification and accurate quantification. No formation of **6** was observed under this condition. (a) The ratio between **1** and **2** was determined to be 1.2:1 by NMR. (b) The ratio between the leftover **1** and deuterated **1** was determined to be 83:17 by mass-spectrometry. The overall ratio between **5** and **2** is calculated to be 18:82.

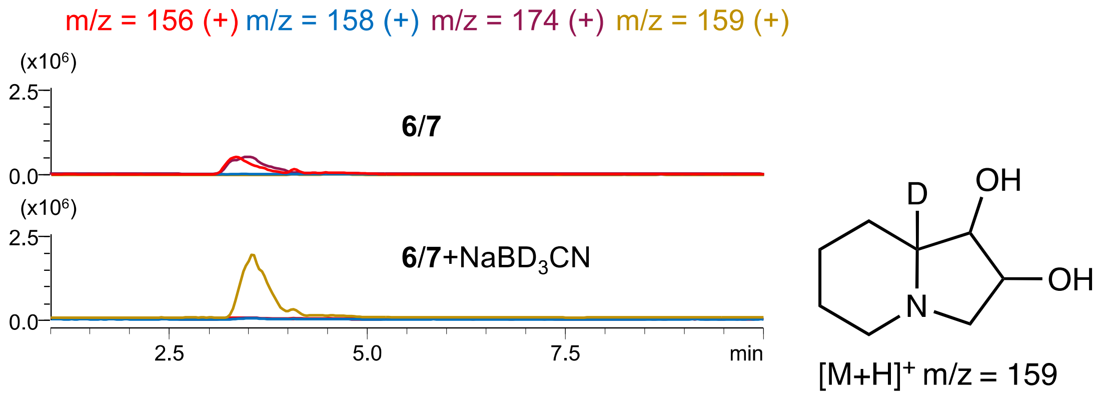

**Figure S14.** Chemical reduction of **6** and **7** by $NaBD_3CN$.

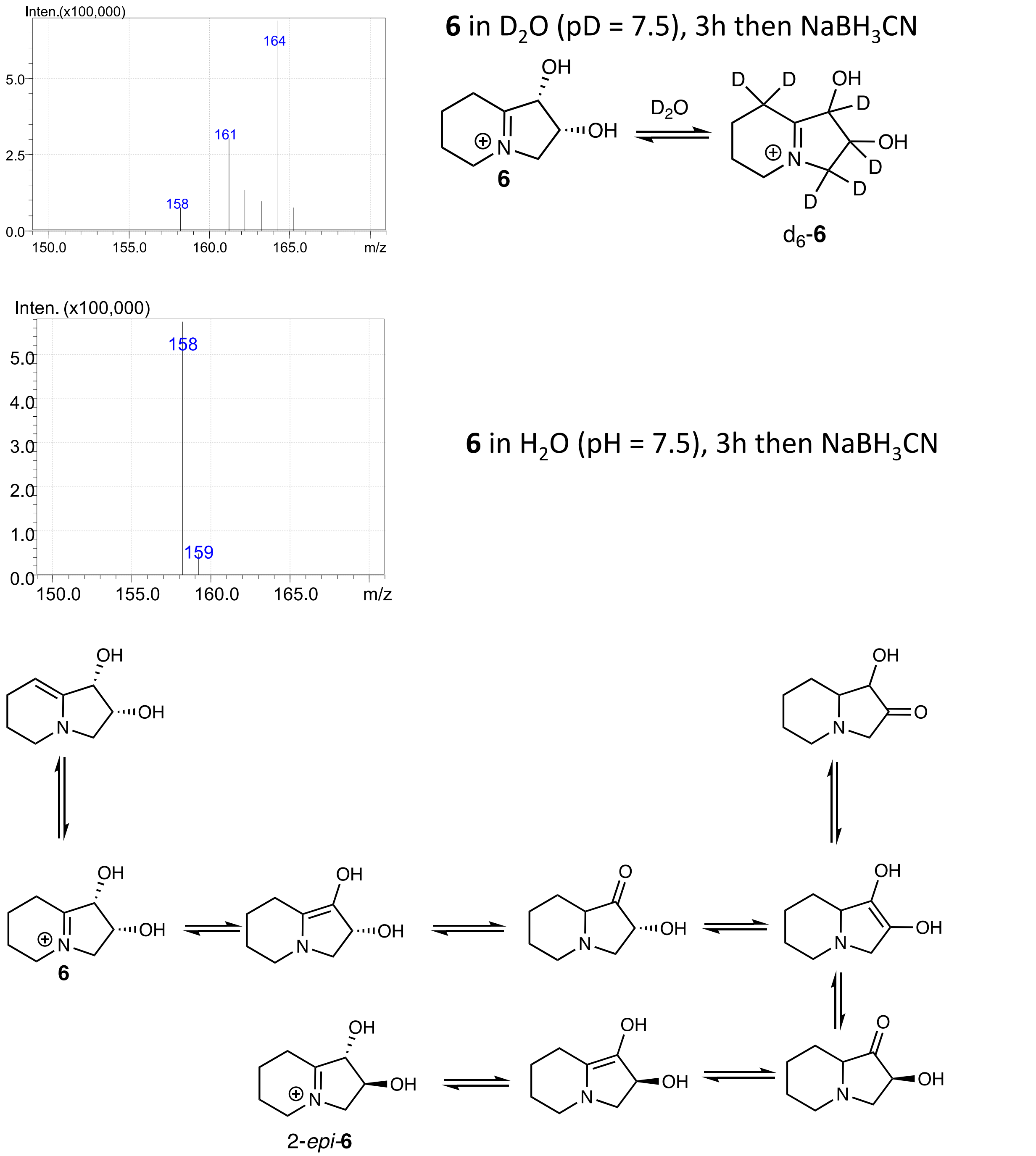

**Figure S15.** Spontaneous tautomerization of **6** is supported by the solvent isotopic labeling of **6** in $D_2O$.

In theory, up to 6 deuterium atoms can be incorporated into **6** through spontaneous tautomerization in $D_2O$. Indeed, trapping the deuterated **6** by chemical reduction revealed a mass shift of +6 Da.

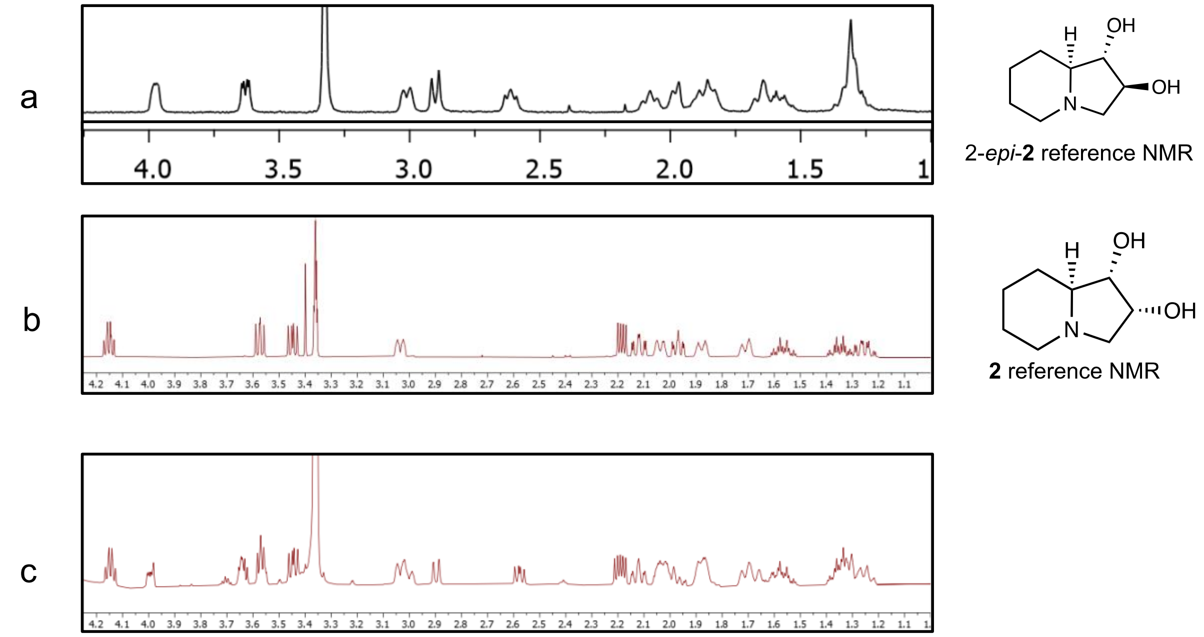

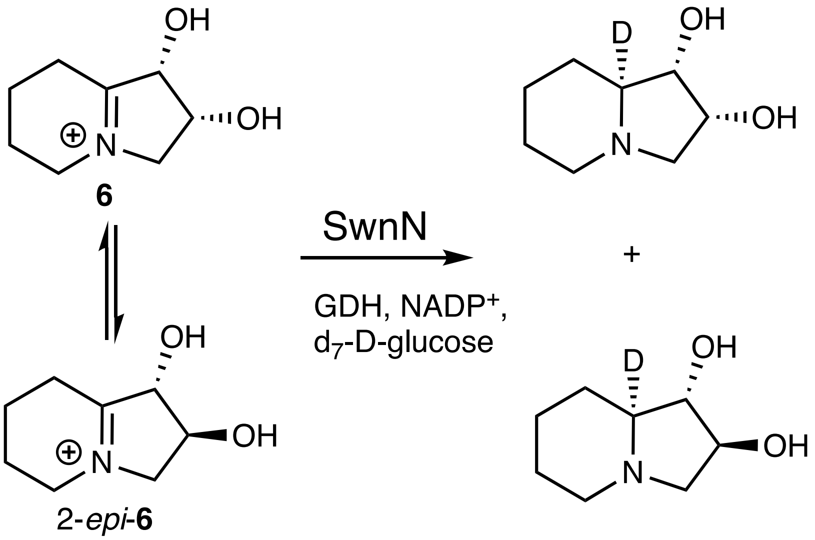

**Figure S16.** Structural characterization of products isolated from SwnN-catalyzed reduction of **6** (containing 33% 2-*epi*-**6**).
a) Literature reported $^1$H NMR (400 MHz) of 2-*epi*-**2** (also known as lentiginosine) in $CD_3OD$.[7] b) $^1$H NMR (400 MHz) of standard **2** in $CD_3OD$. c) $^1$H NMR (400 MHz) of the products isolated from SwnN-catalyzed reduction of compound **6** in $CD_3OD$**.** The reaction was carried out with deuterated NADPH, which was enzymatically synthesized in situ using glucose dehydrogenase (GDH) and d7-D-glucose.

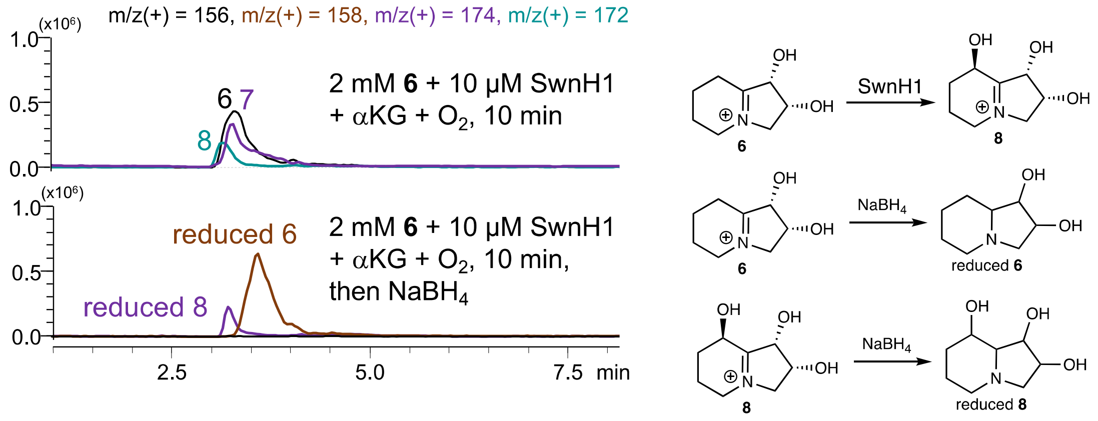

**Figure S17.** Chemical reduction to trap intermediate **8.**

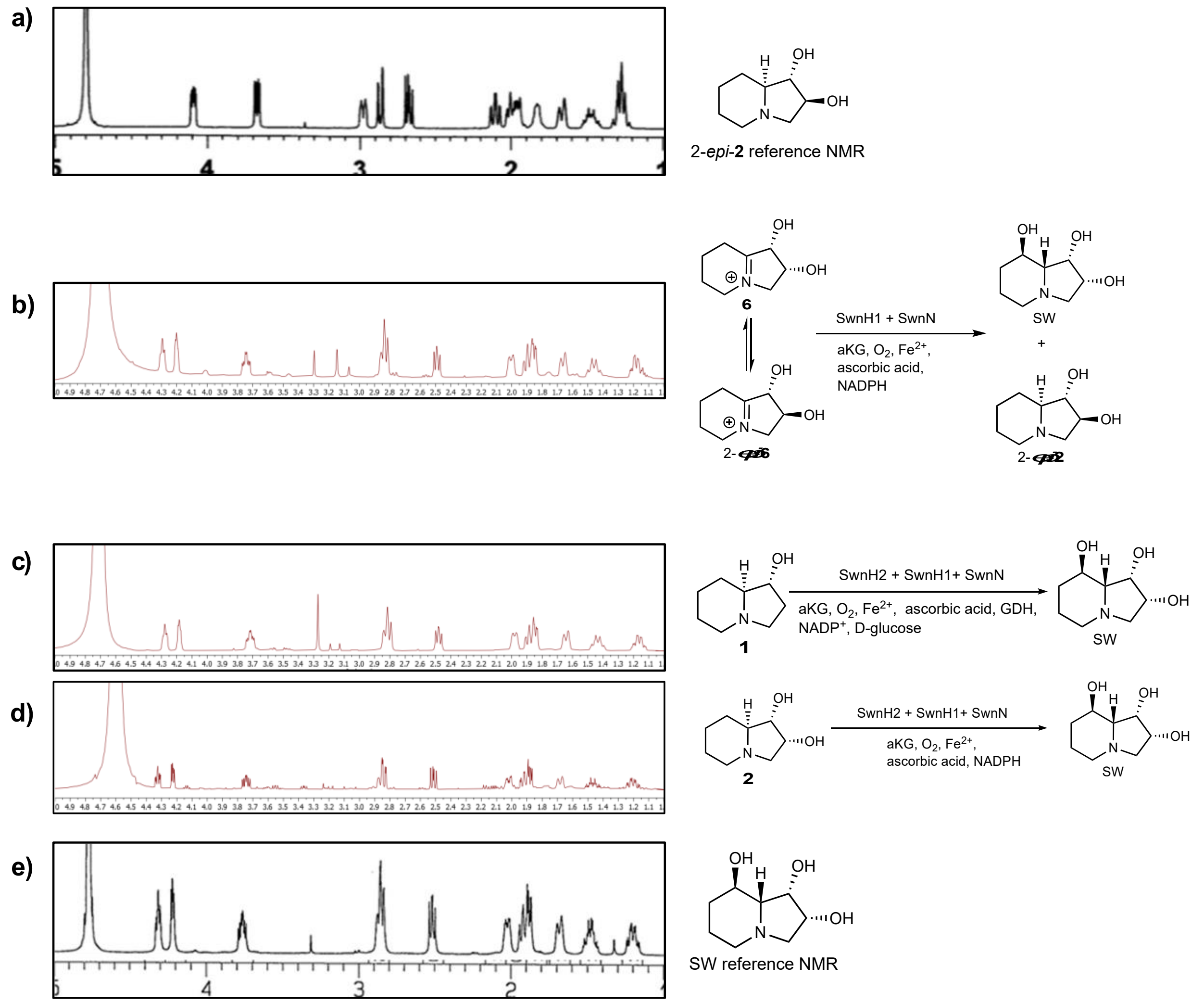

**Figure S18.** Structural confirmation of SW isolated from different enzymatic reactions.
a) Literature reported (400 MHz) of 2-*epi*-**2** (also known as lentiginosine) in $D_2O$.[10] b) $^1$H NMR (500 MHz) of SW in $D_2O$, which was isolated from the SwnH1-SwnN coupled enzymatic reaction with substrate **6** (containing 33% 2-*epi*-**6**). c) $^1$H NMR (500 MHz) of SW in $D_2O$, which was prepared from **2** through the cascade enzymatic assay using SwnH2, SwnH1, and SwnN. d) $^1$H NMR (500 MHz) of SW in $D_2O$, which was prepared from **1** through the cascade enzymatic assay using SwnH2, SwnH1, and SwnN. e) Literature reported $^1$H NMR (500 MHz) of SW in $D_2O$.[11]

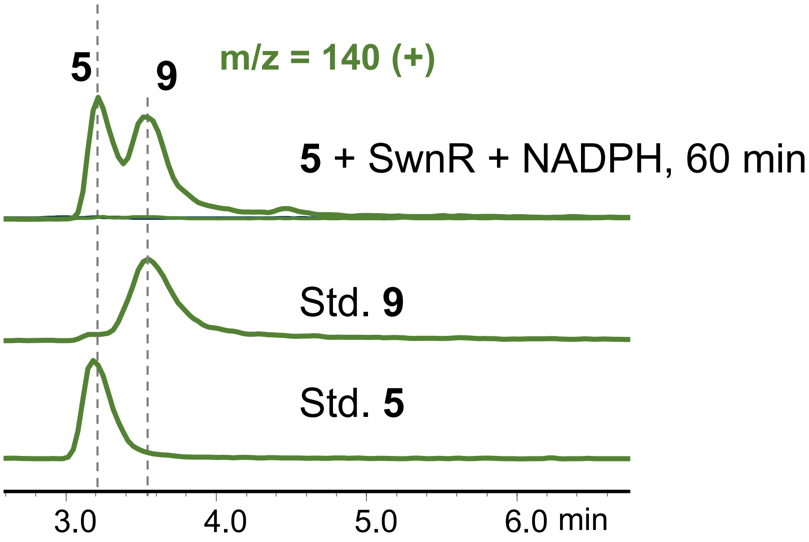

**Figure S19.** Compound **5** spontaneously tautomerizes into compound **9**.

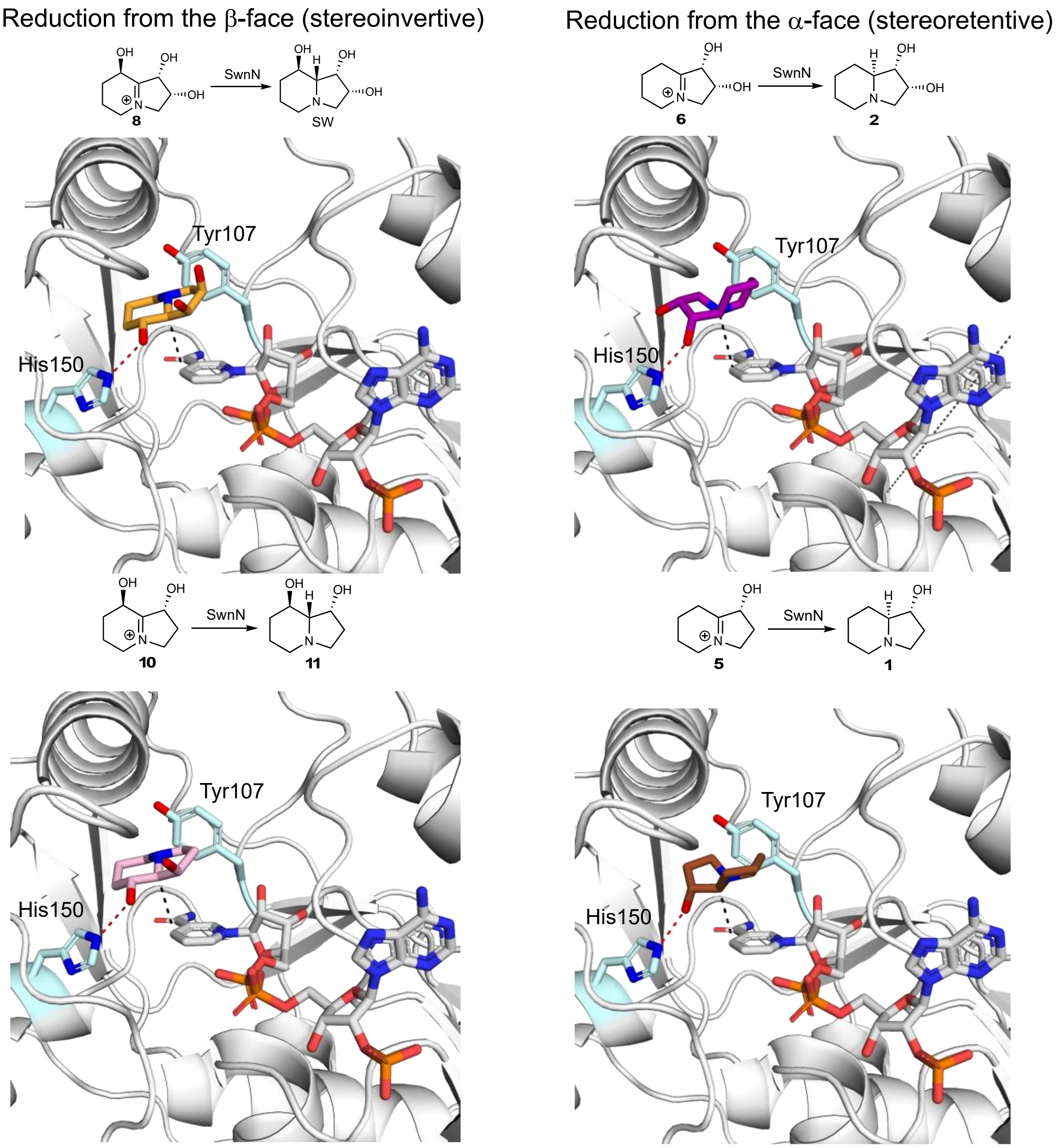

**Figure S20.** Docking model to understand the substrate-dependent stereoselectivity of SwnN. A model SwnN-NADP+ complex was made by using AlphaFold3,[12] then different reaction products were manually docked to the active site pocket. The potential hydride transfer trajectories are shown as black dashes, and the key hydrogen bonding interaction with His150 are shown as red dashes. The model suggests Tyr107 may recognize the tertiary amine group through a cation-π interaction, while His150 may recognize the SwnH1-installed C8-OH group through a hydrogen bond interaction, thereby stereo-specifically reduce the iminium substrate from the β-face. In the absence of the SwnH1-installed C8-OH group, substrates bind to the same pocket with ring flipped such that the C1-OH group makes a hydrogen bond with His150, leading to the reduction from the α-face.

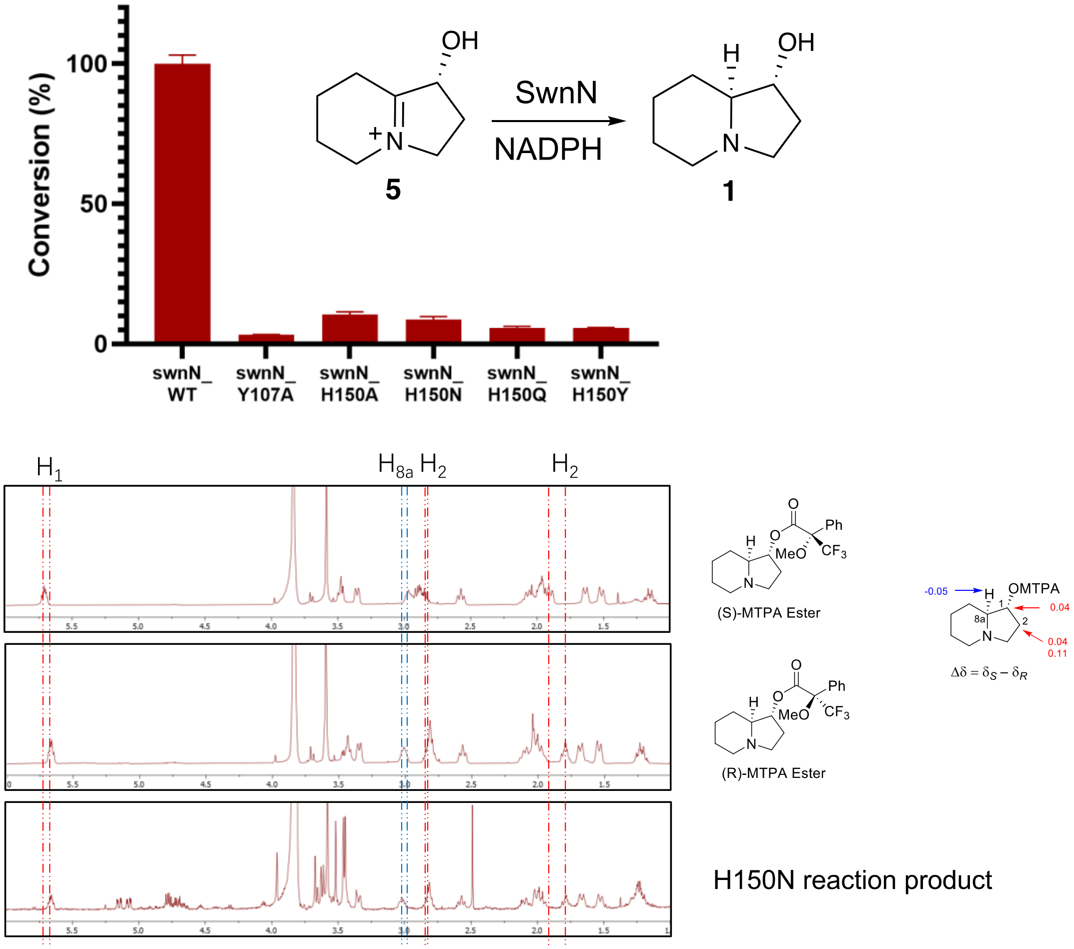

**Figure S21.** Mutagenesis study of SwnN demonstrates the importance of His150 and Tyr107. Different SwnN variants were prepared and their activities were evaluated using substrate 5. As shown by the graph above (data represent mean values ± s.d. from three independent experiments), all variants on Tyr107 and His150 show substantially compromised catalytic activity, consistent with their proposed catalytic importance. Because of the low catalytic activity, only the product from the reaction catalyzed by H150A mutant was isolated and structurally verified to be a single stereoisomer **1** by using Mosher's analysis.

SwnH2-catalyzed oxdiation of **1** and **2**: the ferryl intermediate approaches substrates through the α-face.

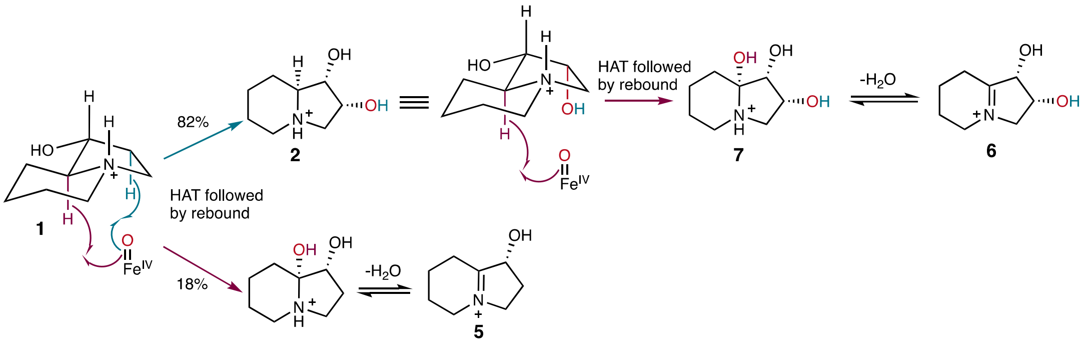

SwnH2-catalyzed oxdiation of **Ac-1**: the ferryl intermediate approaches substrates through the α-face.

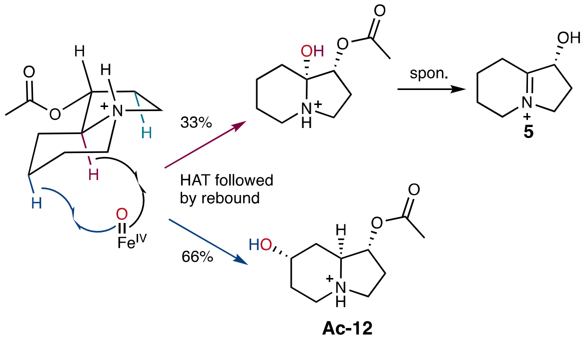

SwnH1-catalyzed hydroxylation of **6**: the ferryl intermediate approaches substrates through the β-face.

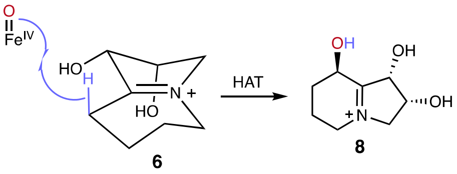

SwnH1-catalyzed amine desaturation of **2** must go through a different route because the $C_{8a}$-H is inaccessible.

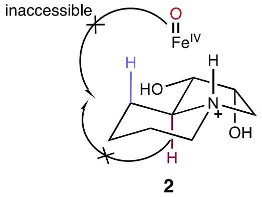

**Figure S22.** Proposed catalytic mechanism for SwnH2.

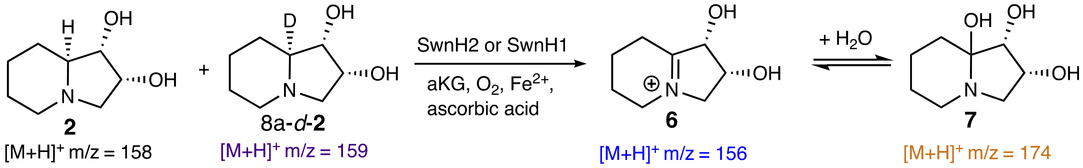

**2**/8a-*d*-**2** + SwnH2 (boiled), 60 min
Peak area (m/z +159): 27013743
Peak area (m/z +158): 25884201

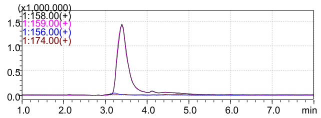

**2**/8a-*d*-**2** + SwnH2, 10 min
Peak area (m/z +159): 34464333
Peak area (m/z +158): 29451713

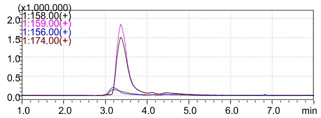

**2**/8a-*d*-**2** + SwnH2, 20 min
Peak area (m/z +159): 28904026
Peak area (m/z +158): 21906635

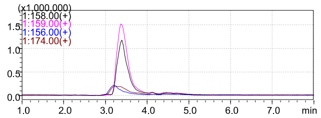

**2**/8a-*d*-**2** + SwnH2, 30 min
Peak area (m/z +159): 26327211
Peak area (m/z +158): 15047016

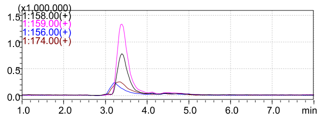

**2**/8a-*d*-**2** + SwnH2, 60 min
Peak area (m/z +159): 17970640
Peak area (m/z +158): 7127401

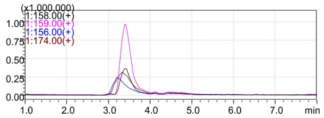

**2**/8a-*d*-**2** + SwnH1, 10 min
Peak area (m/z +159): 30395029
Peak area (m/z +158): 30698912

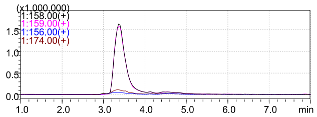

**2**/8a-*d*-**2** + SwnH1, 20 min
Peak area (m/z +159): 26016359
Peak area (m/z +158): 27464977

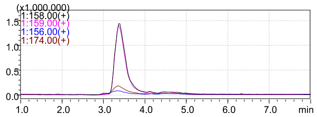

**2**/8a-*d*-**2** + SwnH1, 30 min
Peak area (m/z +159): 22917274
Peak area (m/z +158): 24145106

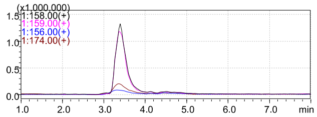

**2**/8a-*d*-**2** + SwnH1, 60 min
Peak area (m/z +159): 20421727
Peak area (m/z +158): 21022868

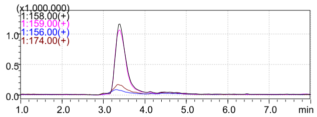

**Figure S23.** Investigating amine oxidation reaction of **2** using 8a-deuterium-labeled substrates. Reaction with SwnH2 shows a clear kinetic isotope effect, as the non-labeled substrate was consumed much faster than the labeled substrate. In contrast, no kinetic rate difference was observed with SwnH1, demonstrating that SwnH2-catalyzed oxidation of **2** experiences a different mechanism as SwnH1.

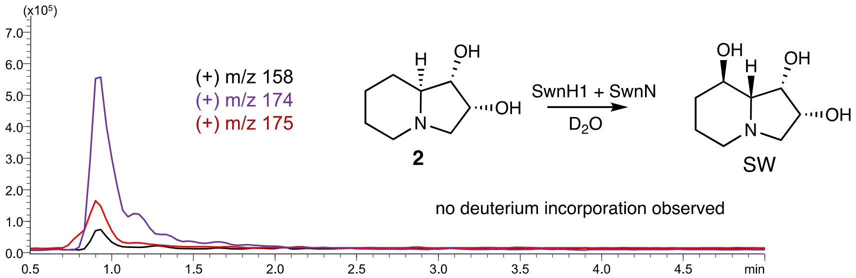

Disfavoring the following mechanism:

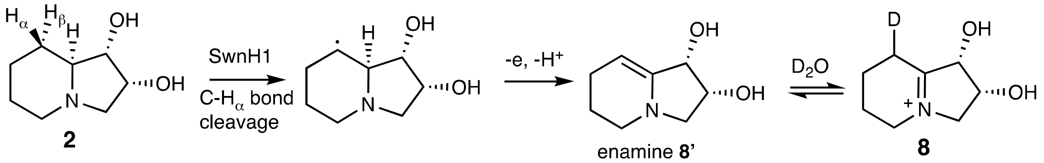

**Figure S24.** No solvent proton transfer is involved in SwnH1-catalyzed oxidation of **2**. The lack of deuterium incorporation disfavor the mechanistic proposal involving an enamine intermediate. Reaction condition: 1 mM substrate **2**, 10 mM αKG, 2 mM ascorbic acid, 0.2 mM $(NH_4)_2Fe(SO_4)_2$, 10 mM NADPH, 10 µM SwnH1, 10 µM SwnN, 50 mM KPi buffer (pD 7.5). Reaction was incubated at 30 ºC for 8 hours.

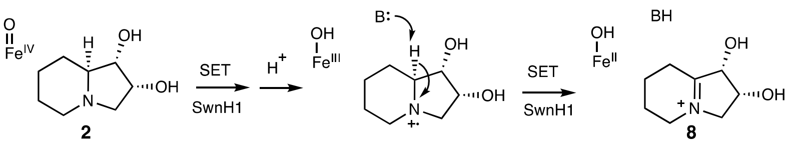

2

8

**Figure S25.** Proposed alternative mechanistic pathway for SwnH1-catalyzed amine desaturation of **2** through an aminyl radical intermediate

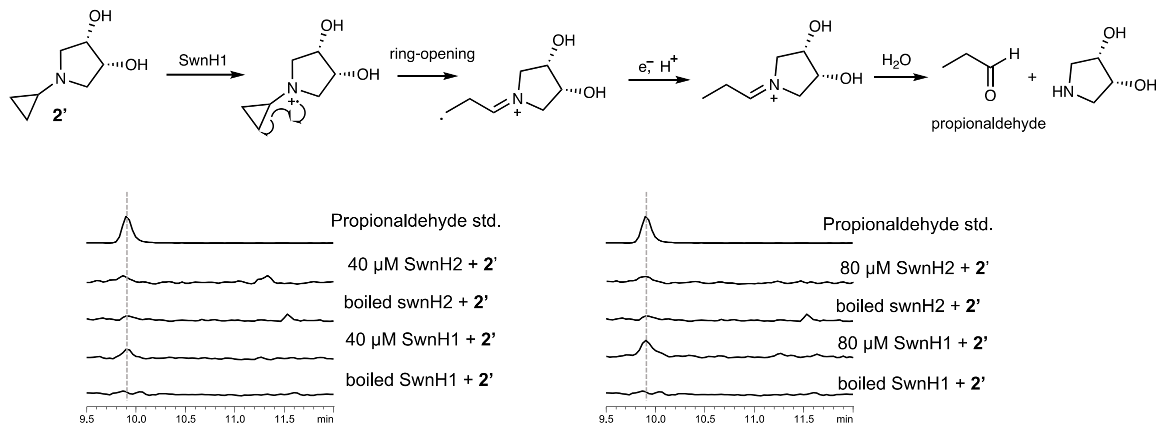

**Figure S26. A radical clock experiment with SwnH1 and SwnH2.**
Detection of ring-opening product, propionaldehyde, for SwnH1 but not SwnH2 supports an aminyl radical mechanism for SwnH1. Reaction condition: 5 mM substrate **2'**, 20 mM αKG, 10 mM ascorbic acid, 0.2 mM $(NH_4)_2Fe(SO_4)_2$, 40 or 80 μM enzyme, 50 mM KPi buffer (pH 6.5). Reaction was incubated at 30 ºC for 20 minutes.

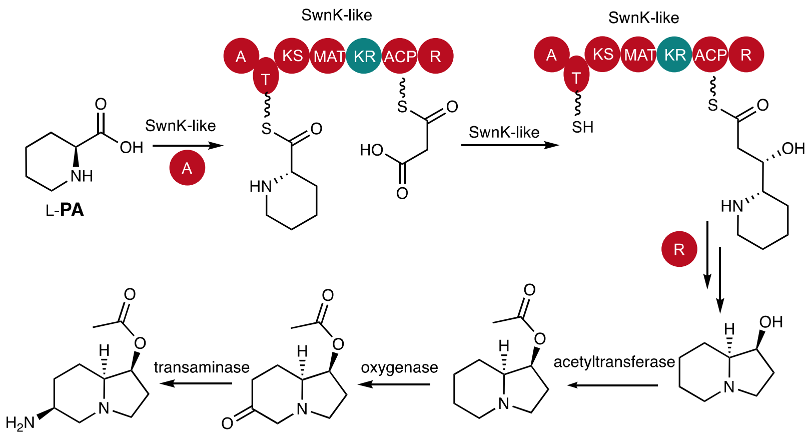

**Figure S27.** Proposed biosynthetic pathway for slaframine.

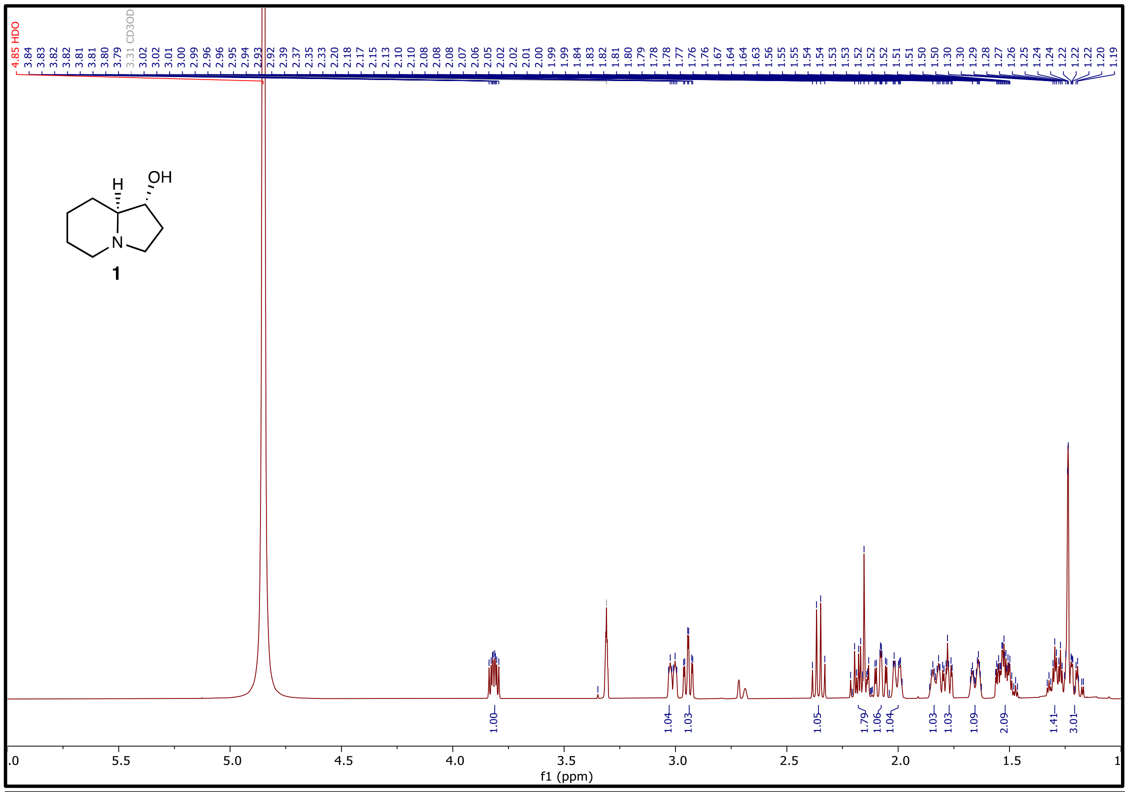

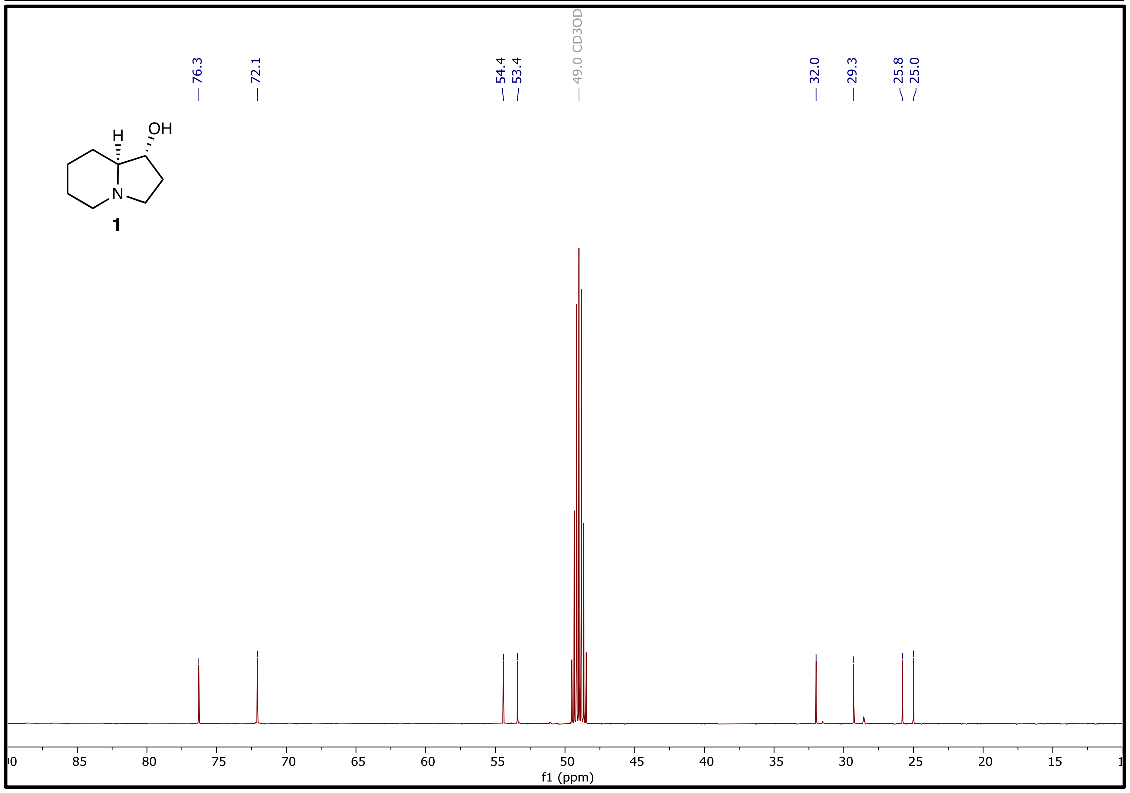

**Figure S28. $^1$H NMR (500 MHz) and $^{13}$C NMR (126 MHz) of 1 in $CD_3OD$.**

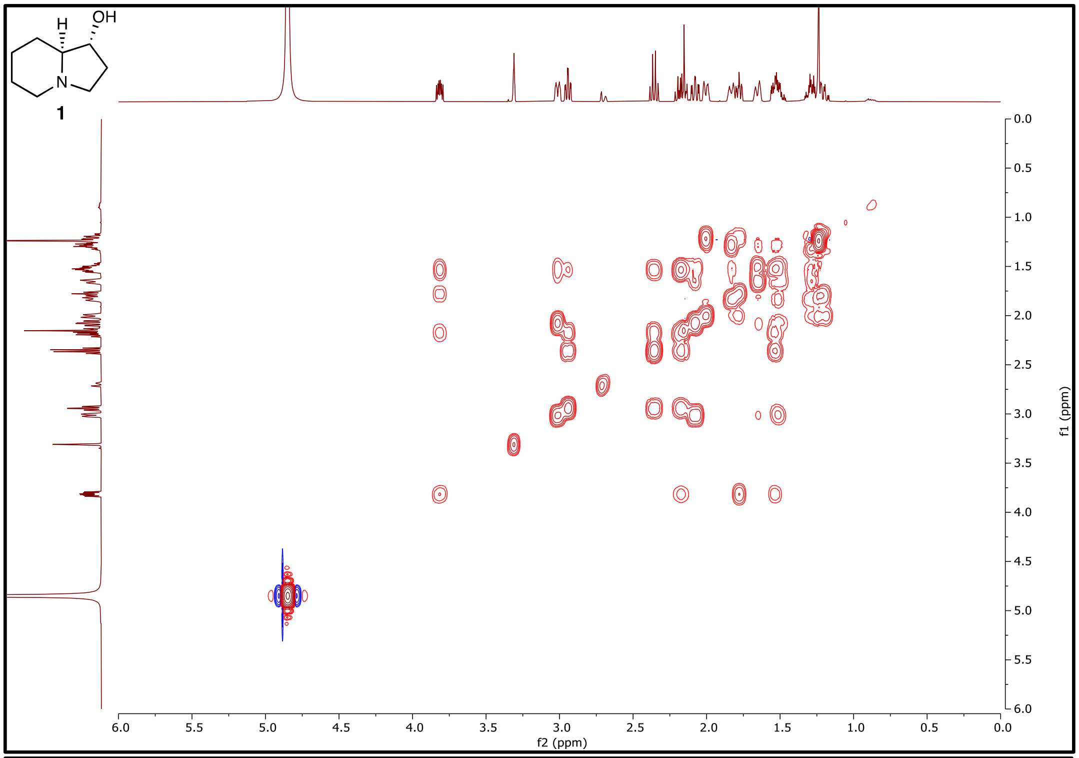

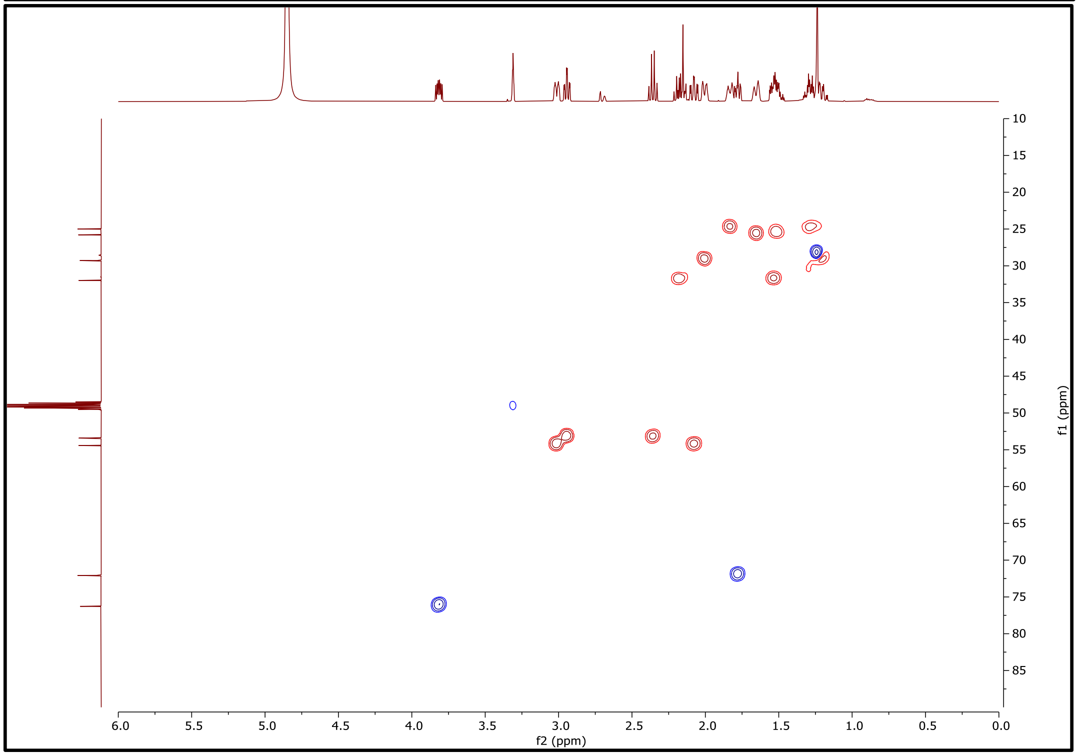

**Figure S29.** $^{1}H$-$^{1}H$ **COSY NMR (500 MHz) and** $^{1}H$-$^{13}C$ **HSQC NMR (500 MHz) of 1 in** $CD_3OD$**.**

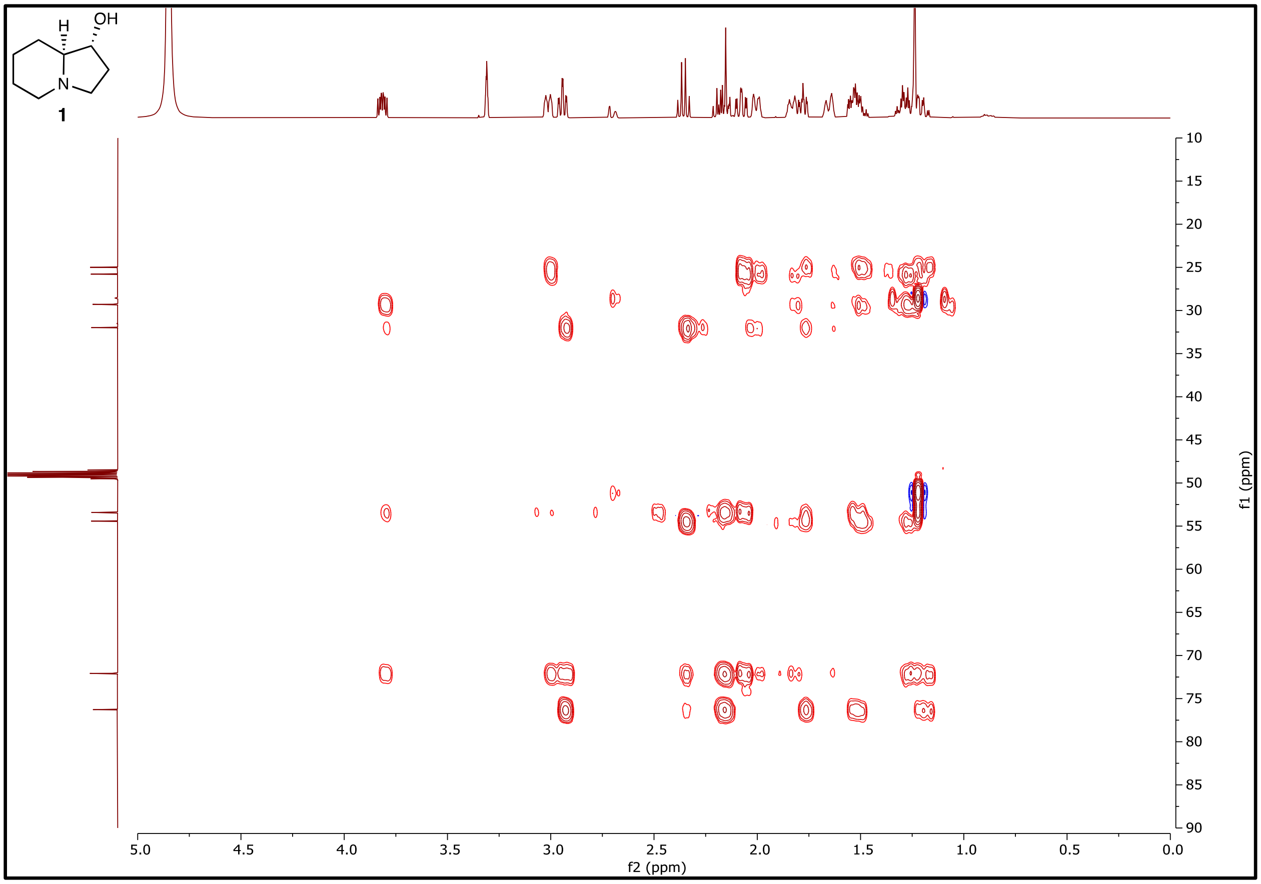

**Figure S30. $^{1}H$-$^{13}C$ HMBC NMR (500 MHz) of 1 in $CD_3OD$.**

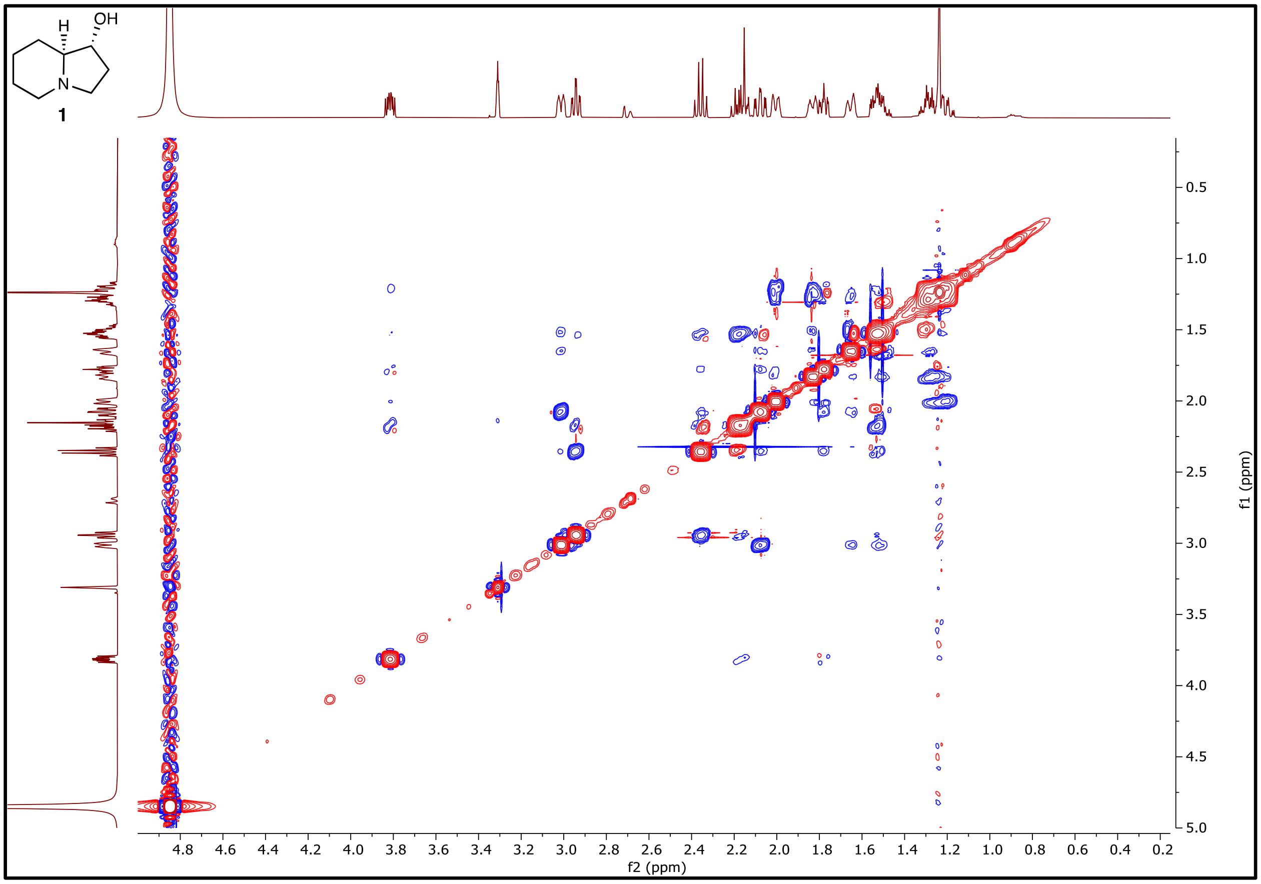

**Figure S31. $^{1}H$-$^{1}H$ NOESY NMR (500 MHz) of 1 in $CD_3OD$.**

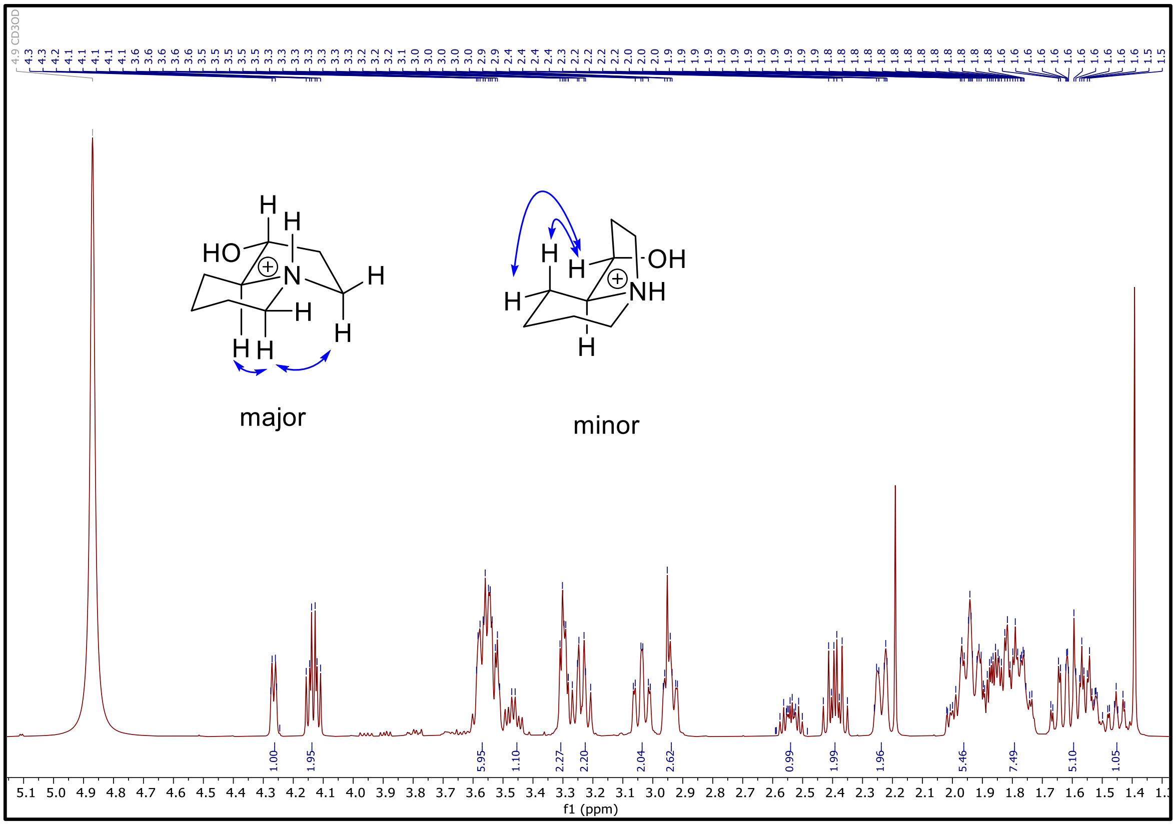

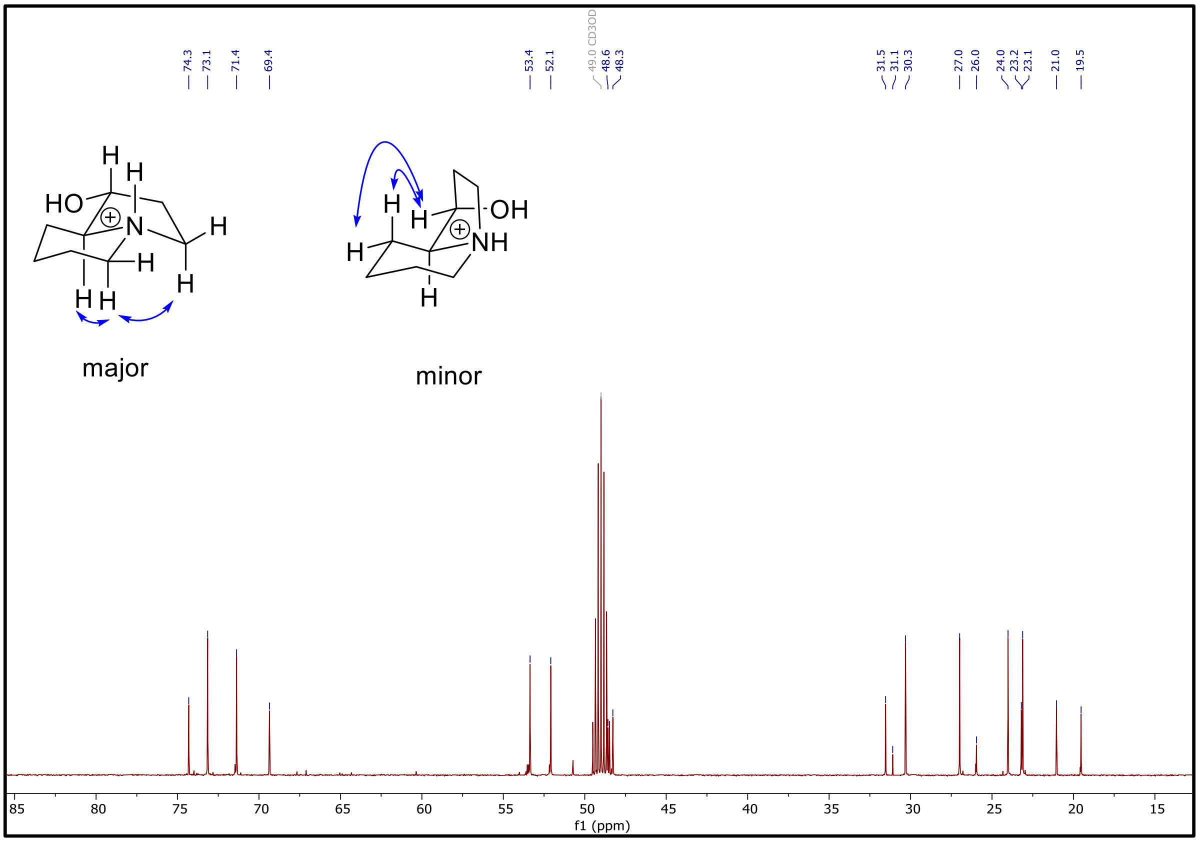

**Figure S32. [1]H NMR (500 MHz) and [13]C NMR (126 MHz) of 1•HCl in $CD_3OD$.**

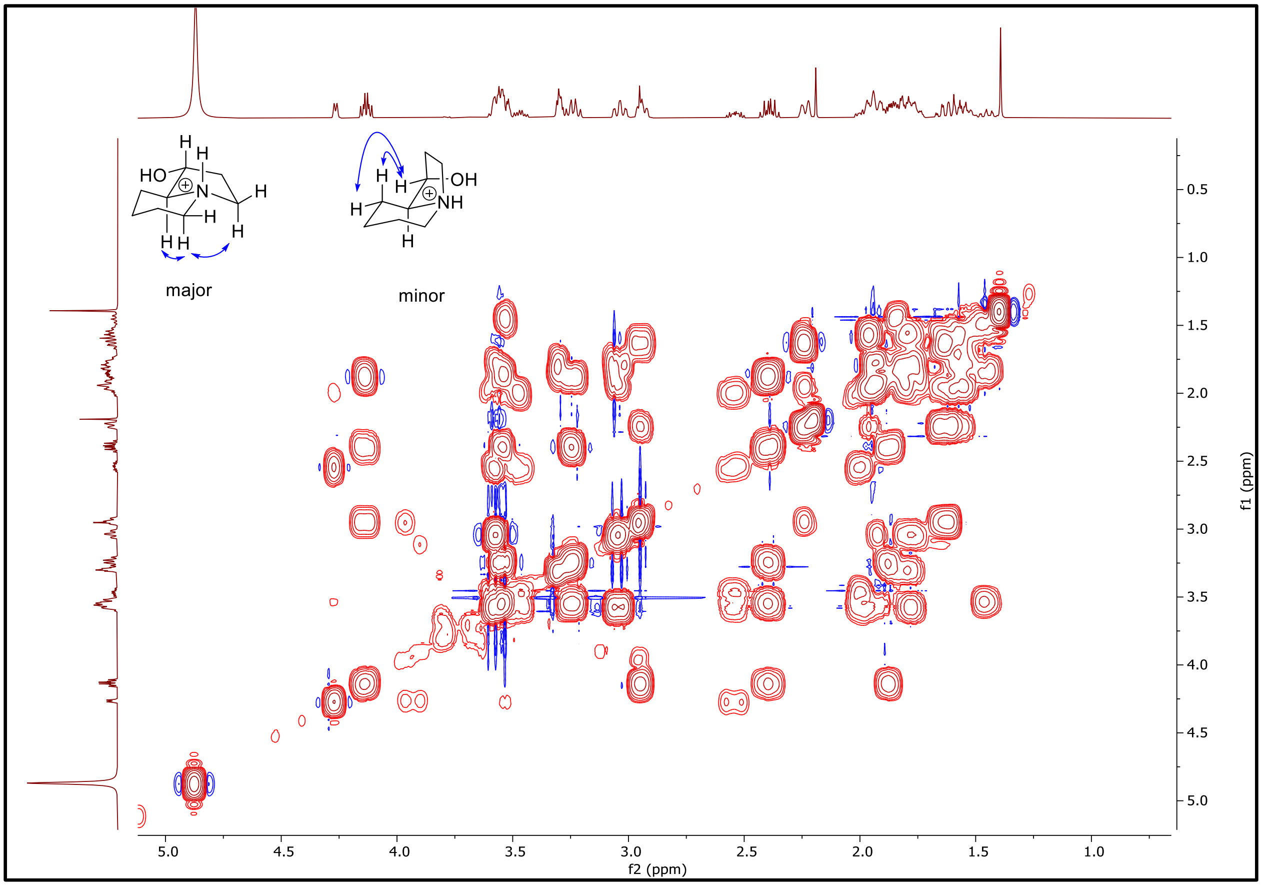

**Figure S33. $^1H$-$^1H$ COSY NMR spectrum (500 MHz) of 1•HCl in $CD_3OD$.**

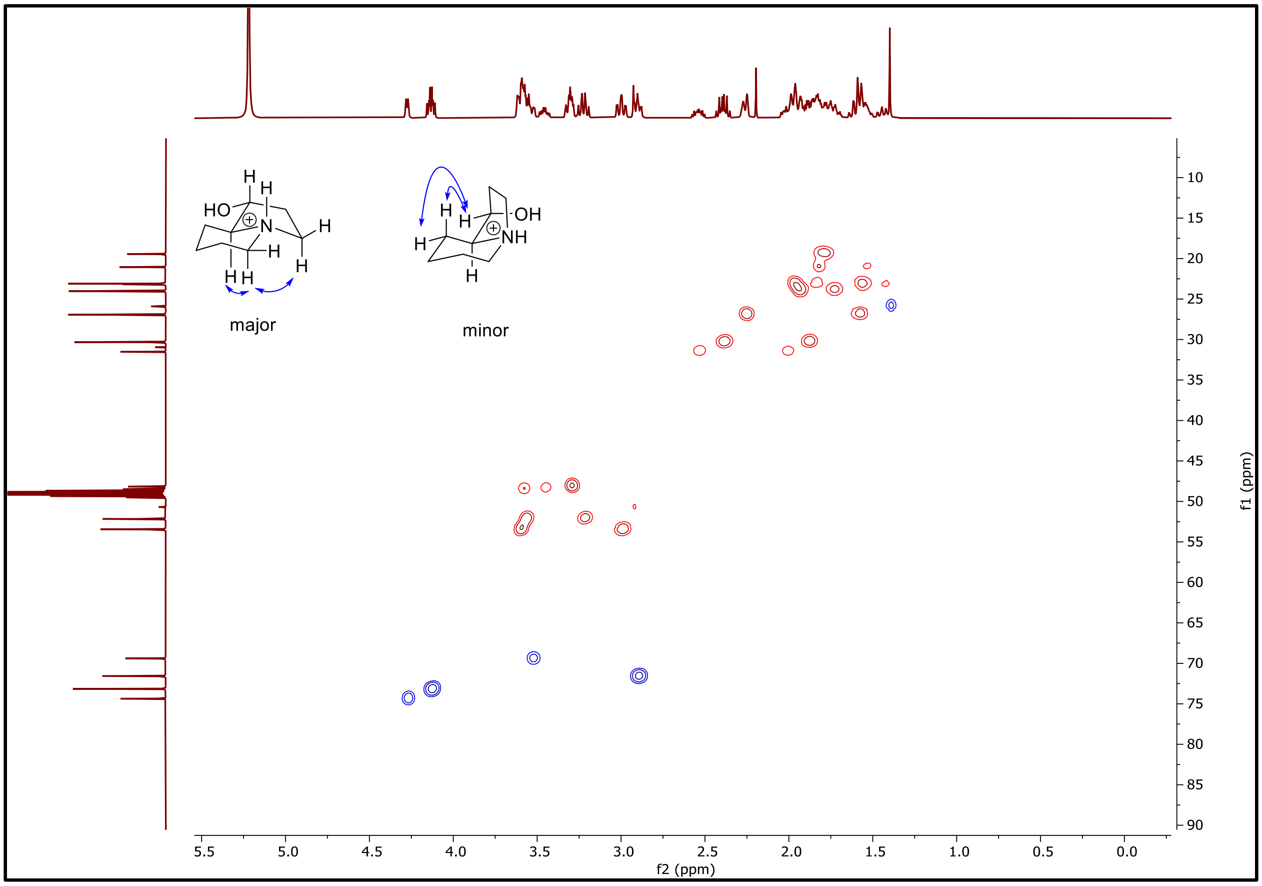

**Figure S34. $^{1}$H-$^{13}$C HSQC NMR (500 MHz) of 1•HCl in $CD_3OD$.**

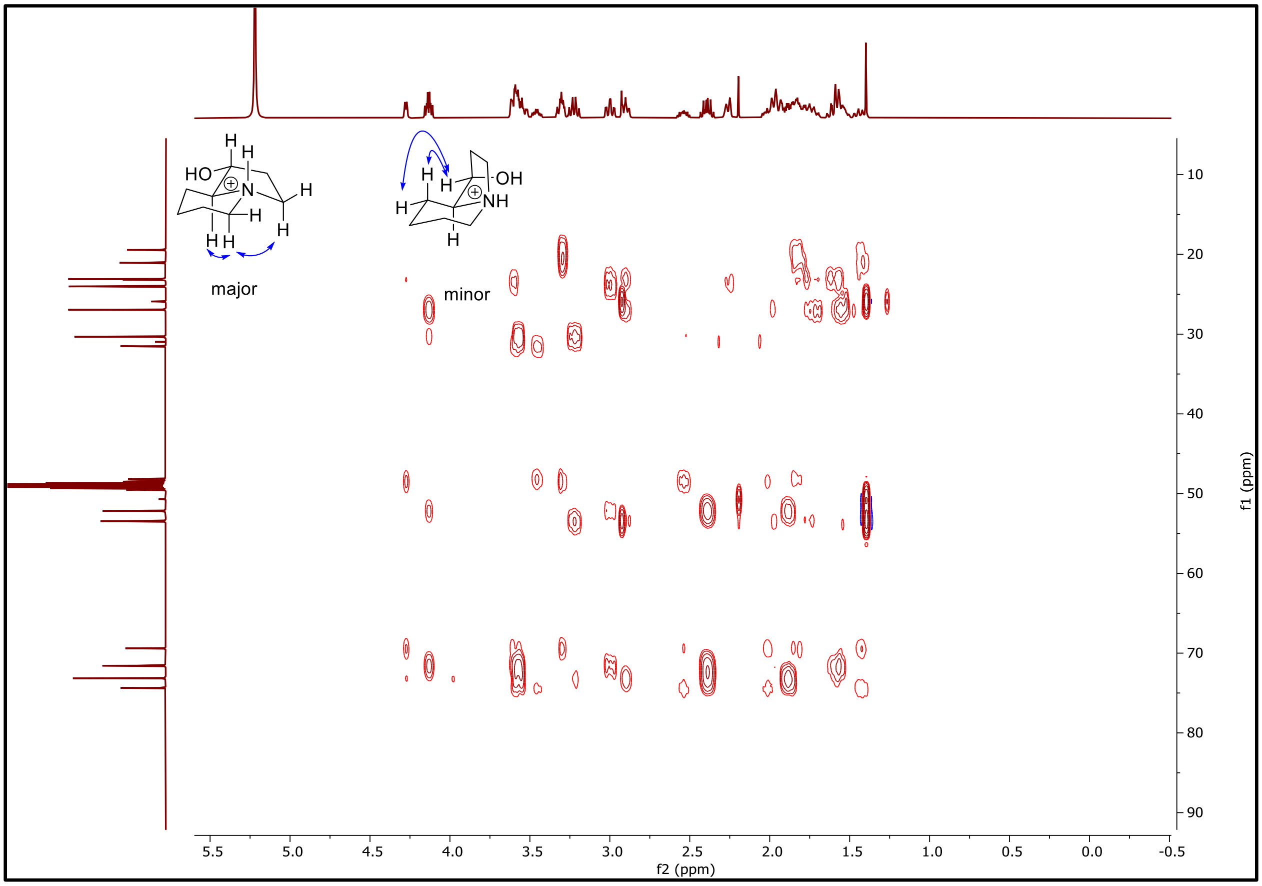

**Figure S35. $^{1}H$-$^{13}C$ HMBC NMR (500 MHz) of 1•HCl in $CD_3OD$.**

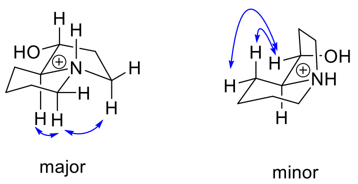

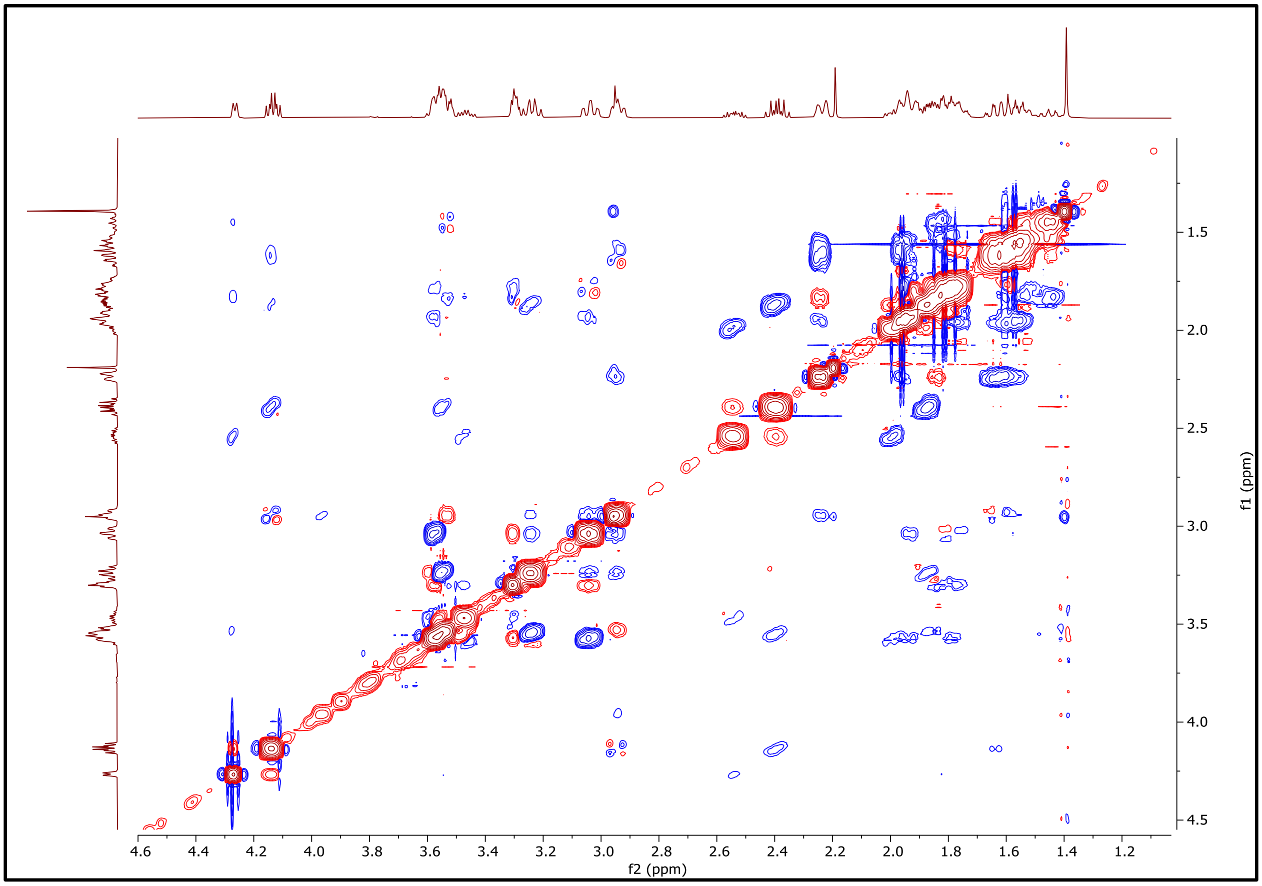

**Figure S36. [1]H-[1]H NOESY NMR (500 MHz) of 1•HCl in CD3OD.**

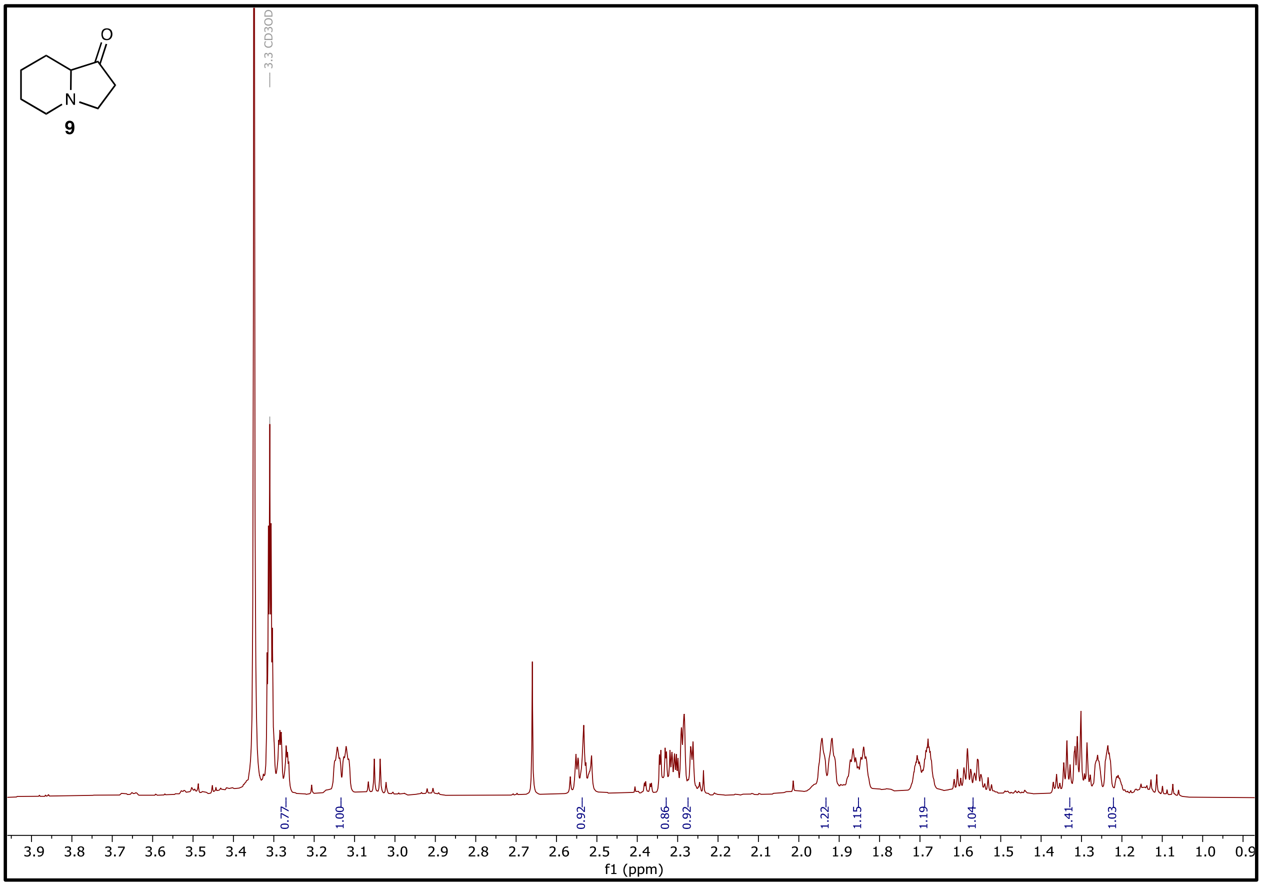

**Figure S37. $^{1}H$ NMR (500 MHz) of 9 in $CD_3OD$.**

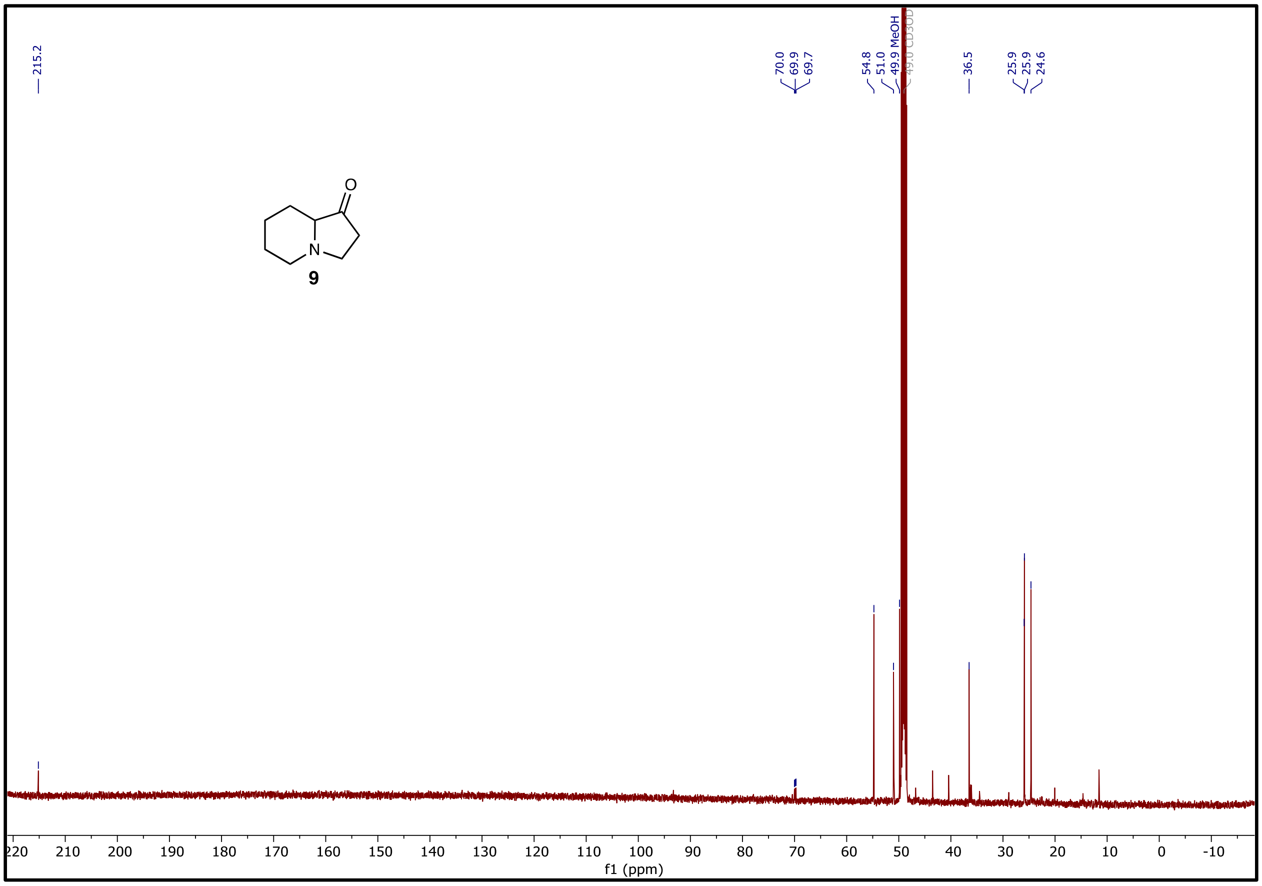

**Figure S38. $^{13}C$ NMR spectrum (126 MHz) of 9 in $CD_3OD$.**

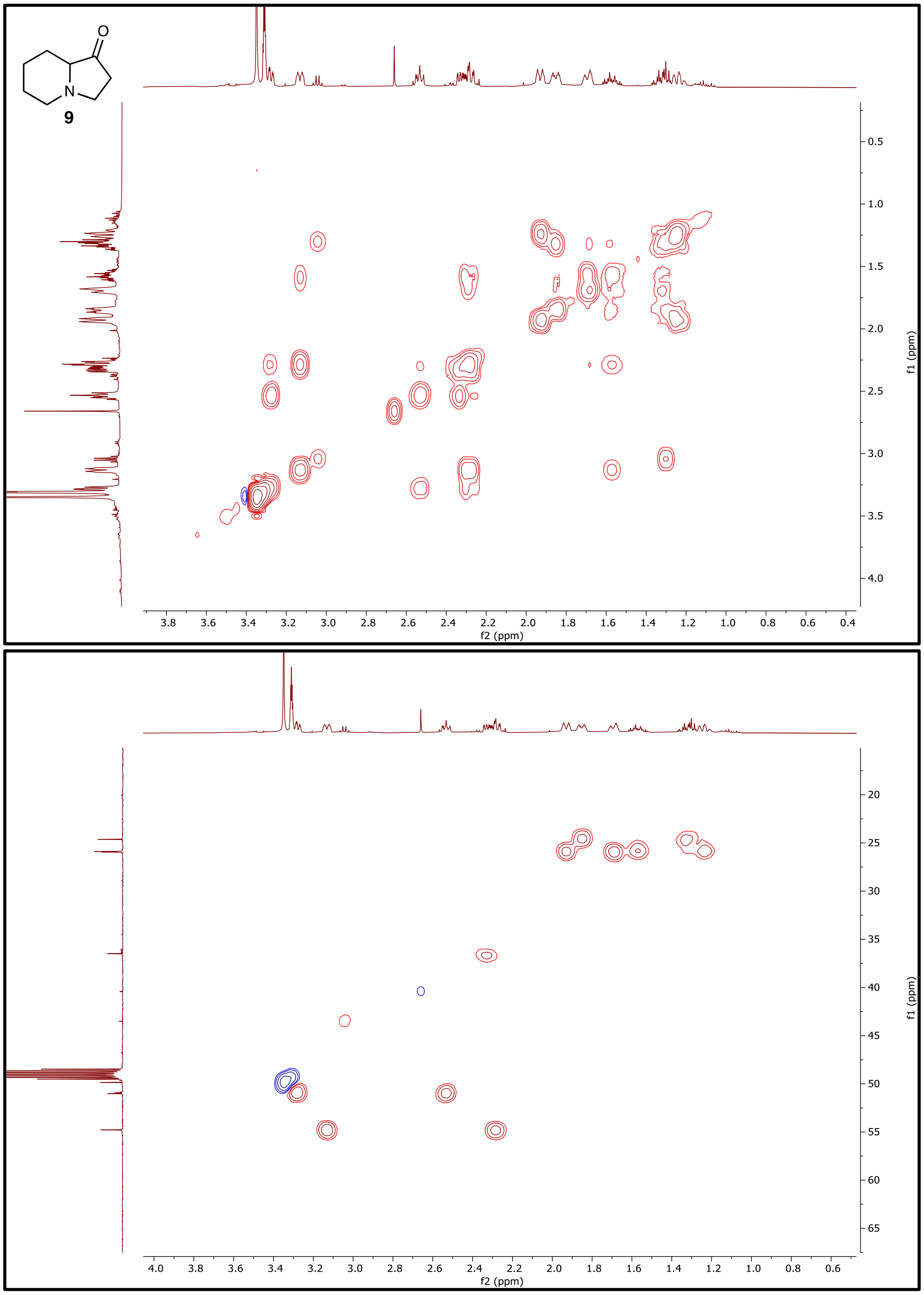

**Figure S39. $^{1}H$-$^{1}H$ COSY and $^{1}H$-$^{13}C$ HSQC NMR (500 MHz) spectra of 9 in $CD_3OD$.**

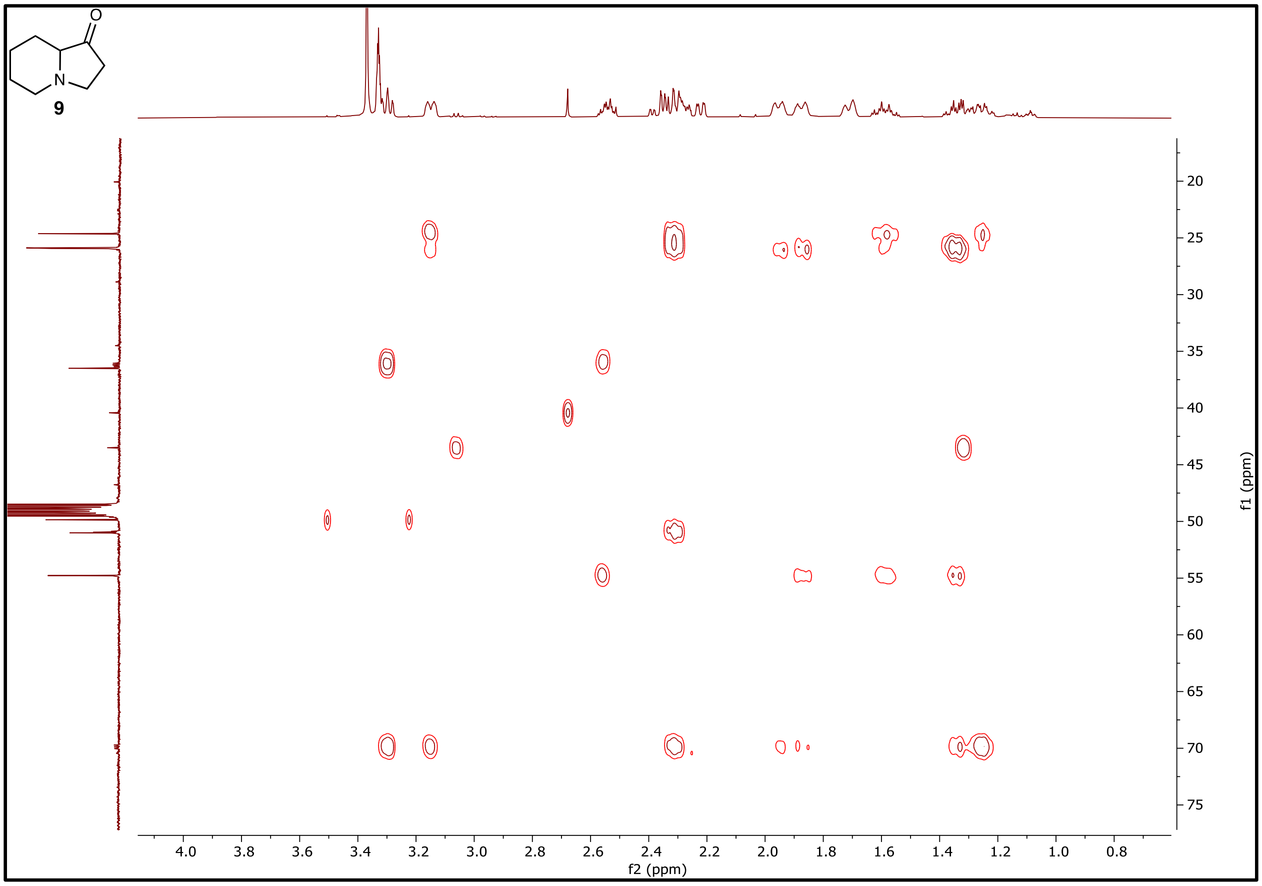

**Figure S40.** $^{1}H$-$^{13}C$ **HMBC NMR (500 MHz) of 9 in** $CD_3OD$**.**

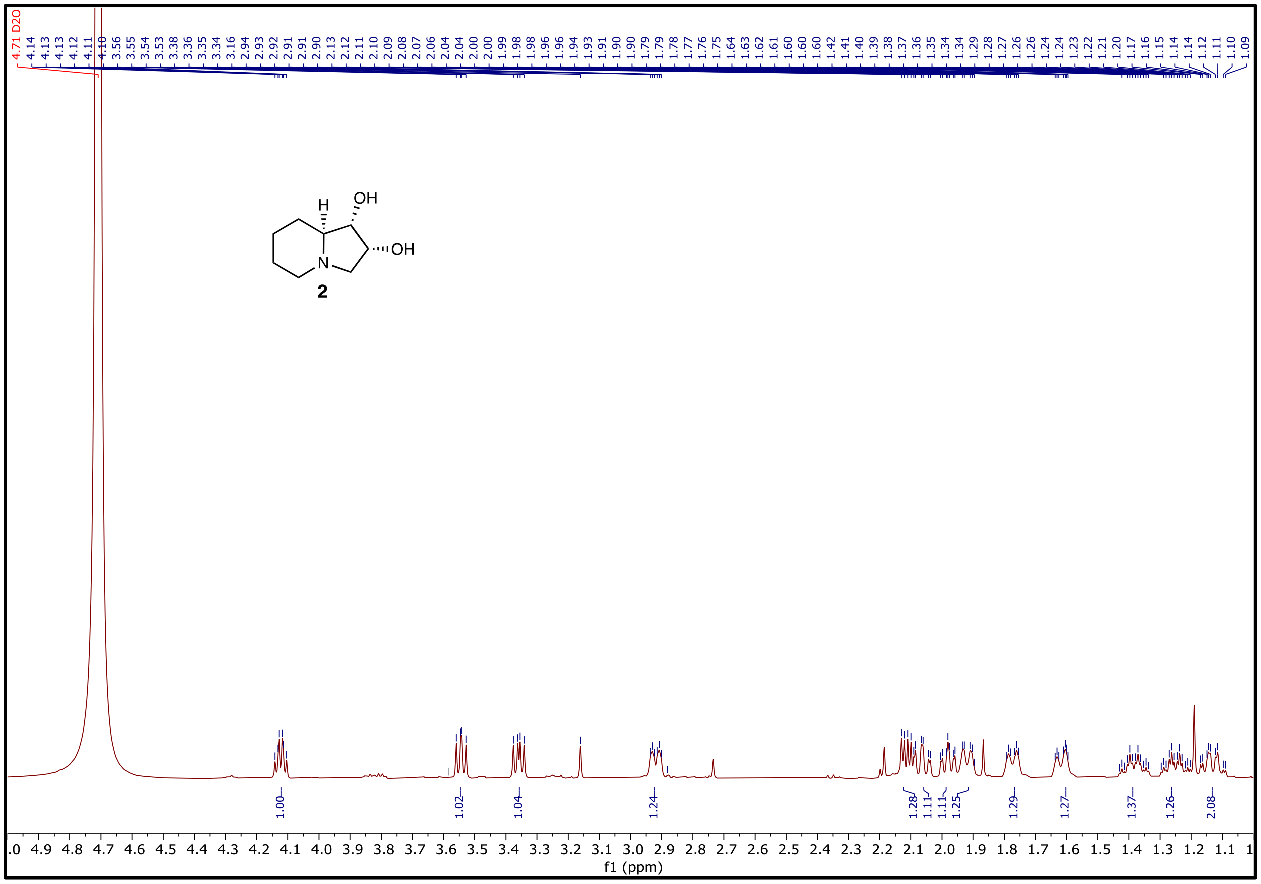

**Figure S41. $^1$H NMR (500 MHz) and of 2 in $D_2O$.**

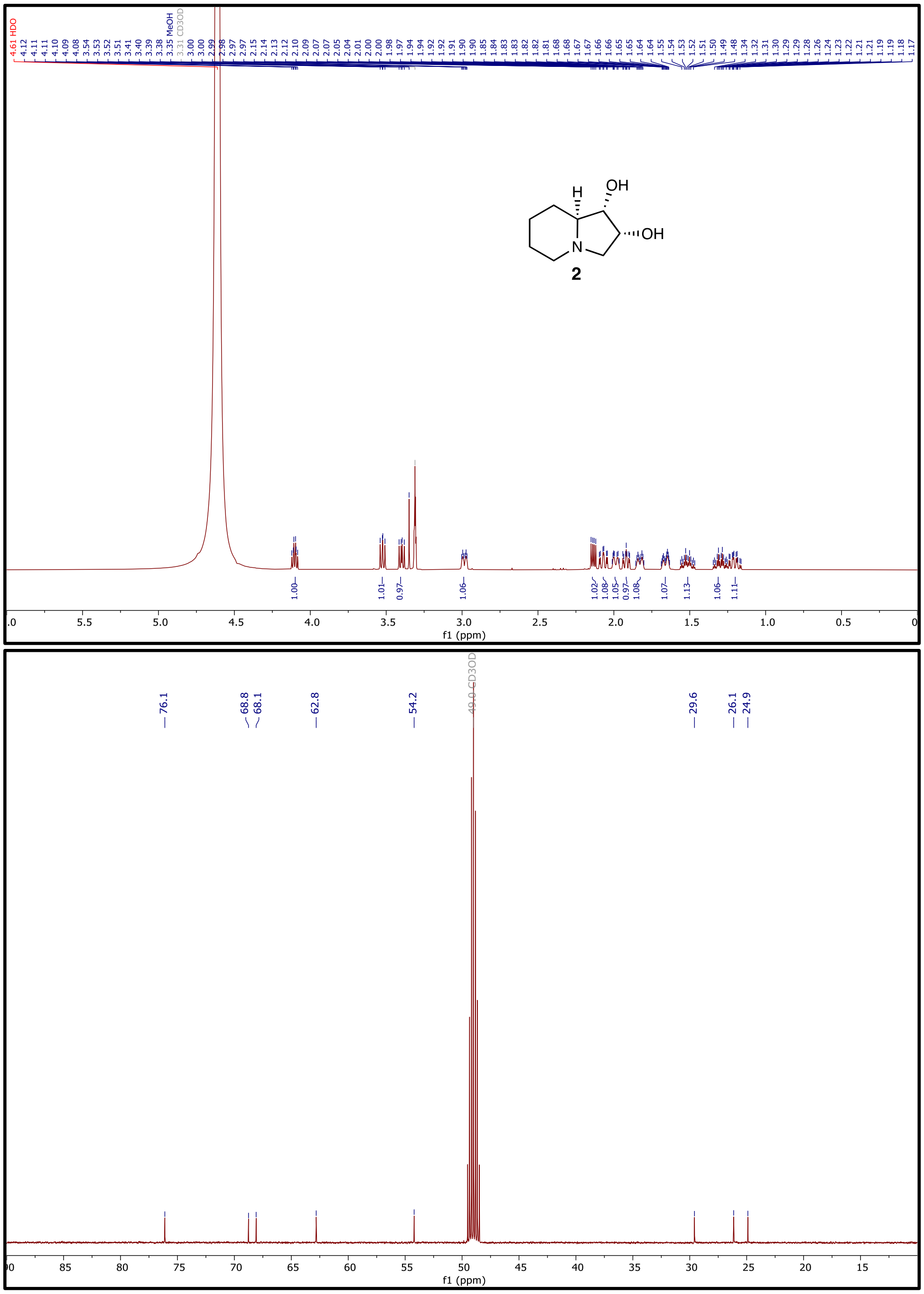

**Figure S42. $^1$H NMR (500 MHz) and $^{13}$C NMR (126 MHz) spectra of 2 in $CD_3OD$.**

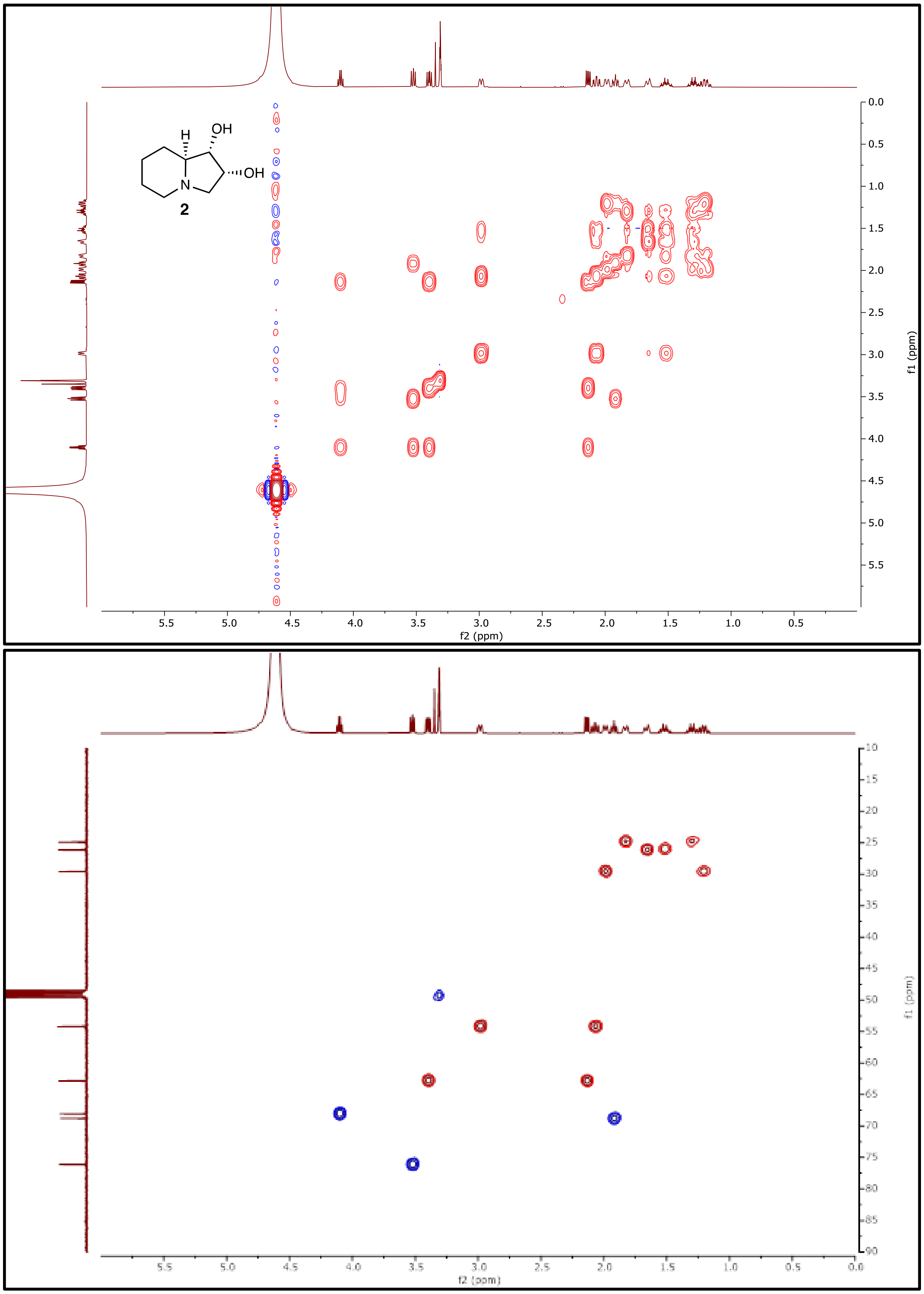

**Figure S43.** $^{1}H$-$^{1}H$ **COSY NMR (500 MHz) and** $^{1}H$-$^{13}C$ **HSQC NMR (500 MHz) of 2 in** $CD_3OD$**.**

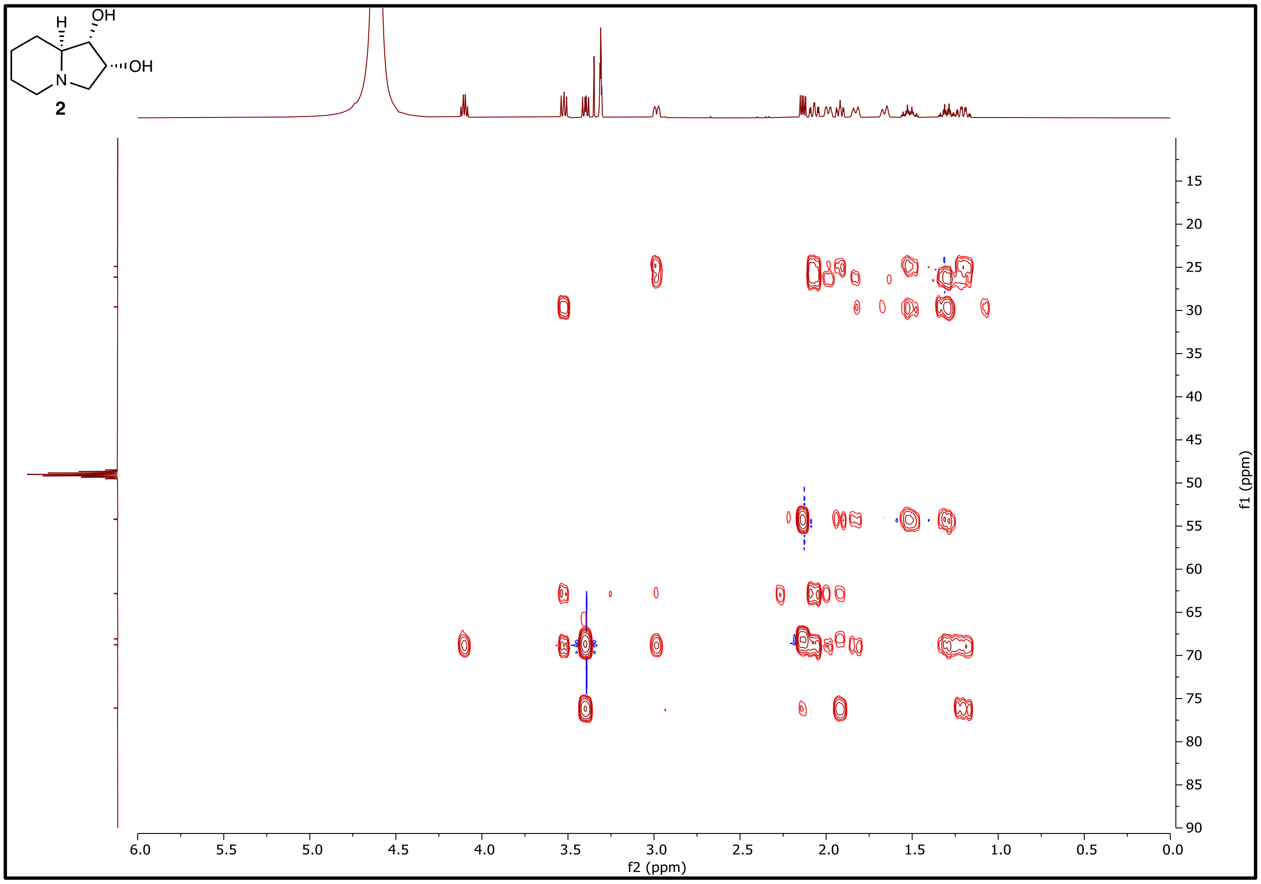

**Figure S44. $^1$H-$^{13}$C HMBC NMR (500 MHz) of 2 in $CD_3OD$.**

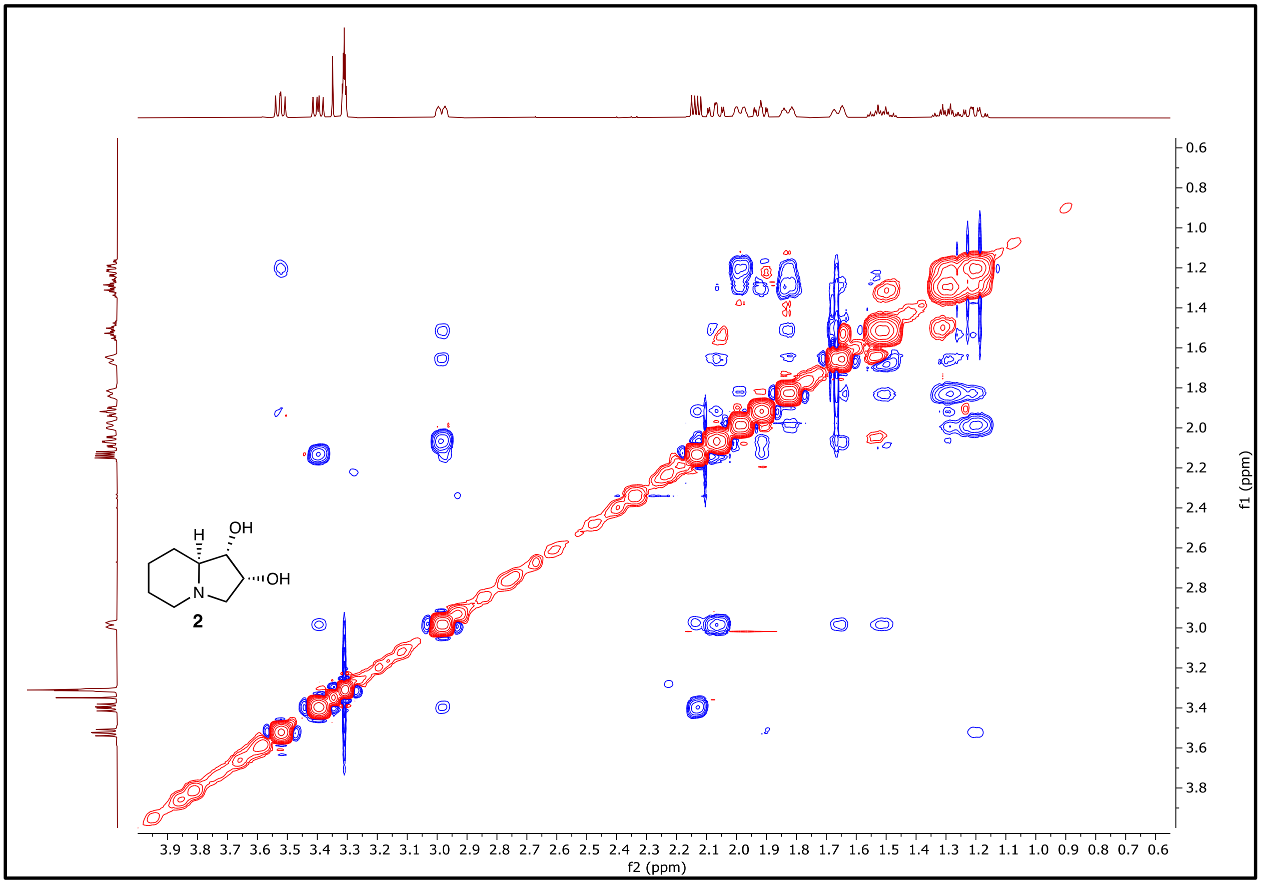

**Figure S45. $^1H$-$^1H$ NOESY NMR (500 MHz) of 2 in $CD_3OD$.**

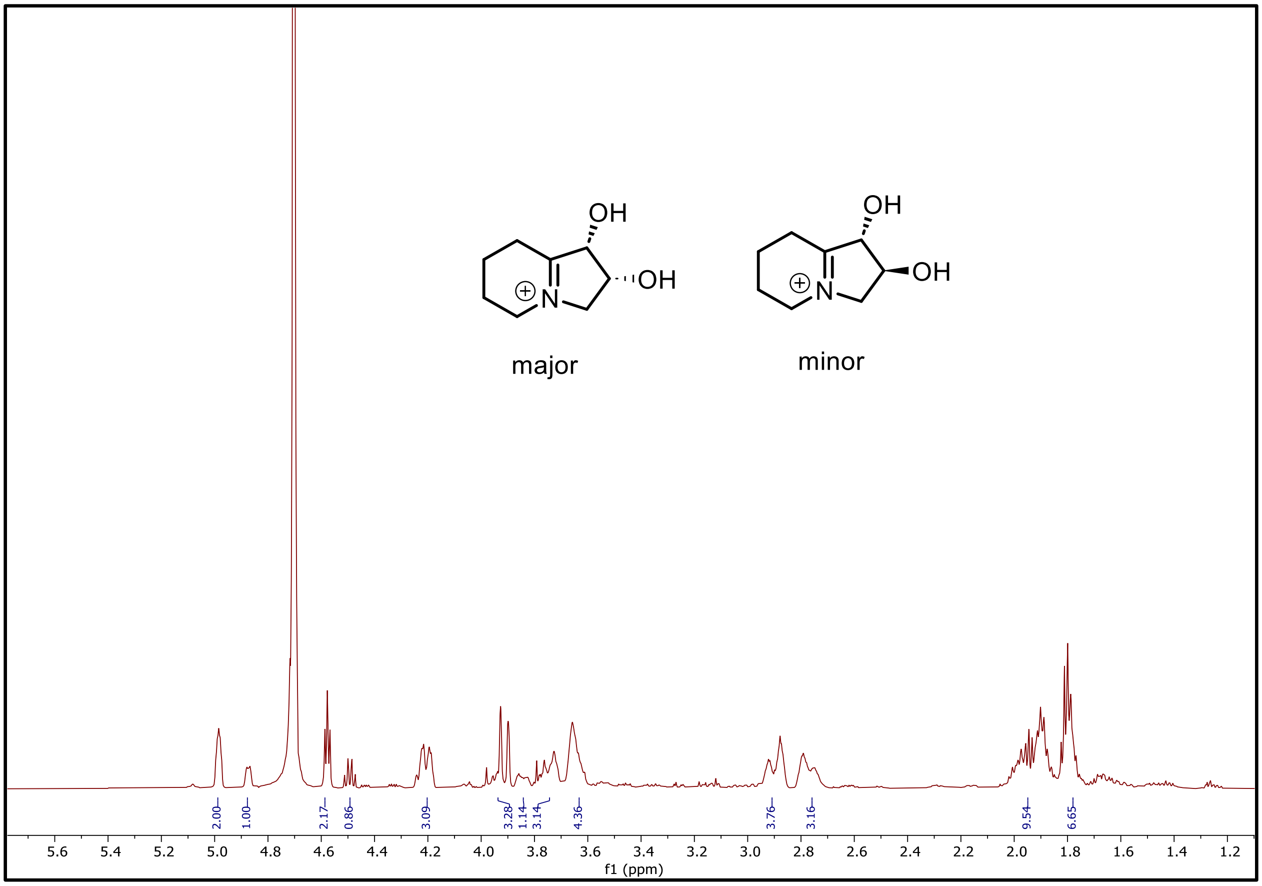

**Figure S46. [1]H NMR (500 MHz) of 6 and 2-*epi*-6 in $D_2O$.**

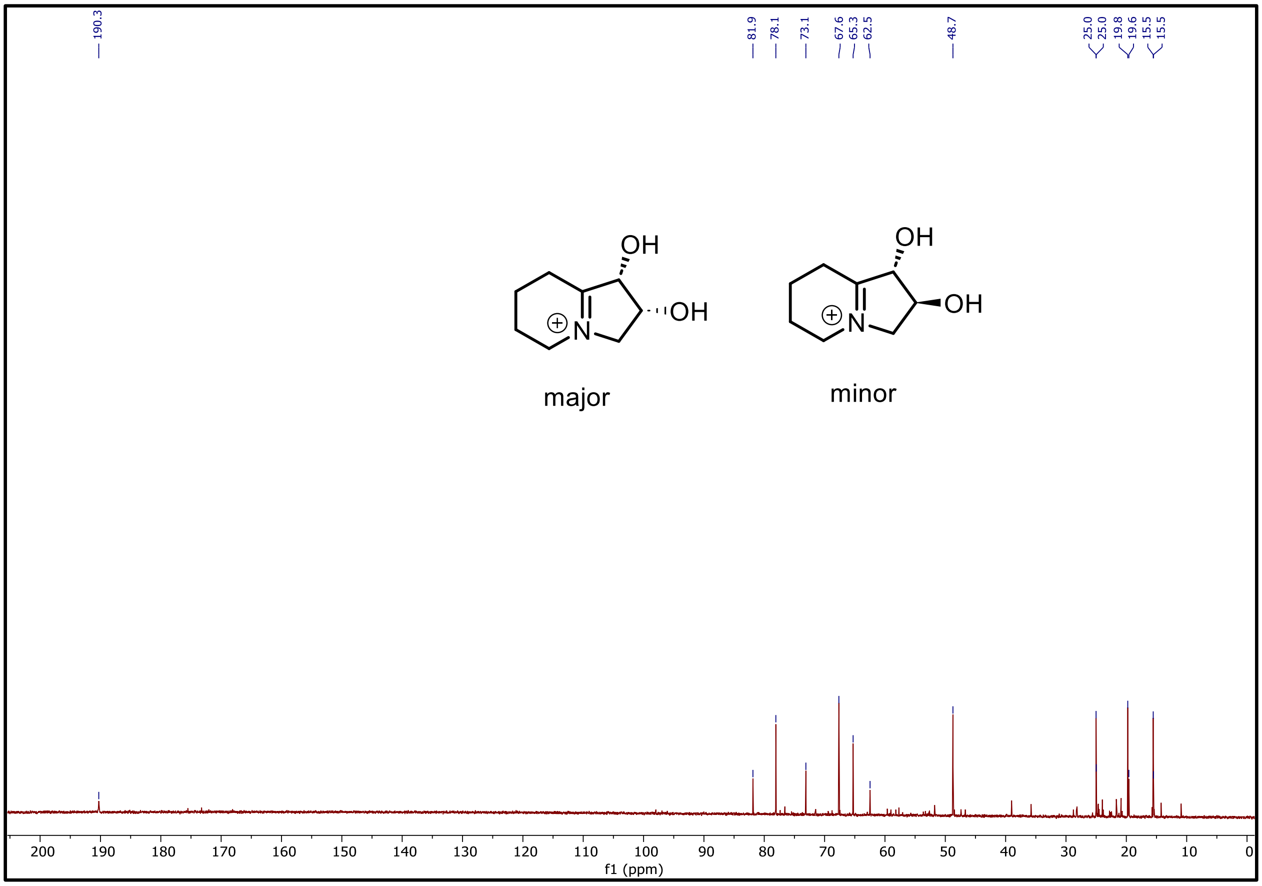

**Figure S47. $^{13}C$ NMR spectrum (126 MHz) of 6 and 2-*epi*-6 in $D_2O$.**

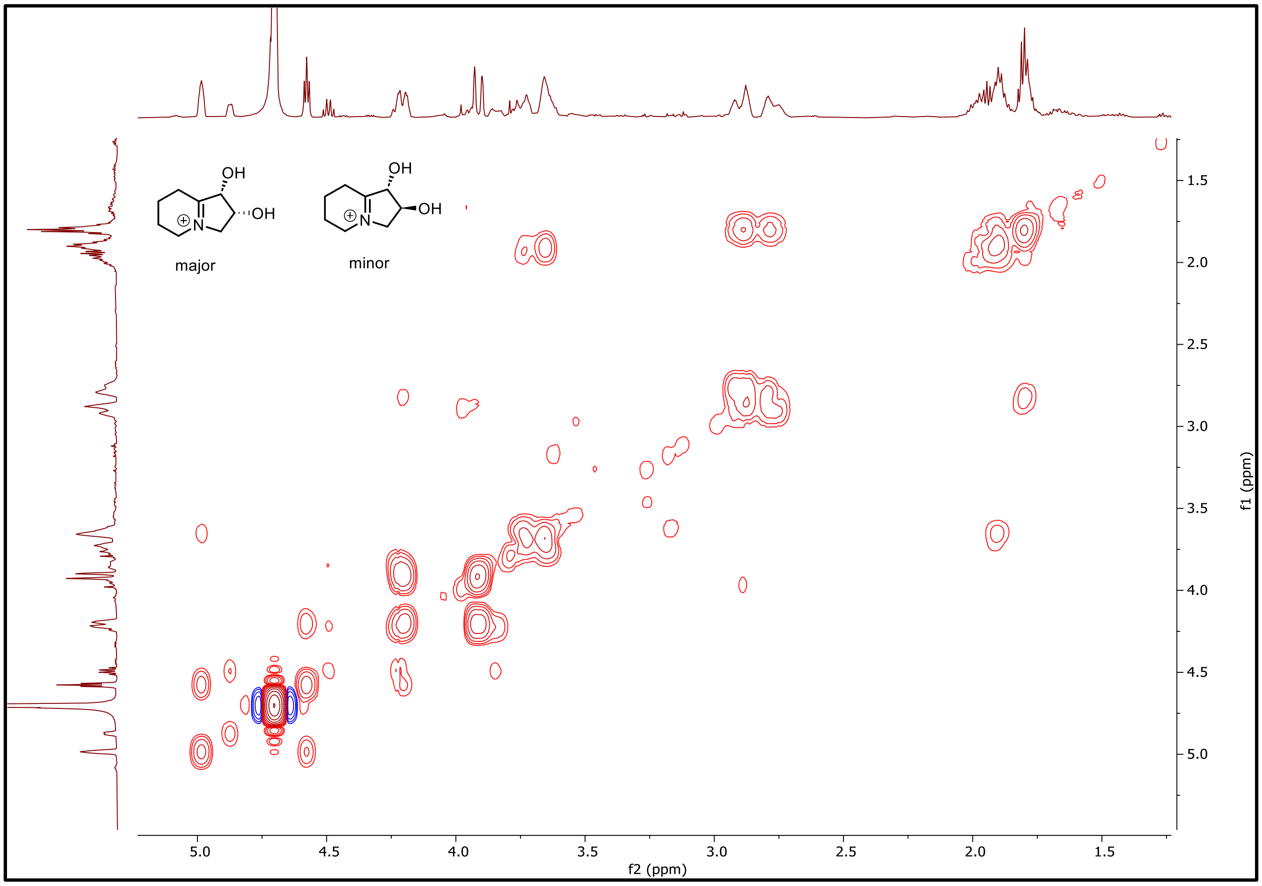

**Figure S48.** $^1$**H-**$^1$**H COSY NMR spectrum (500 MHz) of 6 and 2-*epi*-6 in D**$_2$**O.**

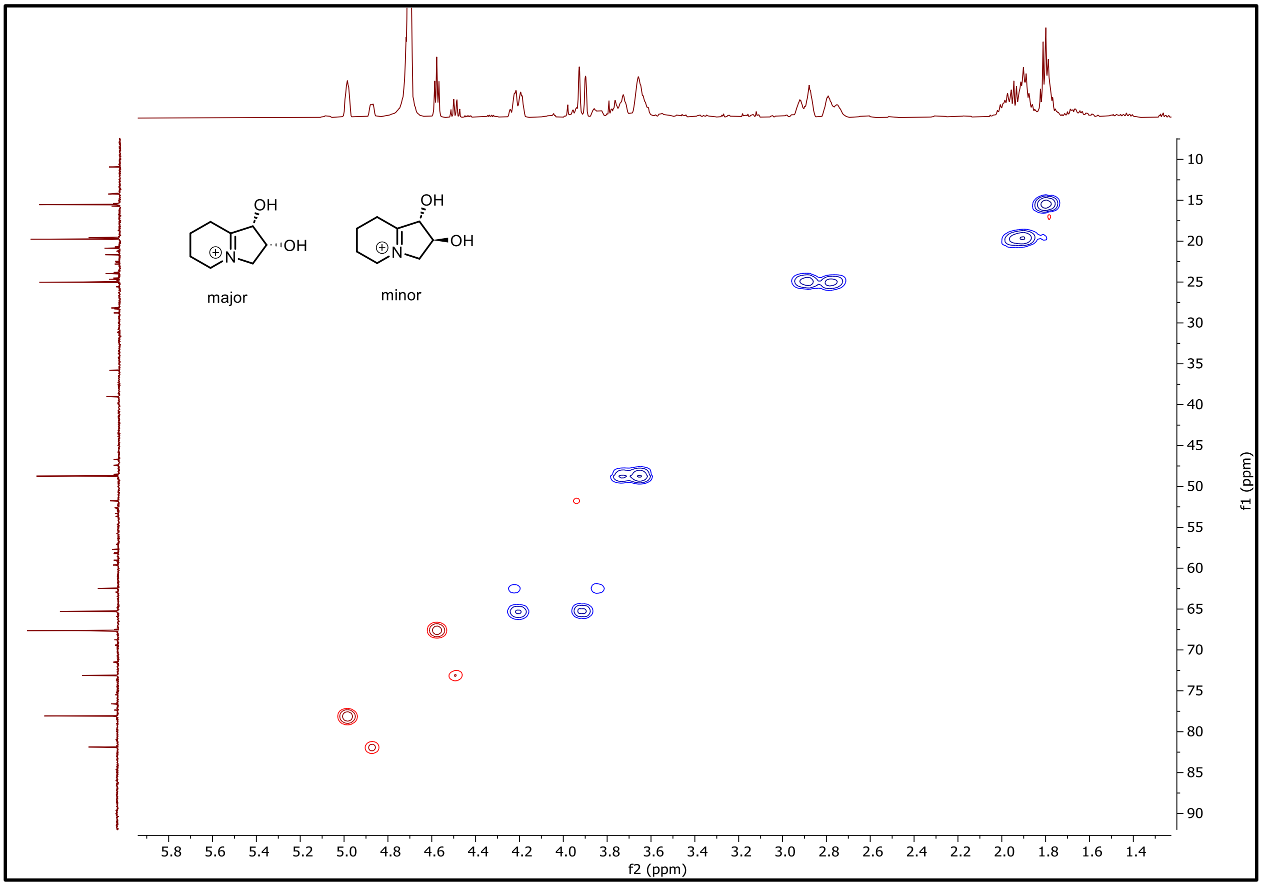

**Figure S49. $^{1}H$-$^{13}C$ HSQC NMR (500 MHz) of 6 and 2-*epi*-6 in $D_2O$.**

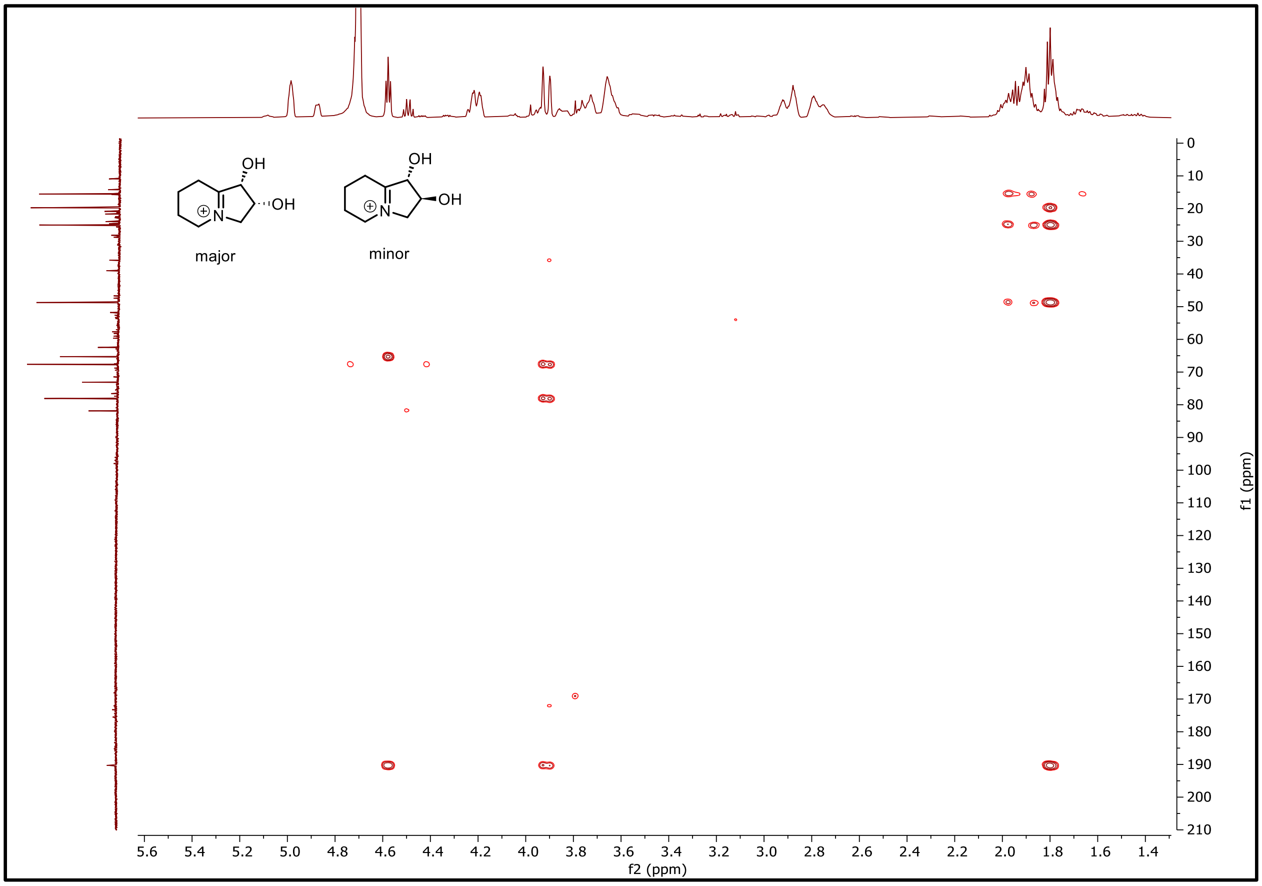

**Figure S50. $^{1}H$-$^{13}C$ HMBC NMR (500 MHz) of 6 and 2-*epi*-6 in $D_2O$.**

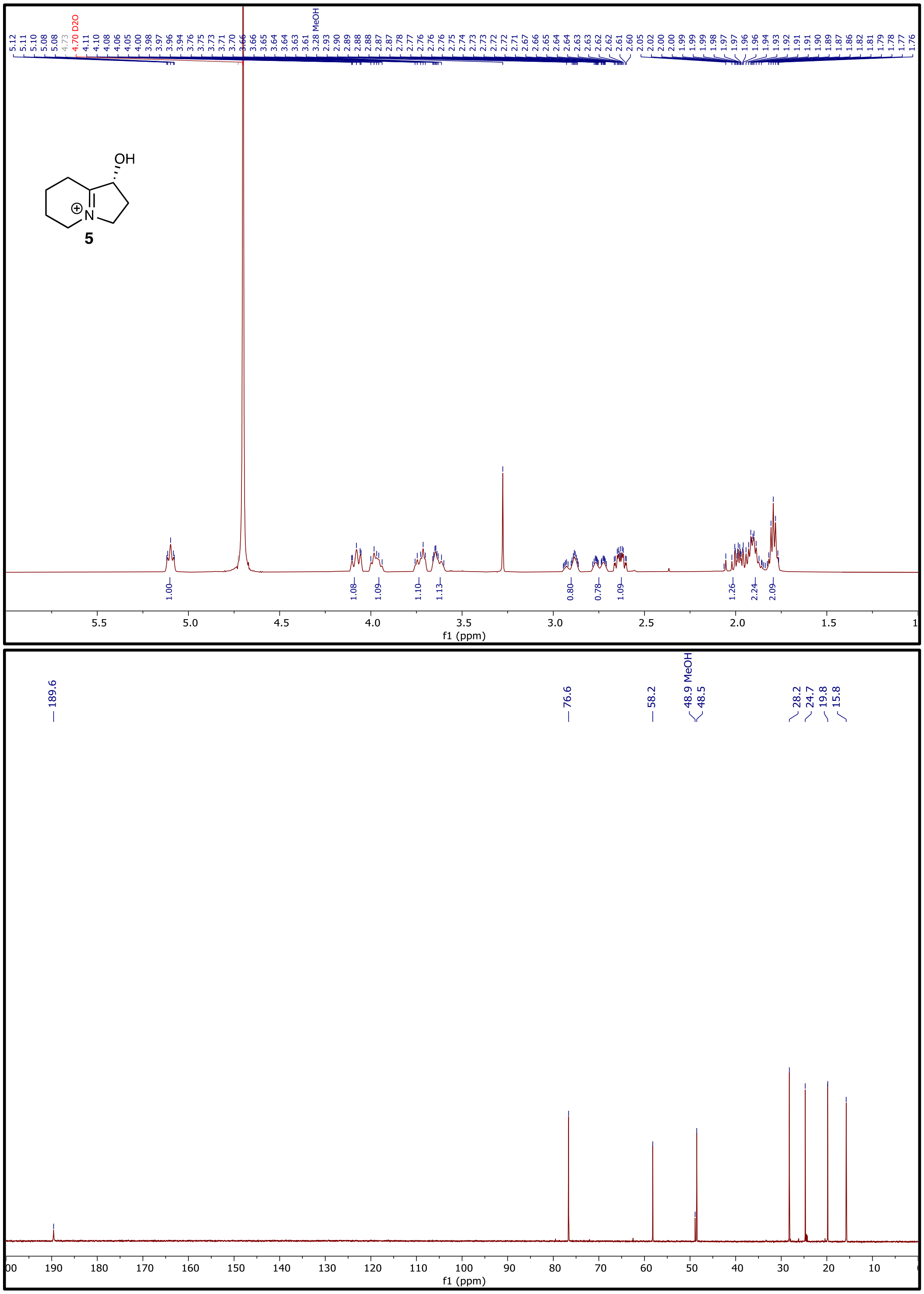

**Figure S51. $^{1}H$ NMR (500 MHz) and $^{13}C$ NMR spectra of 5 in $D_2O$.**

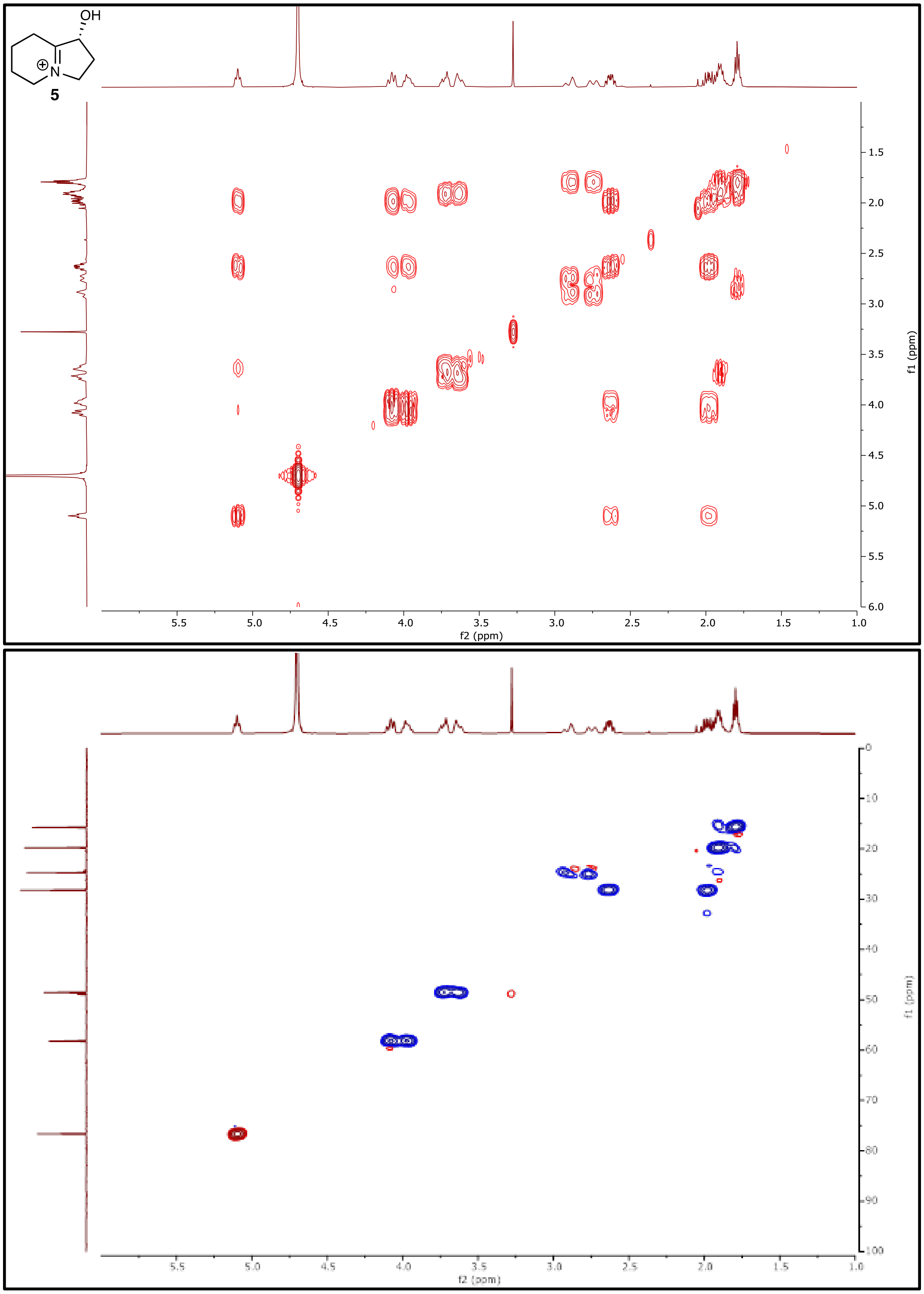

**Figure S52.** **$^1H$-$^1H$ COSY NMR (500 MHz) and $^1H$-$^{13}C$ HSQC NMR (500 MHz) of 5 in $D_2O$.**

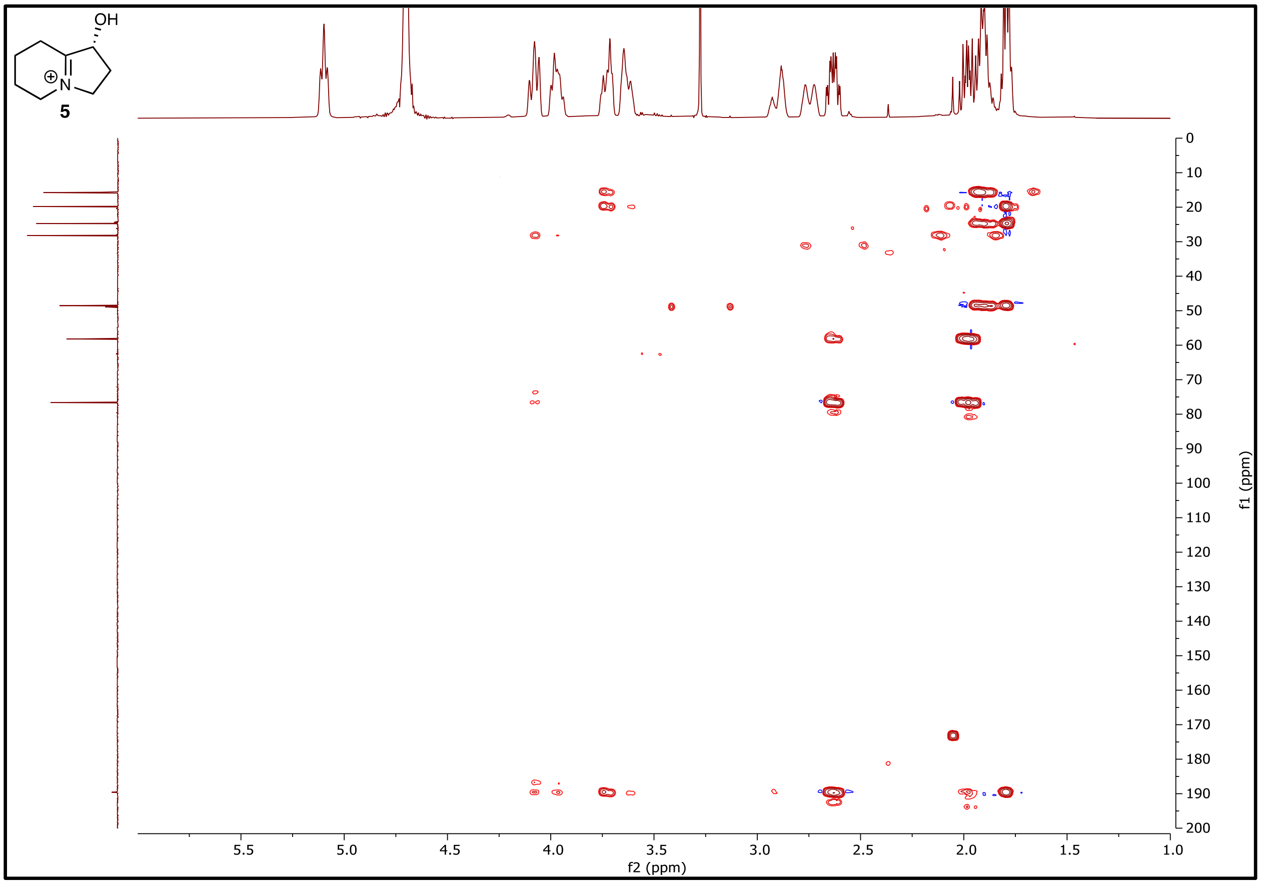

**Figure S53. $^1$H-$^{13}$C HMBC NMR (500 MHz) of 5 in $D_2O$.**

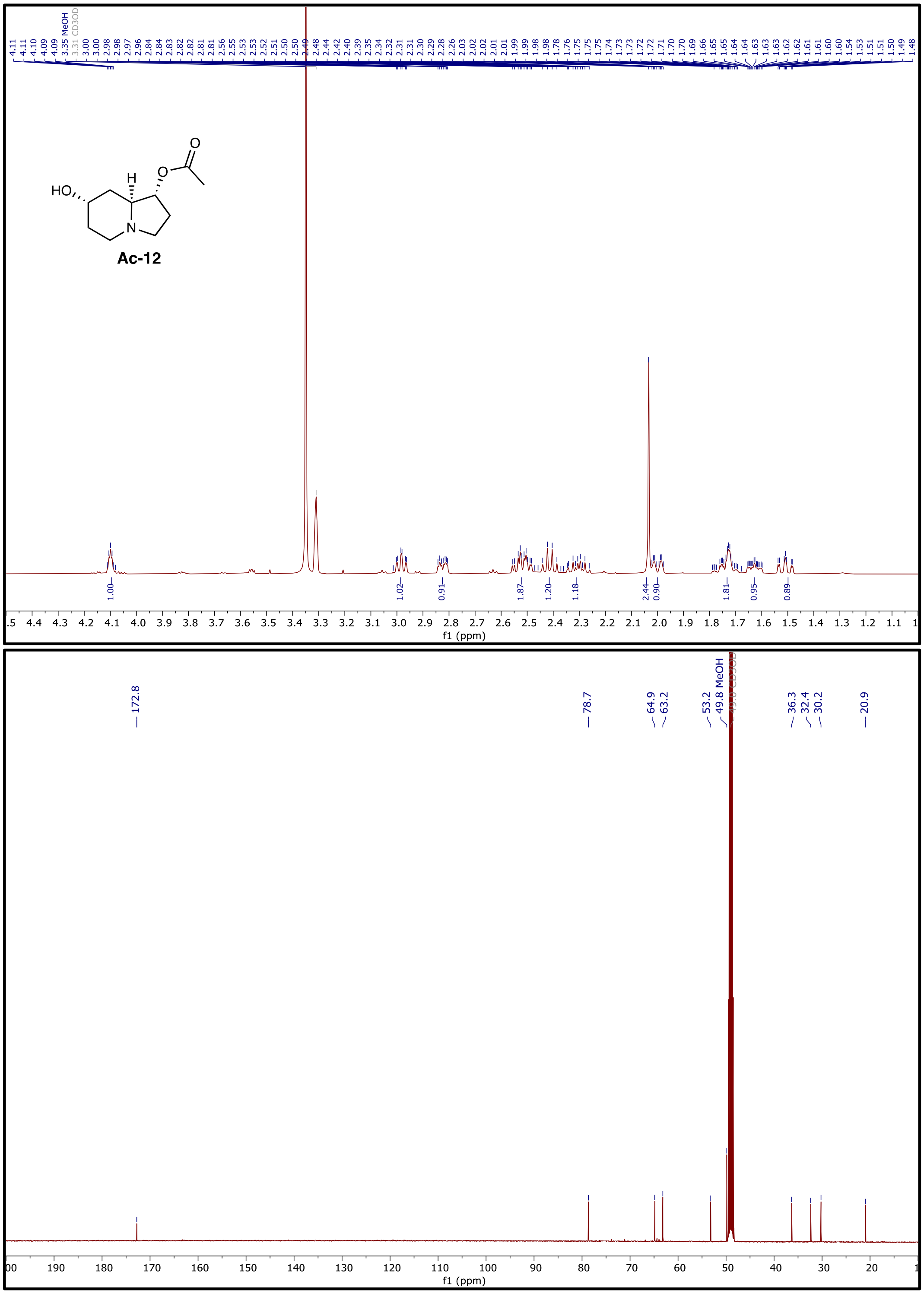

**Figure S54. $^{1}$H NMR (500 MHz) and of Ac-12 in $CD_3OD$.**

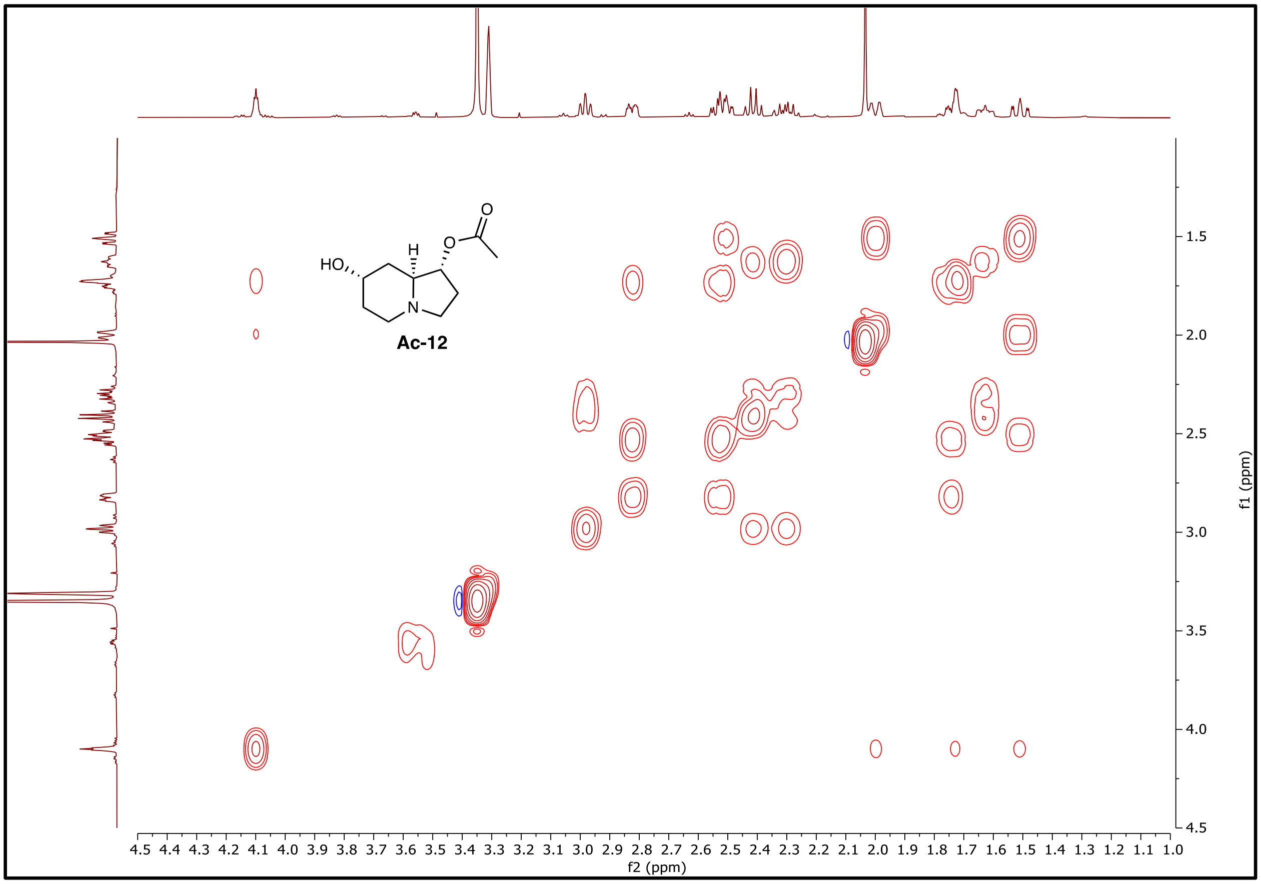

**Figure S55.** $^{1}H$-$^{1}H$ **COSY NMR spectrum (500 MHz) of Ac-12 in** $CD_3OD$**.**

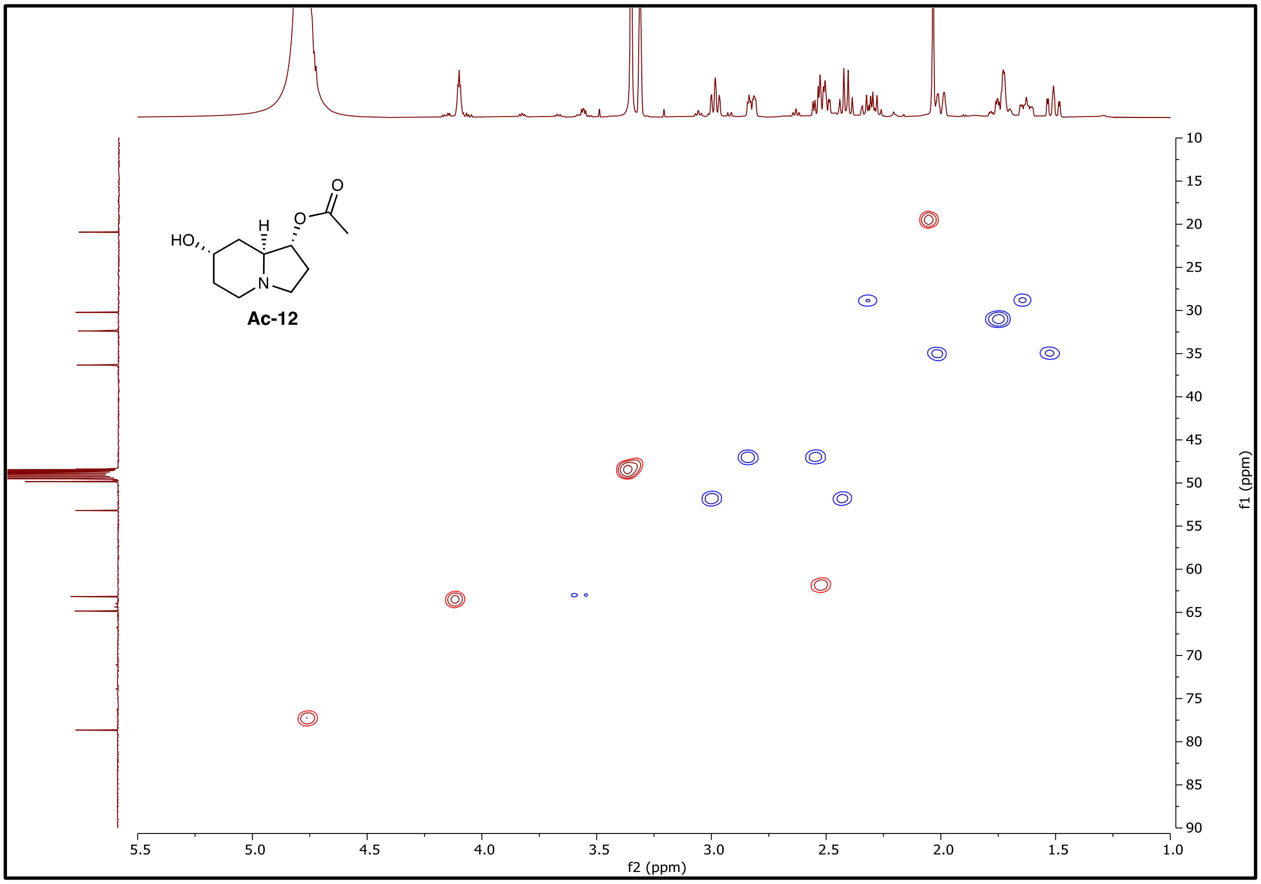

**Figure S56. $^{1}H$-$^{13}C$ HSQC NMR (500 MHz) of Ac-12 in $CD_3OD$.**

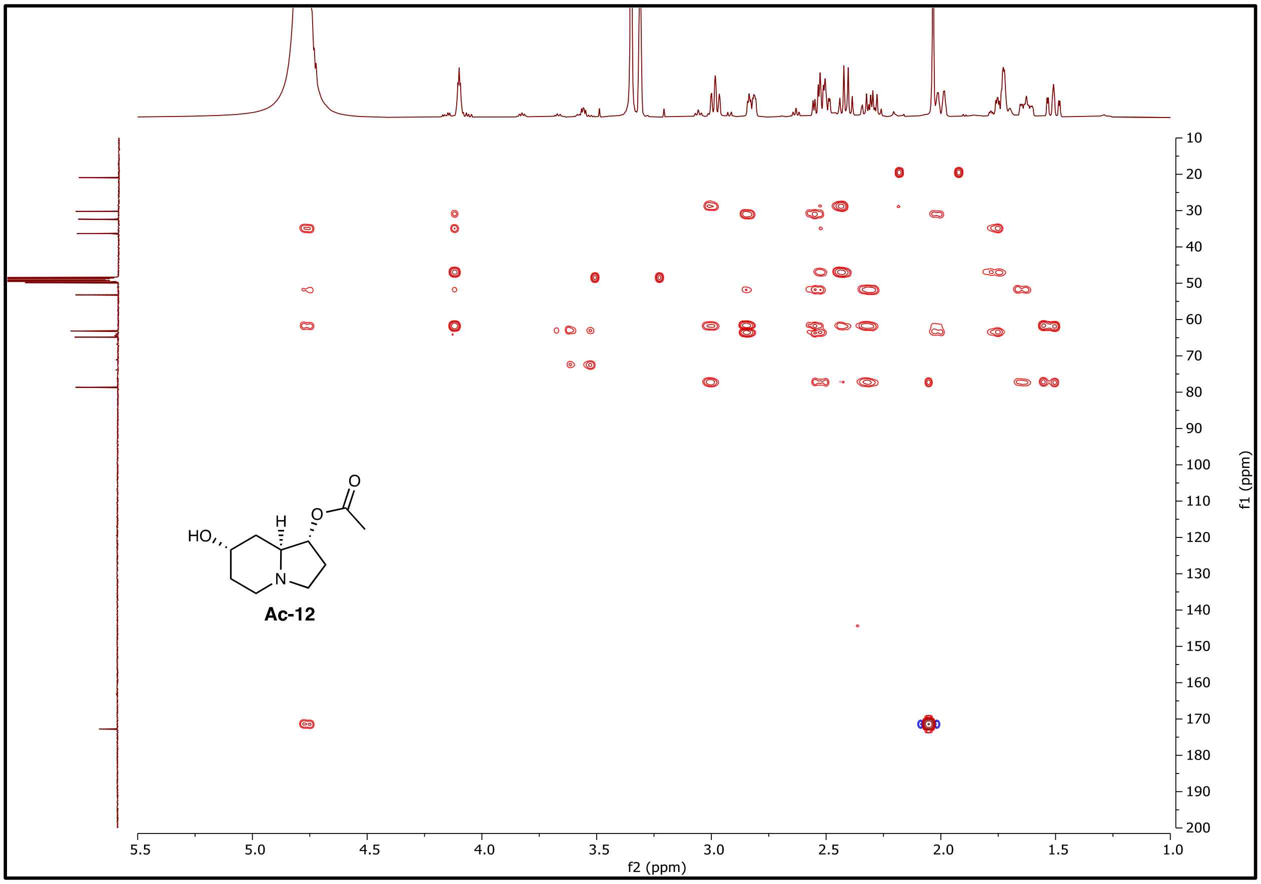

**Figure S57.** $^{1}$**H-**$^{13}$**C HMBC NMR (500 MHz) of Ac-12 in CD$_3$OD.**

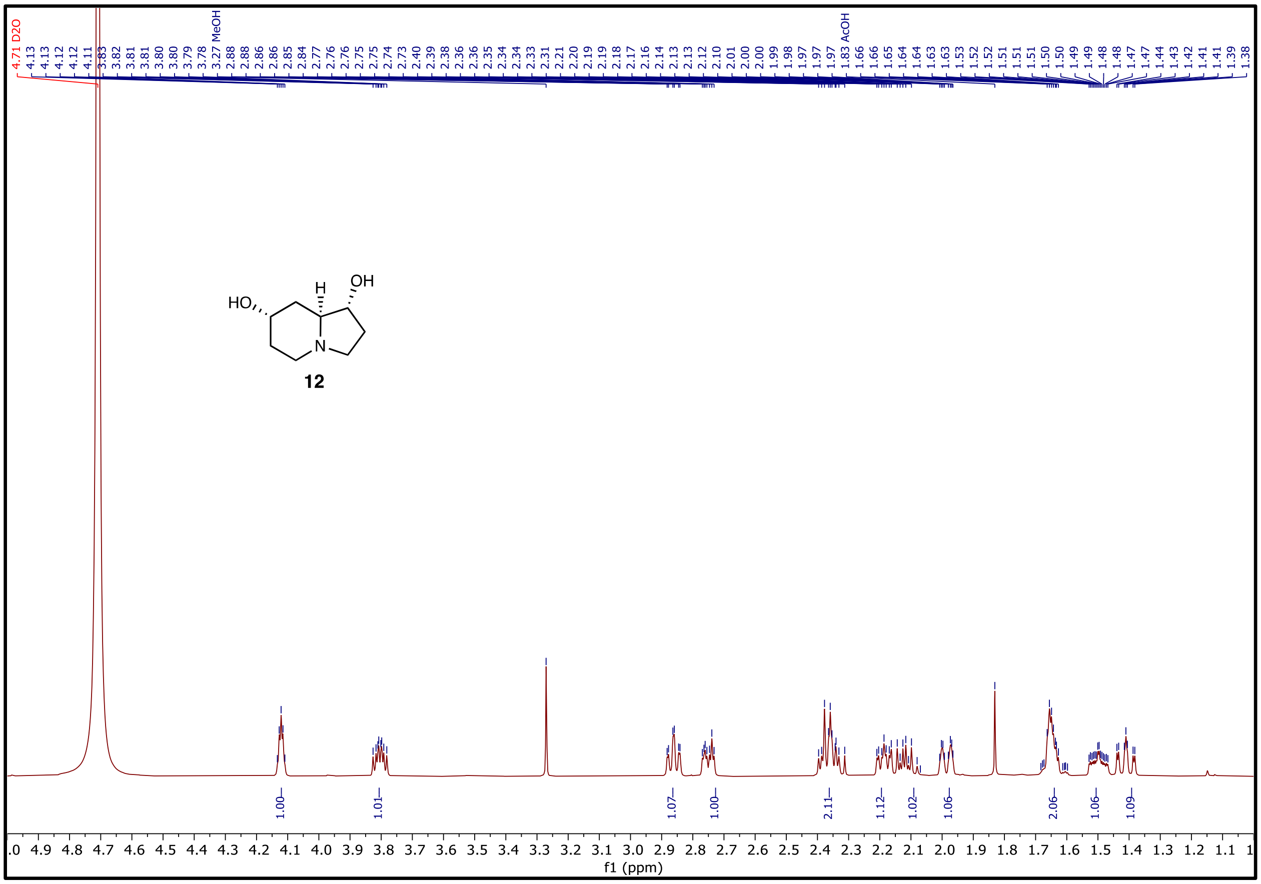

**Figure S58. $^1H$ NMR (500 MHz) of 12 in $D_2O$.**

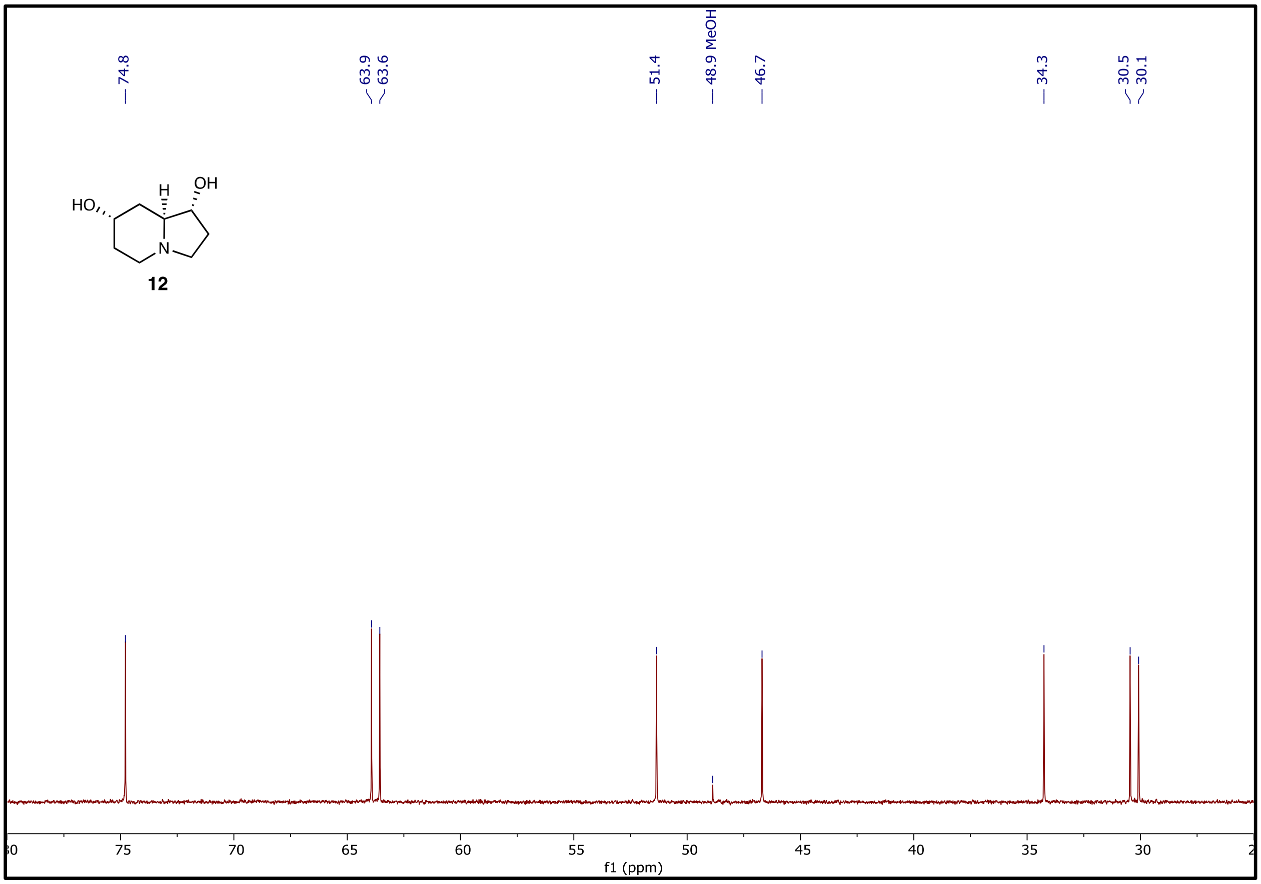

**Figure S59. $^{13}C$ NMR spectrum (126 MHz) of 12 in $D_2O$.**

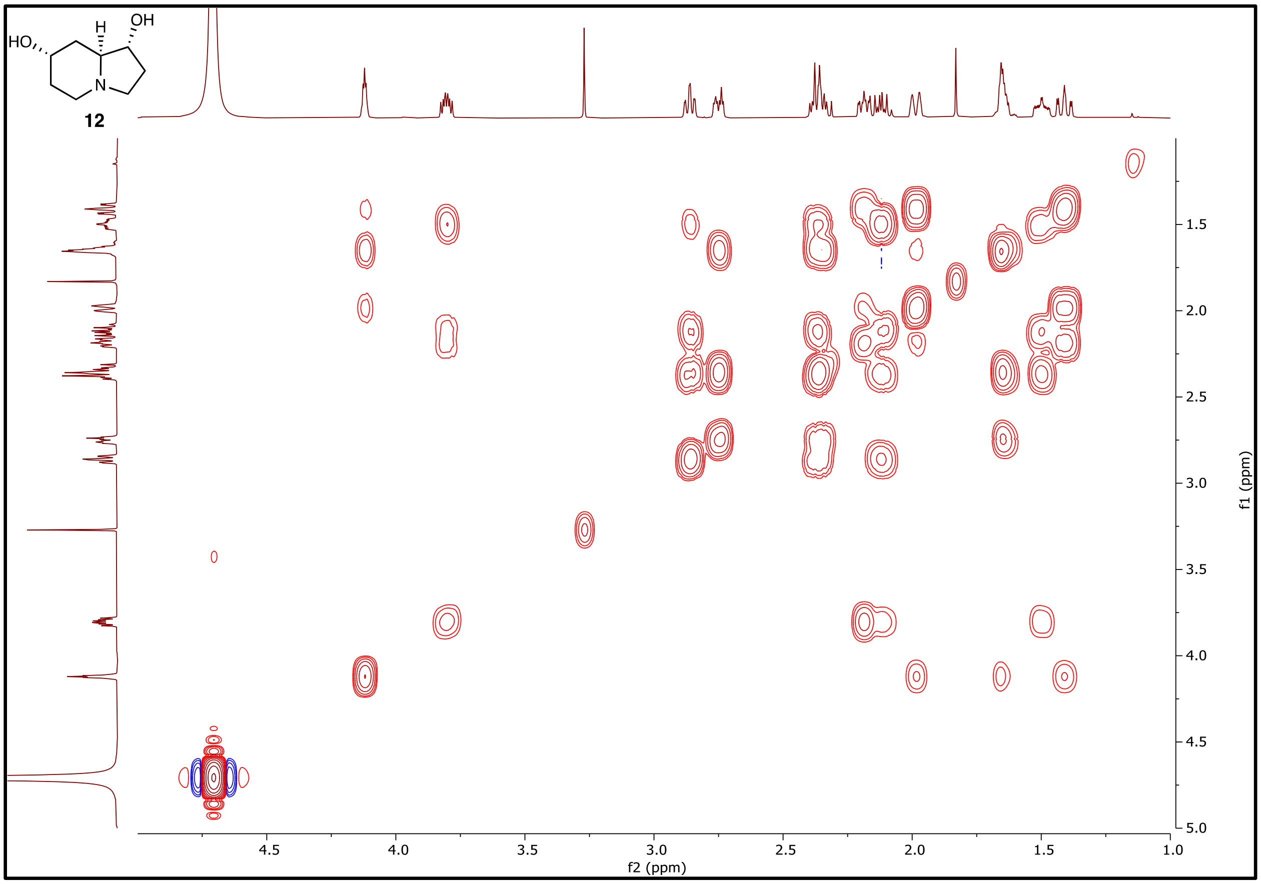

**Figure S60. $^1H$-$^1H$ COSY NMR spectrum (500 MHz) of 12 in $D_2O$.**

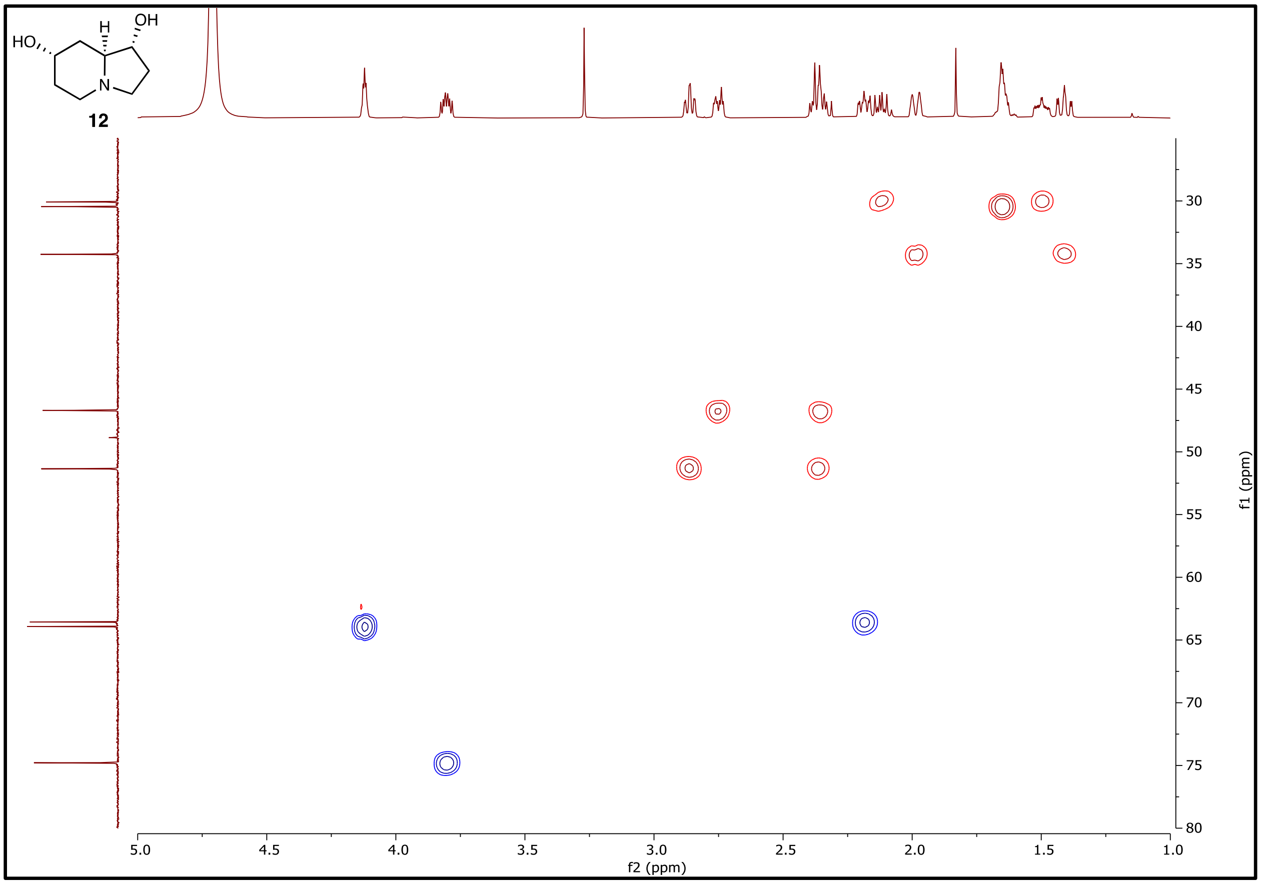

**Figure S61. $^1$H-$^{13}$C HSQC NMR (500 MHz) of 12 in $D_2O$.**

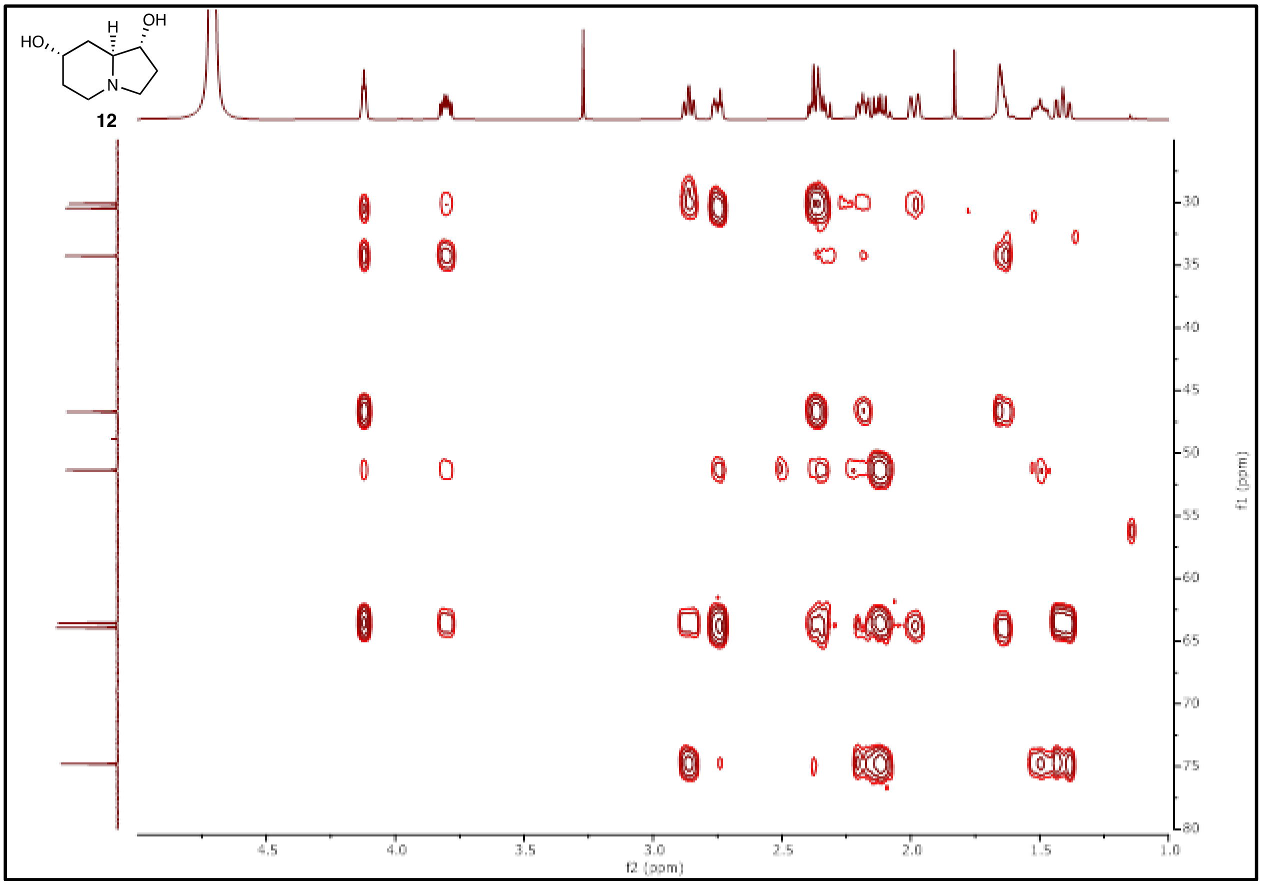

**Figure S62. $^{1}H$-$^{13}C$ HMBC NMR (500 MHz) of 12 in $D_2O$.**

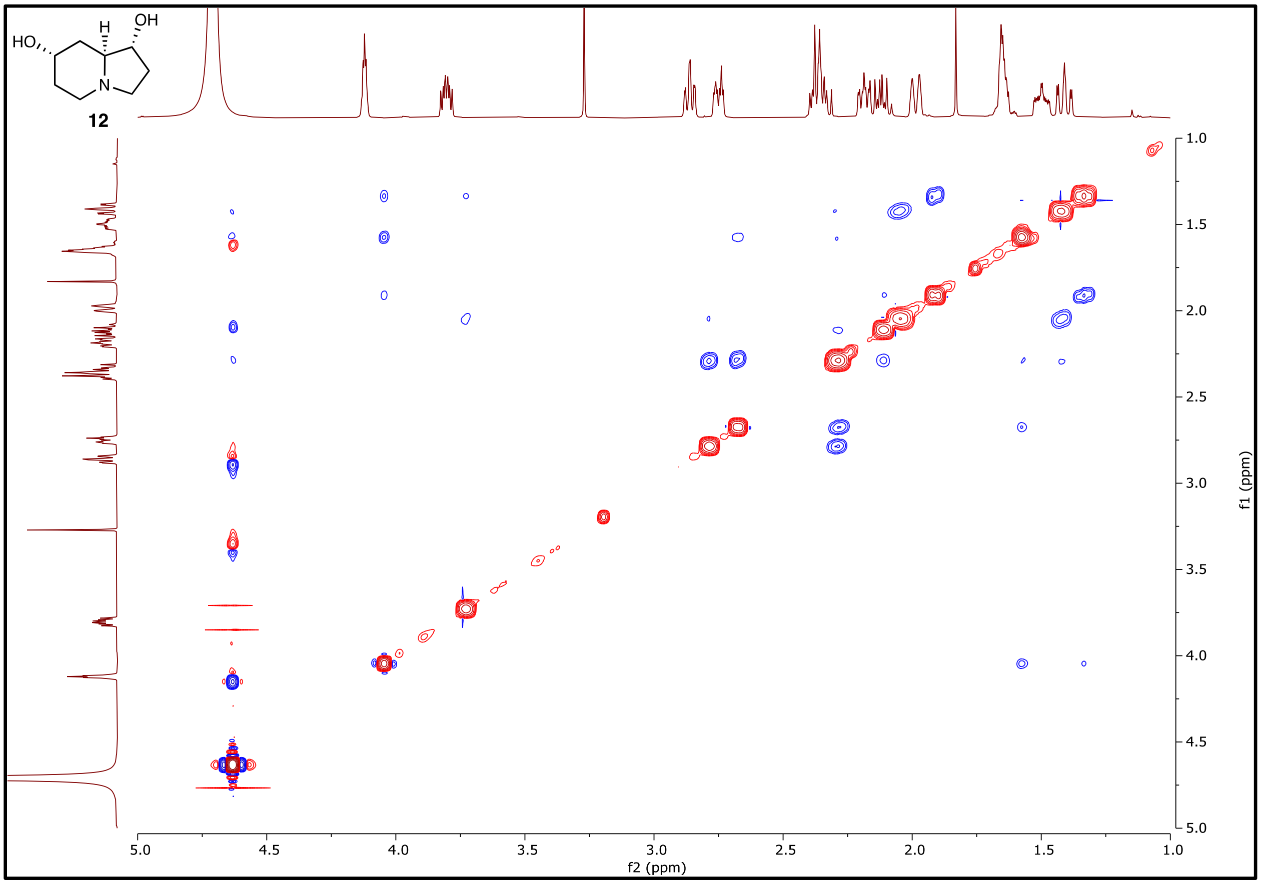

**Figure S63. $^{1}H$-$^{1}H$ NOESY NMR (500 MHz) of 12 in $D_2O$.**

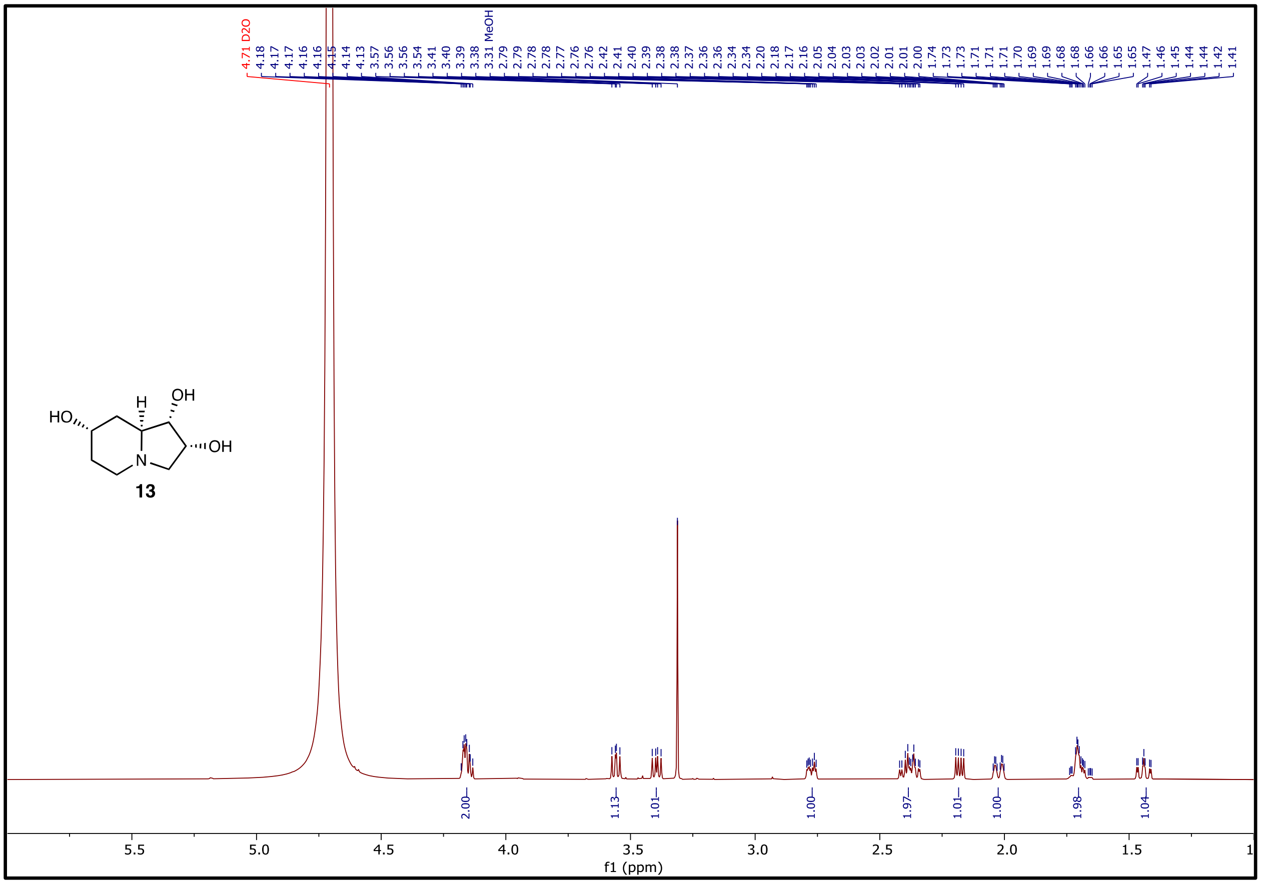

**Figure S64. $^{1}H$ NMR (500 MHz) of 13 in $D_2O$.**

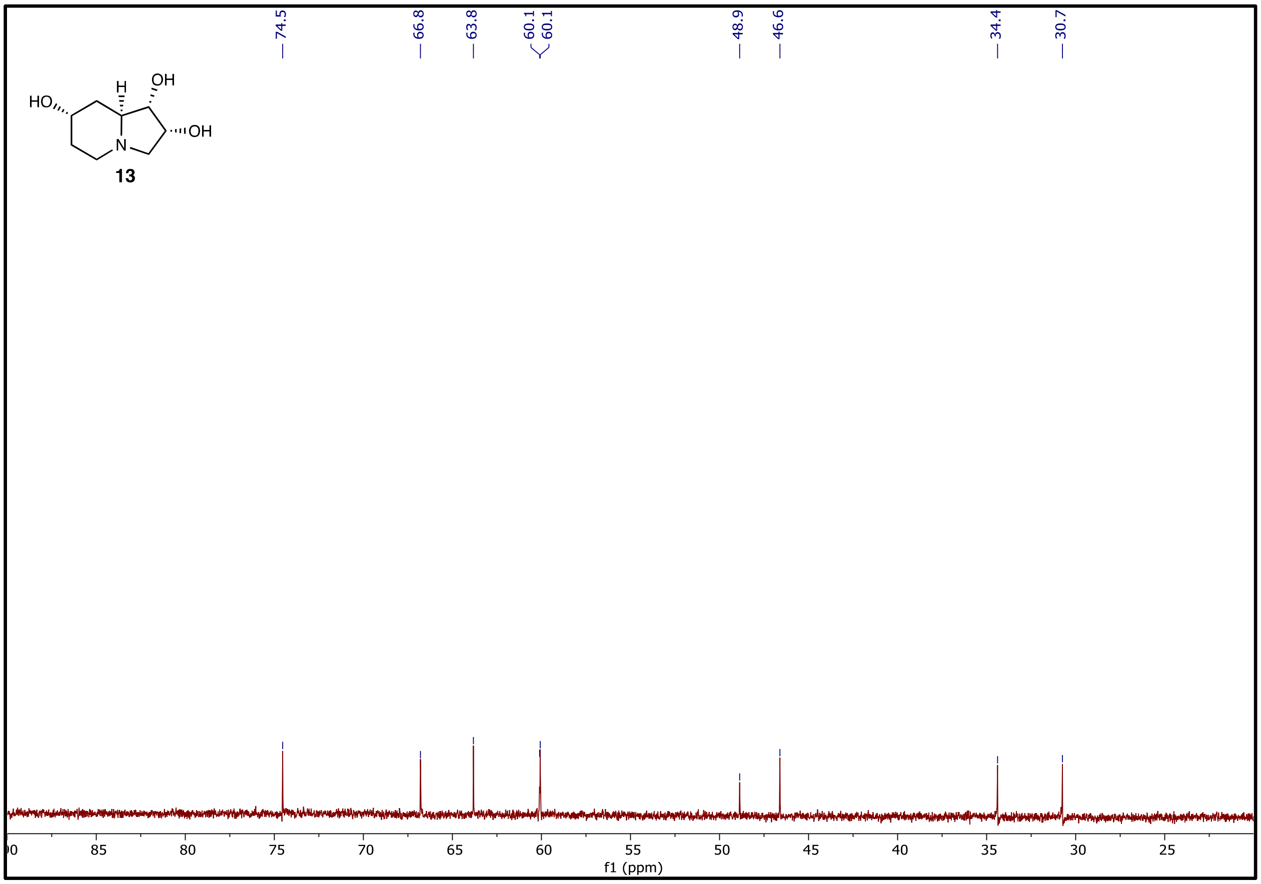

**Figure S65. $^{13}C$ NMR spectrum (126 MHz) of 13 in $D_2O$.**

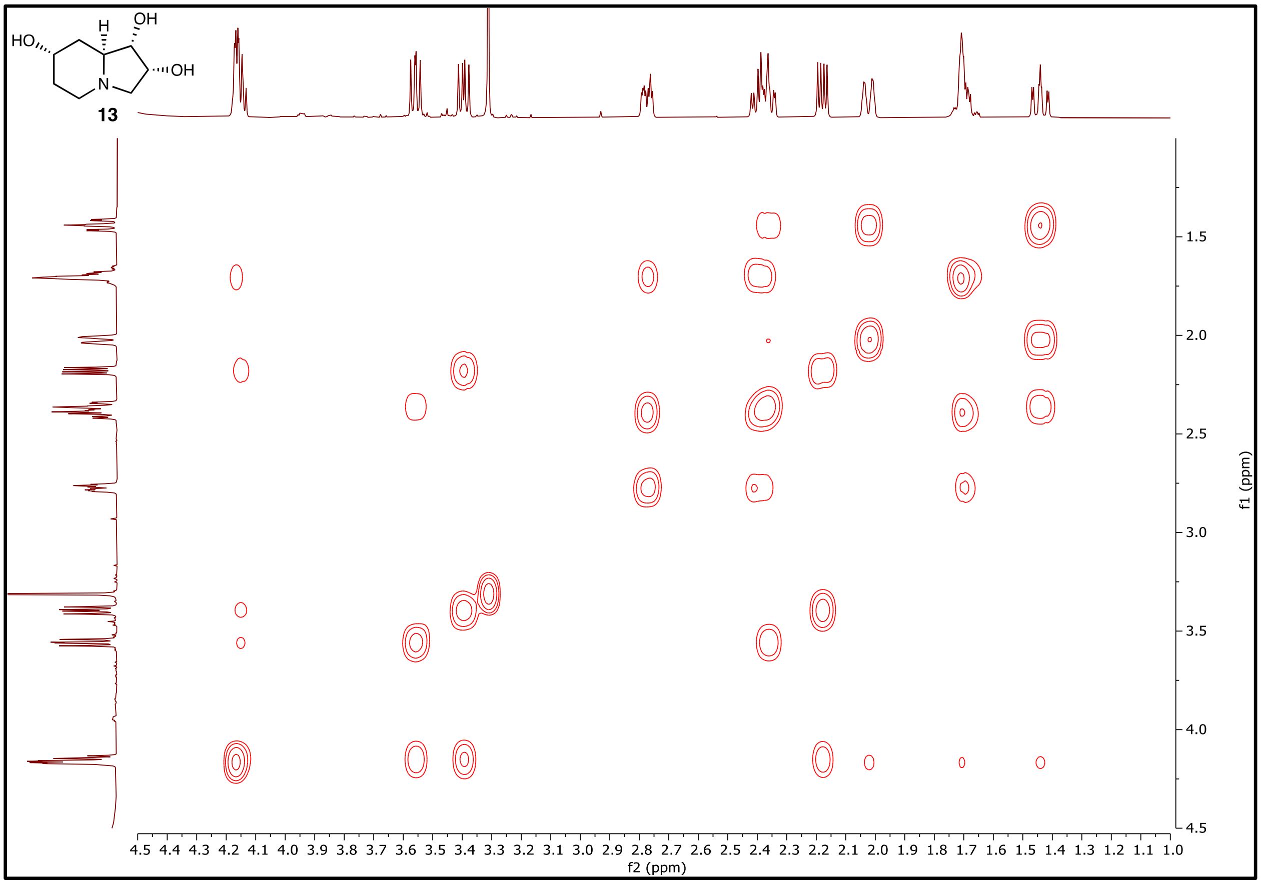

**Figure S66.** **$^1H$-$^1H$ COSY NMR spectrum (500 MHz) of 13 in $D_2O$.**

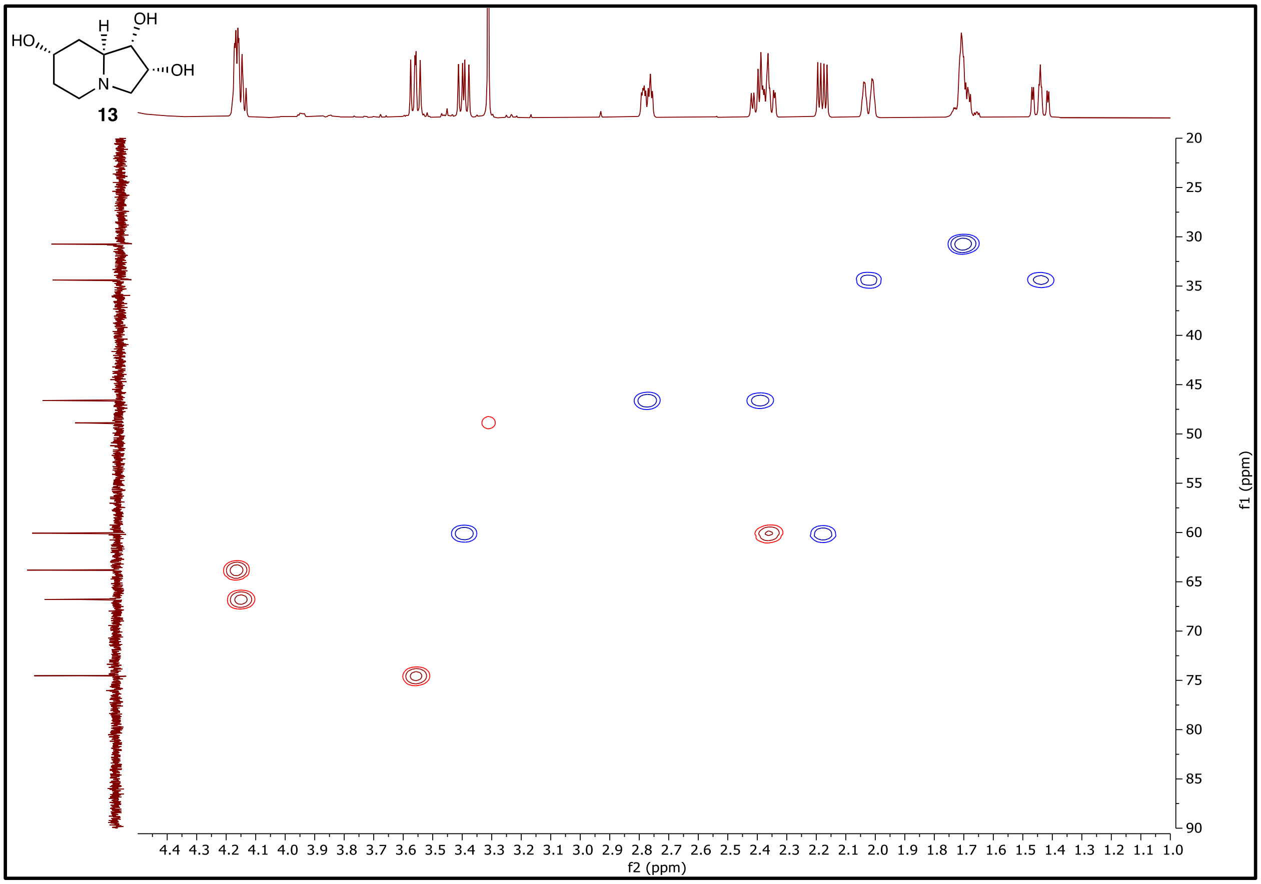

**Figure S67. $^1H$-$^{13}C$ HSQC NMR (500 MHz) of 13 in $D_2O$.**

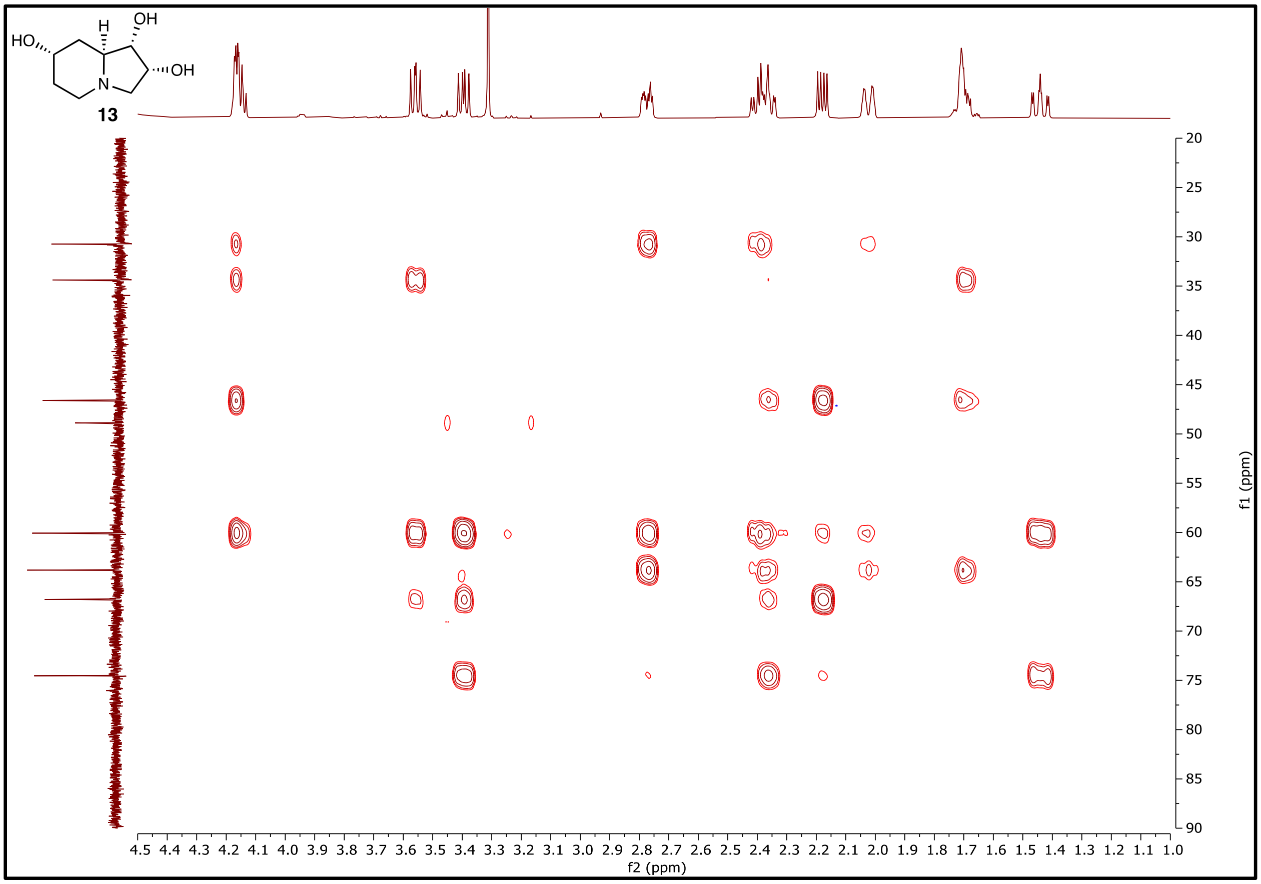

**Figure S68. $^{1}H$-$^{13}C$ HMBC NMR (500 MHz) of 13 in $D_2O$.**

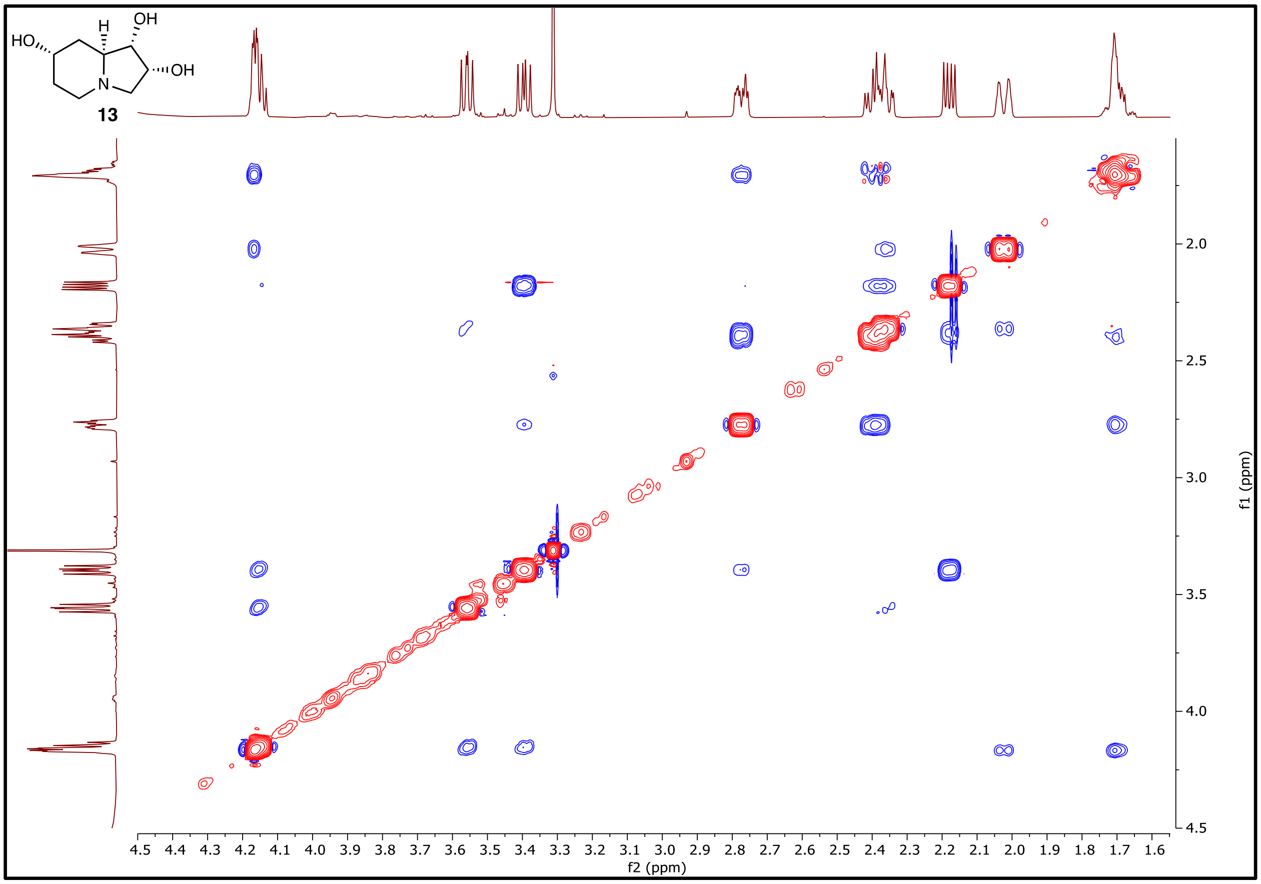

**Figure S69.** **$^1$H-$^1$H NOESY NMR (500 MHz) of 13 in $D_2O$.**

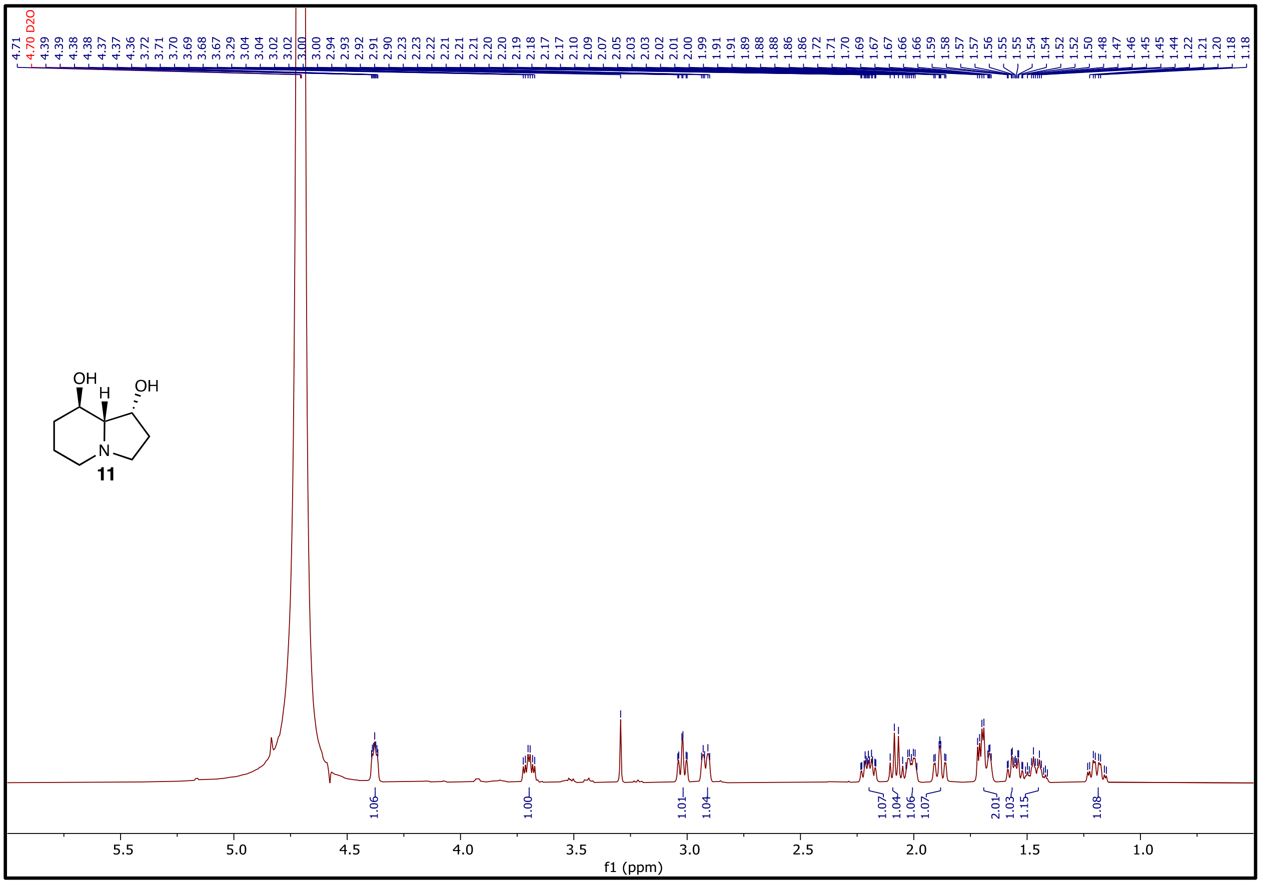

**Figure S70. $^1$H NMR (500 MHz) of 11 in $D_2O$.**

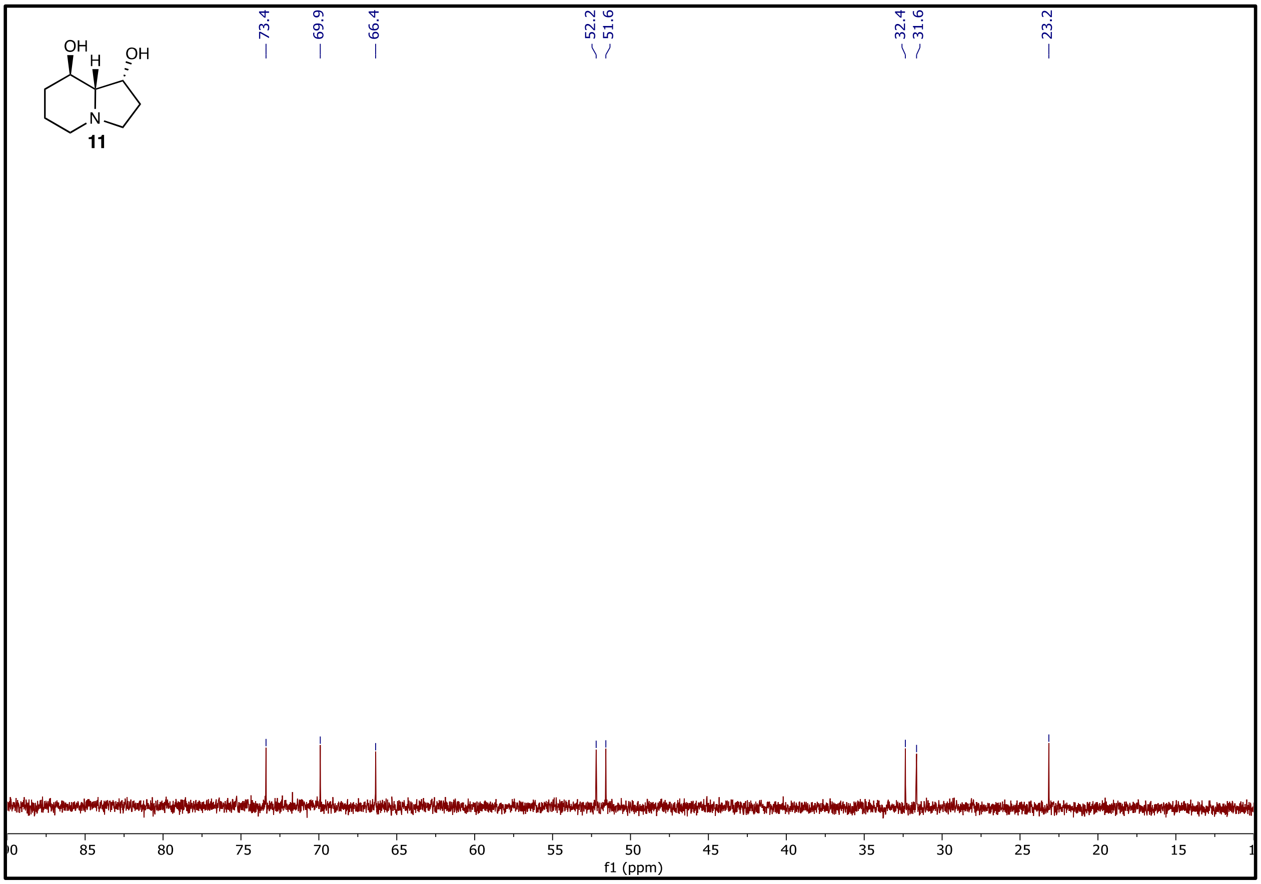

**Figure S71. $^{13}C$ NMR spectrum (126 MHz) of 11 in $D_2O$.**

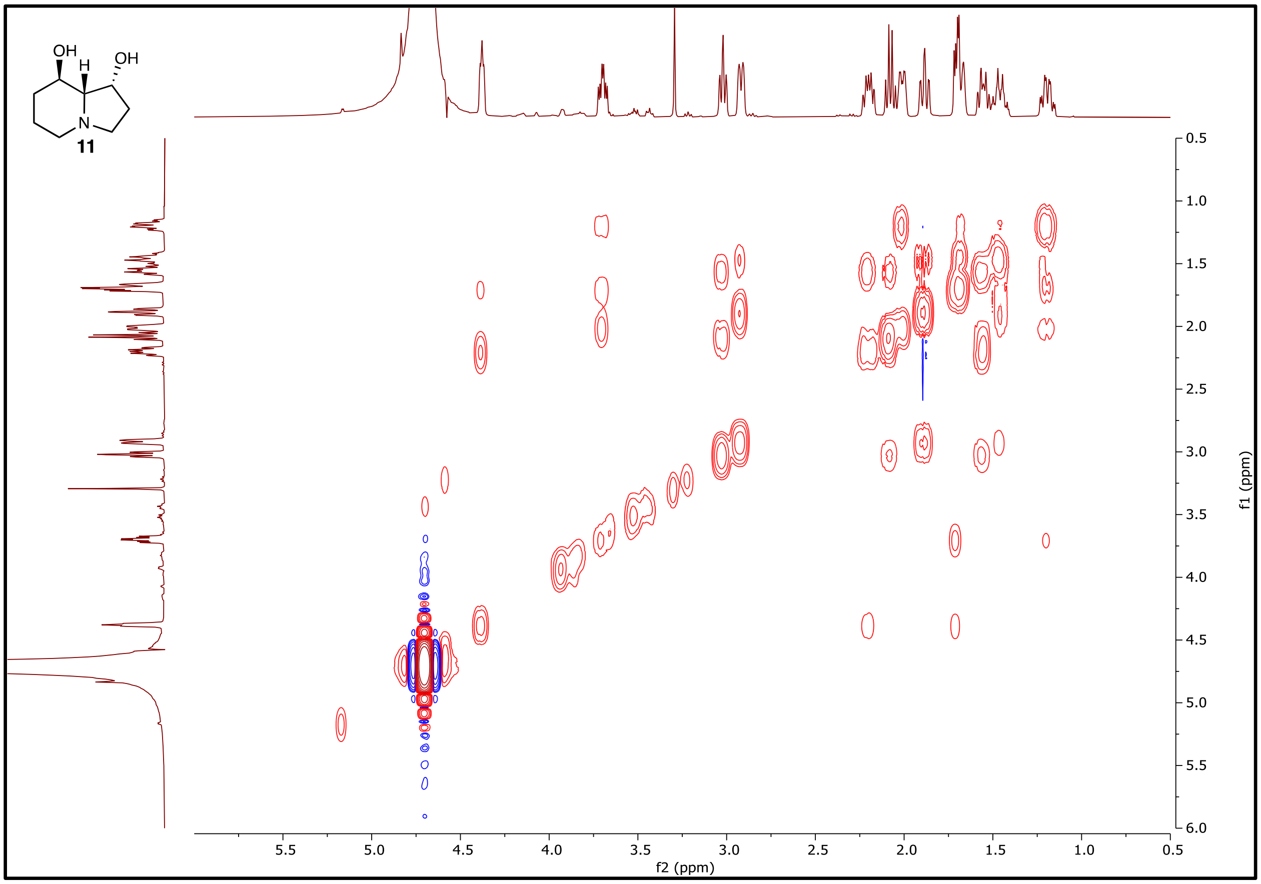

**Figure S72. $^1H$-$^1H$ COSY NMR spectrum (500 MHz) of 11 in $D_2O$.**

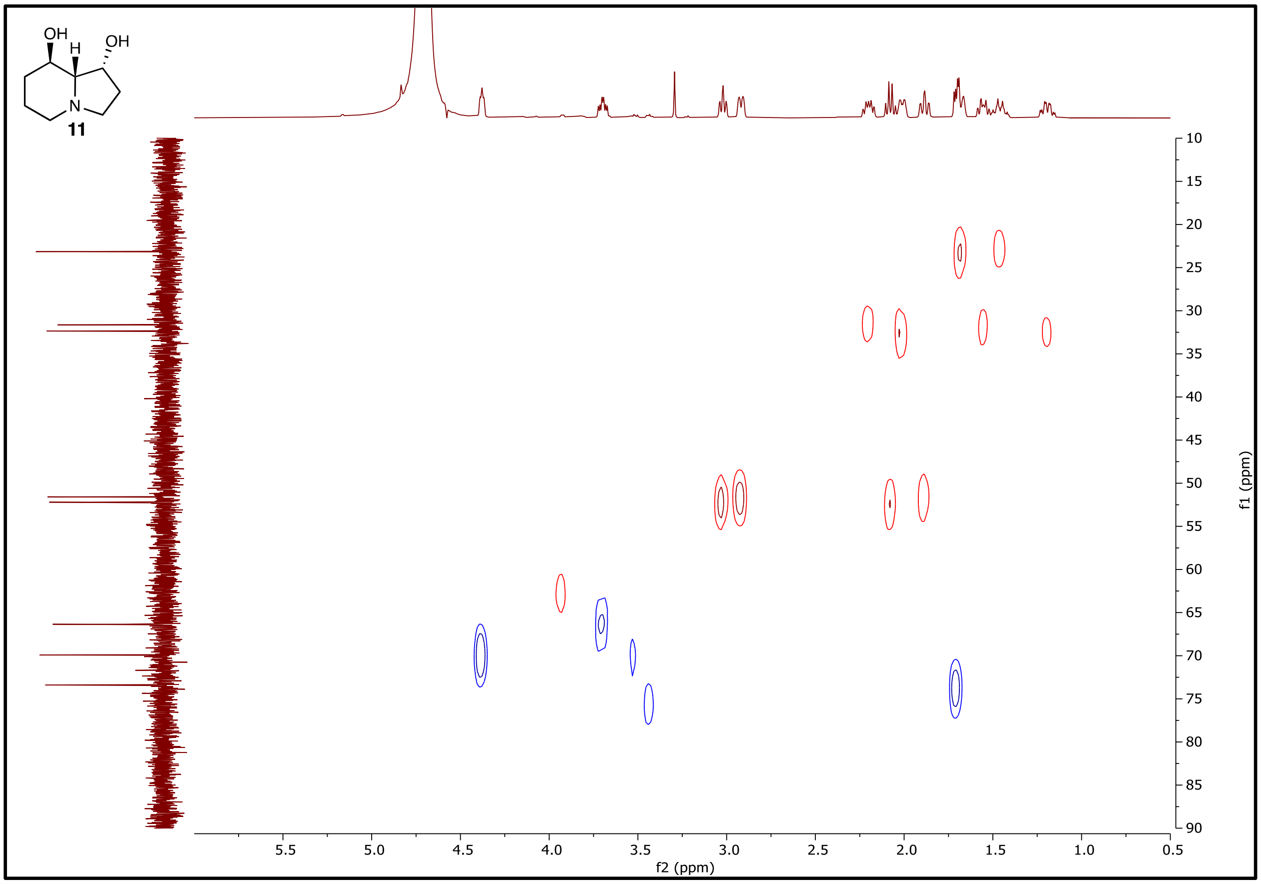

**Figure S73. $^1H$-$^{13}C$ HSQC NMR (500 MHz) of 11 in $D_2O$.**

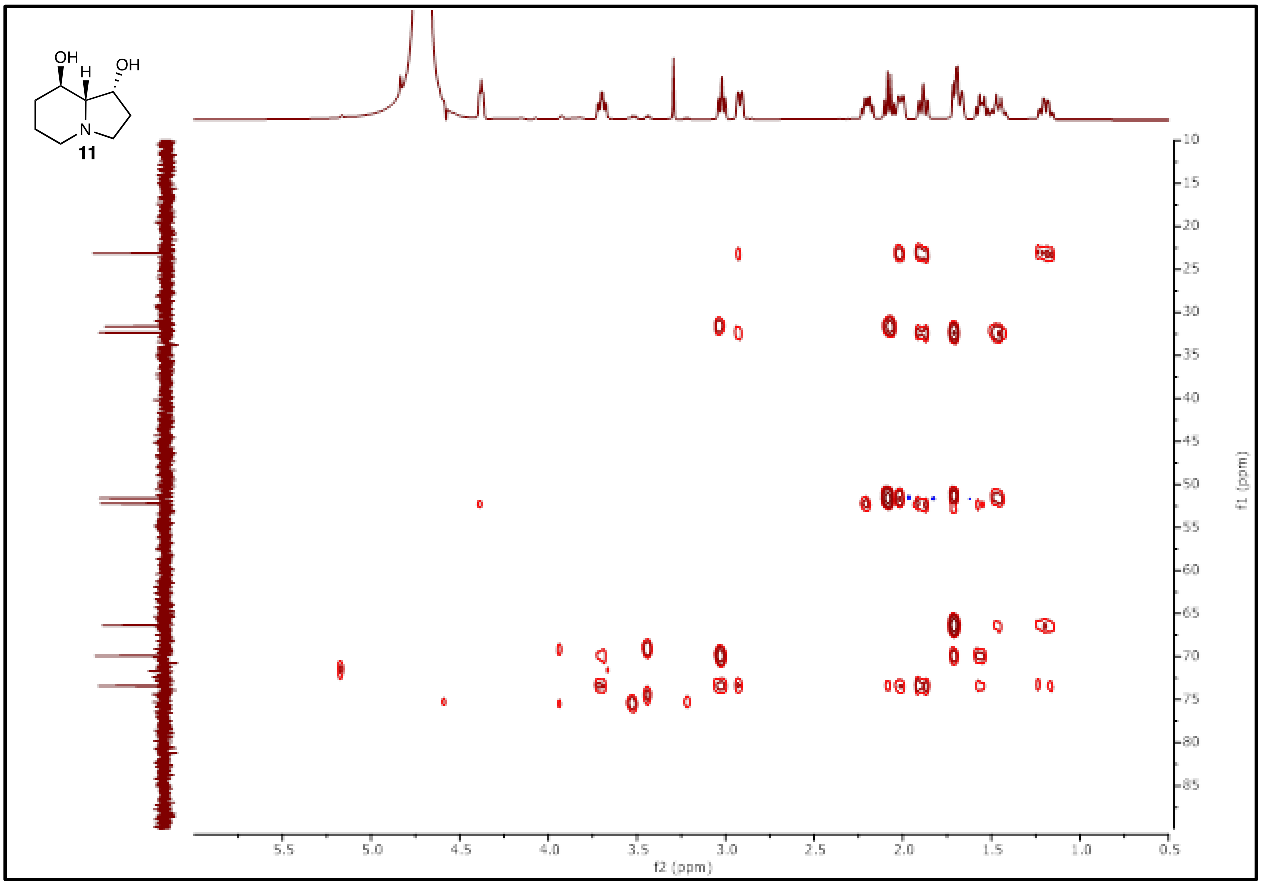

**Figure S74. $^1H$-$^{13}C$ HMBC NMR (500 MHz) of 11 in $D_2O$.**

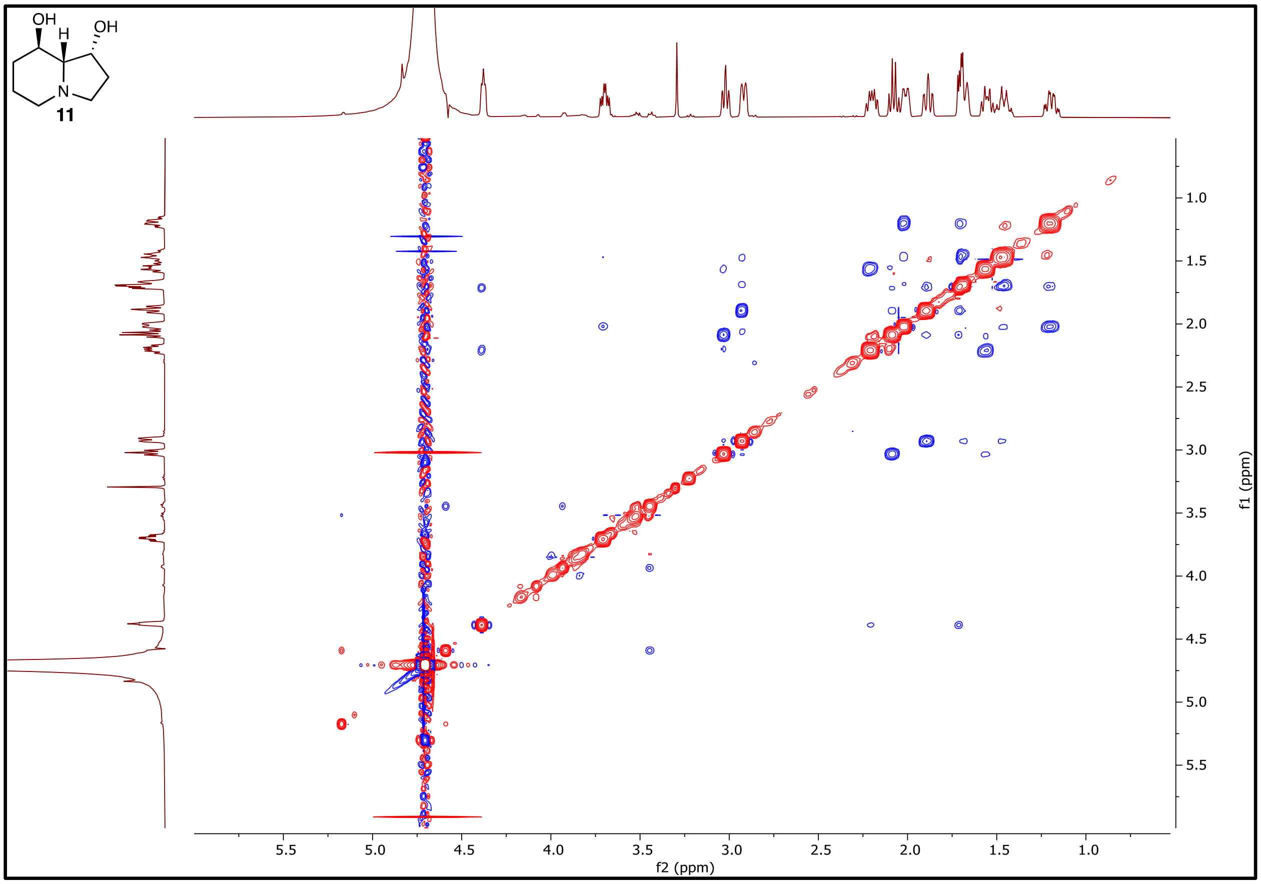

**Figure S62.** $^1H$-$^1H$ **NOESY NMR (500 MHz) of 11 in** $D_2O$**.**

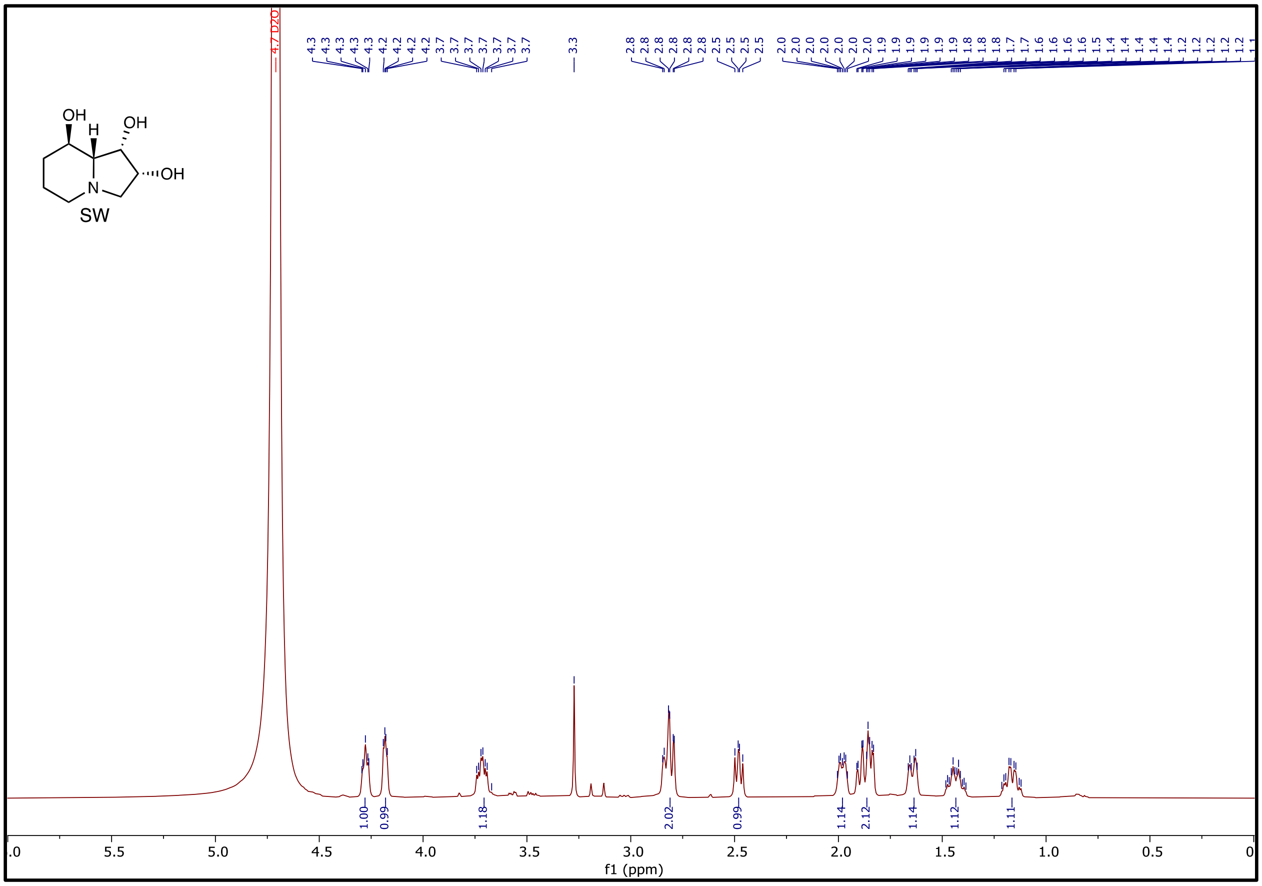

**Figure S74. ${}^{1}H$ NMR (500 MHz) of SW in $D_2O$.**

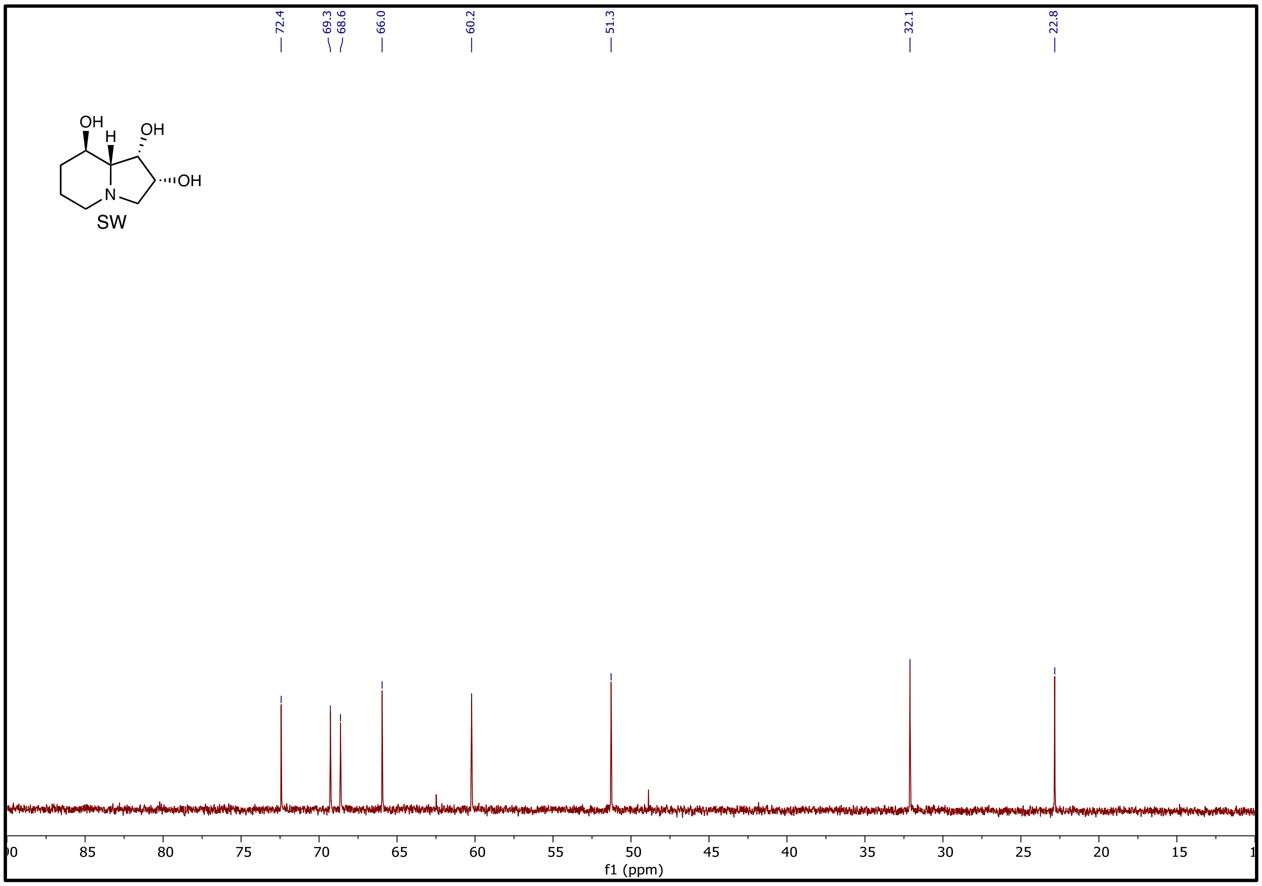

**Figure S75.** $^{13}$**C NMR spectrum (126 MHz) of SW in** $D_2O$**.**

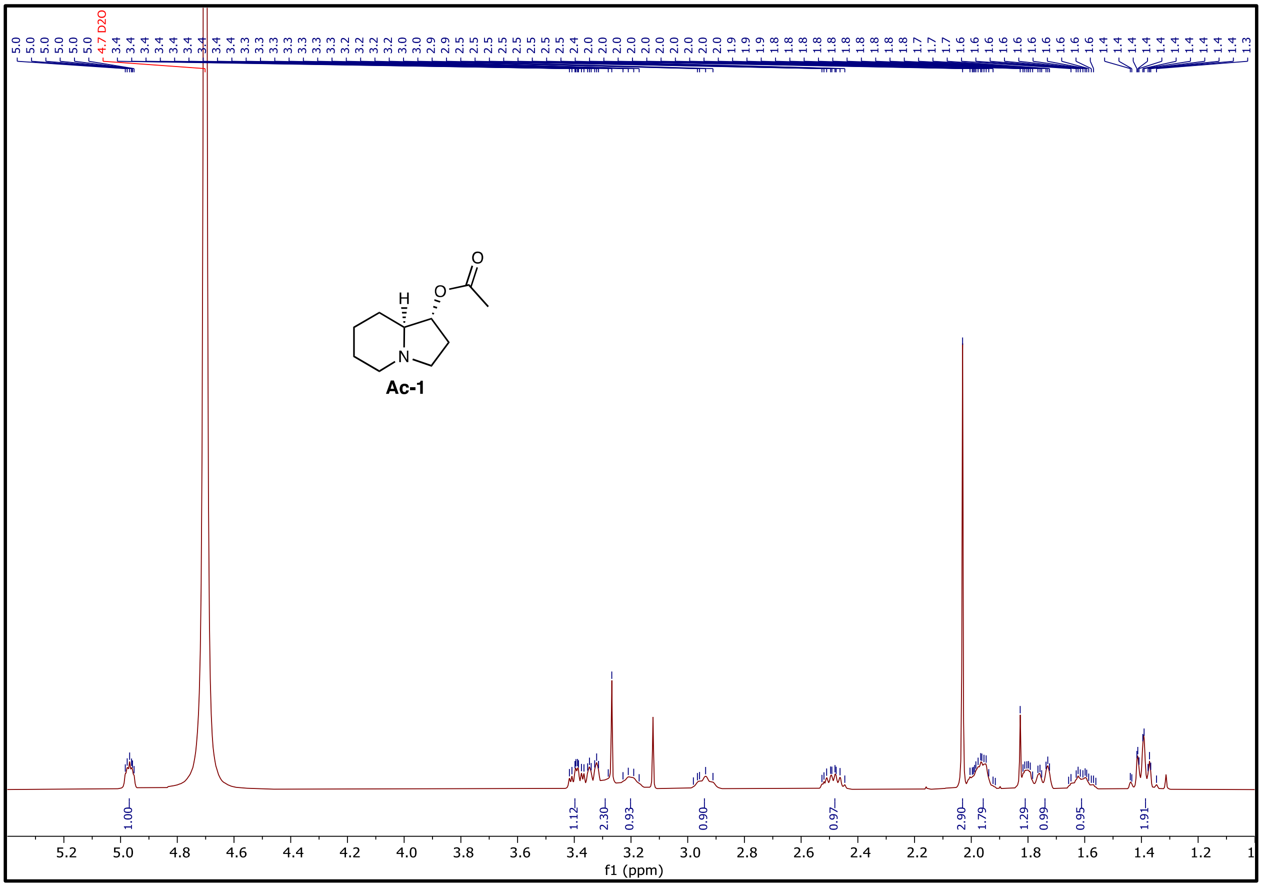

**Figure S76. $^1$H NMR (500 MHz) of 1-Ac in $D_2O$.**

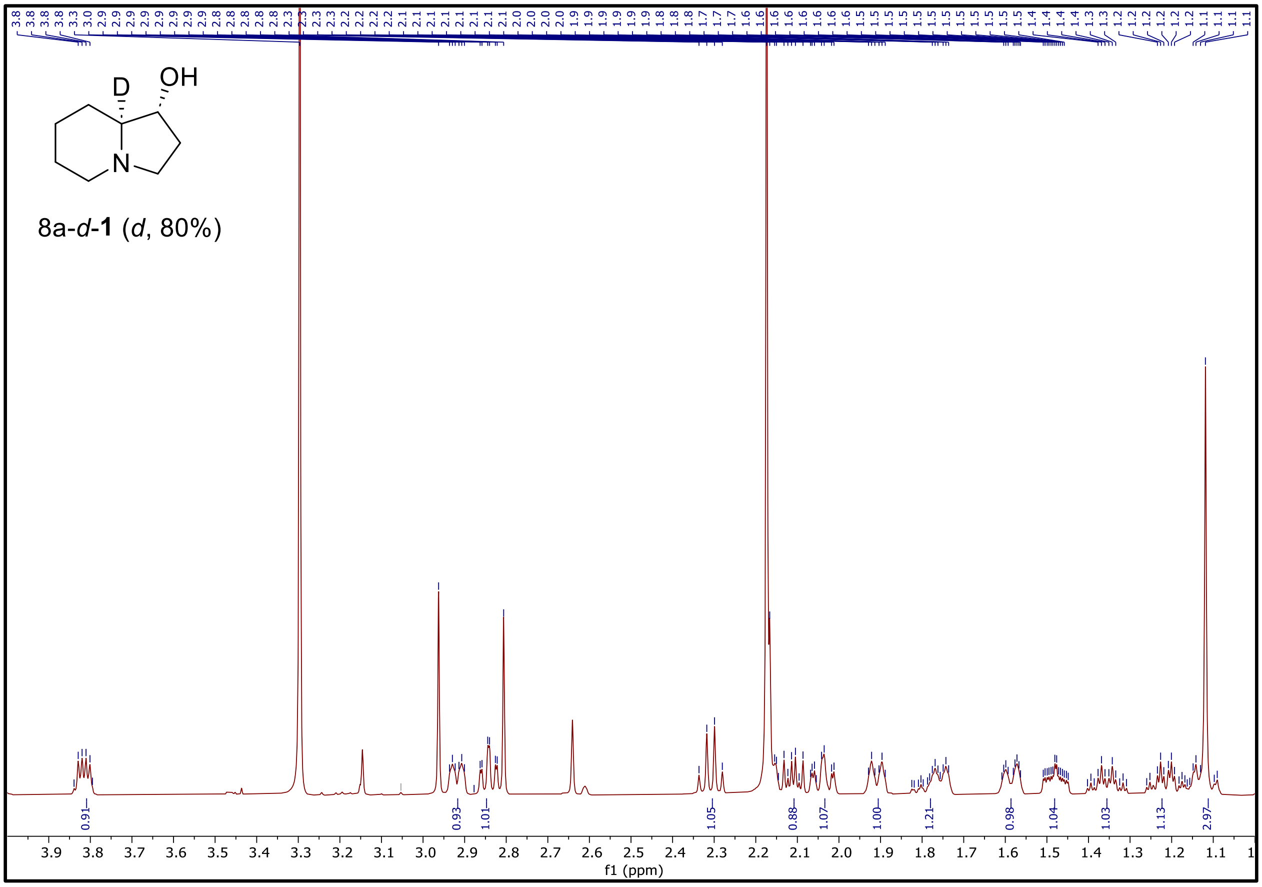

**Figure S77. [1]H NMR (500 MHz) of 8*a*-*d*-1 (*d*, 80%) in $D_2O$.**

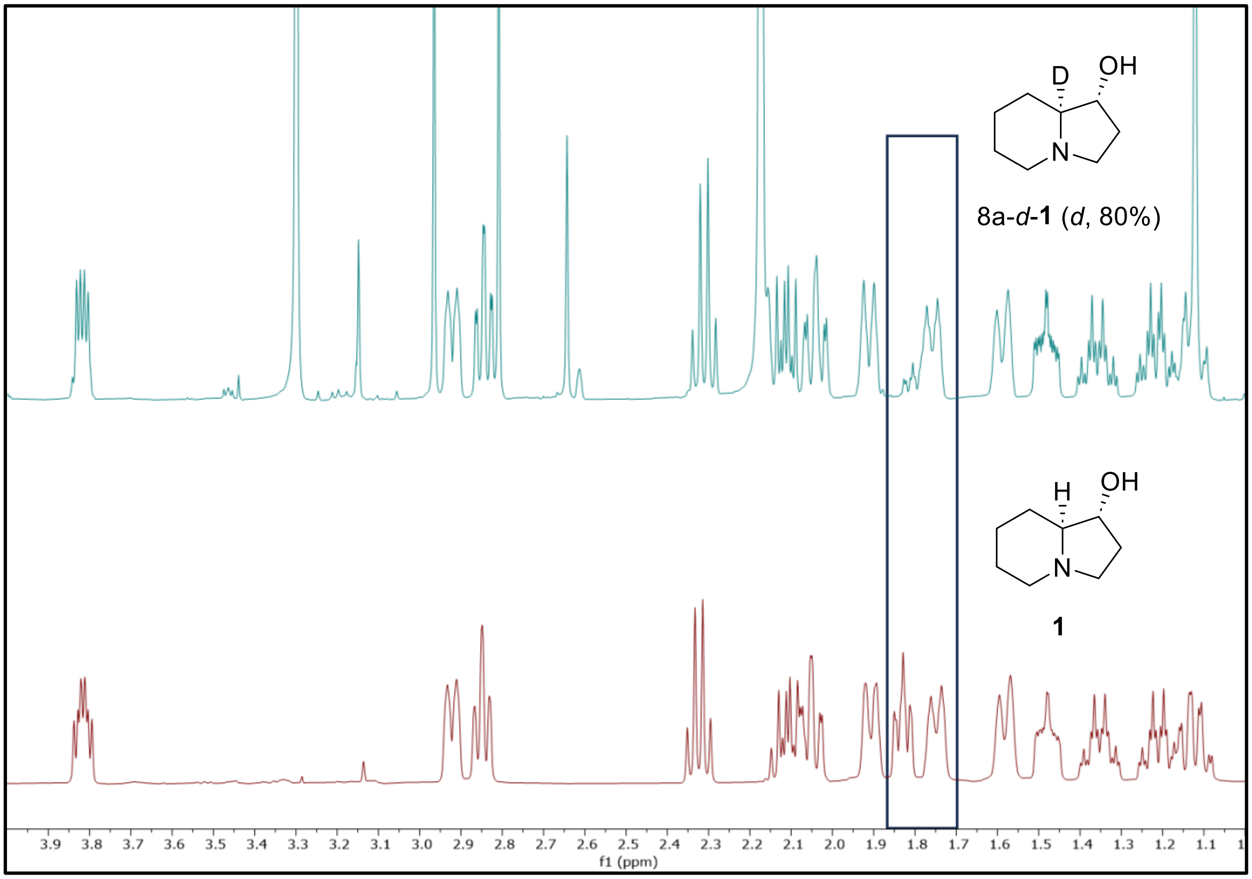

**Figure S78. The stack of $^{1}$H NMR (500 MHz) of 8a-*d*-1 (*d*, 80%, upper) and 1 (bottom) in $D_2O$.**

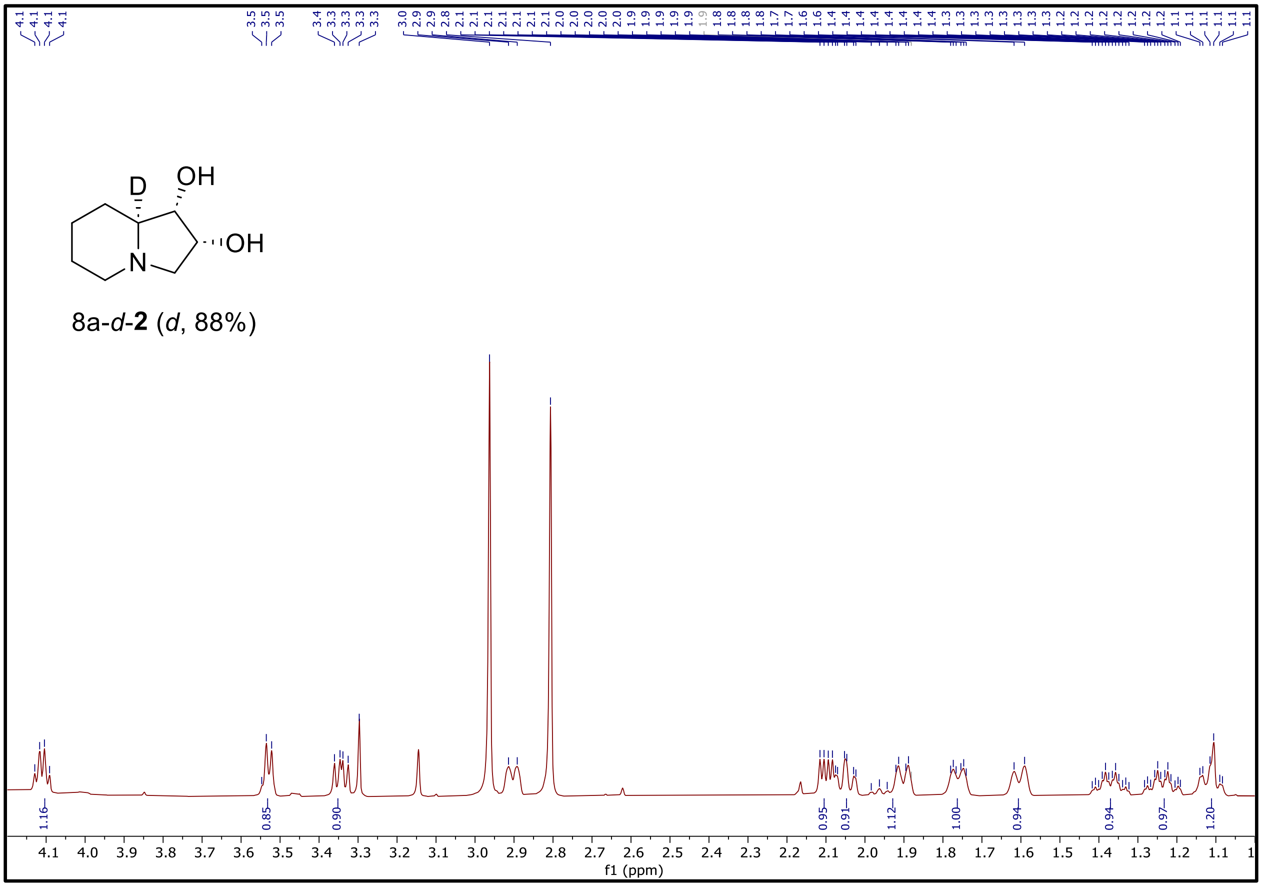

**Figure S79.** $^1$**H NMR (500 MHz) of 8*a*-*d*-2 (*d*, 88%) in** $D_2O$**.**

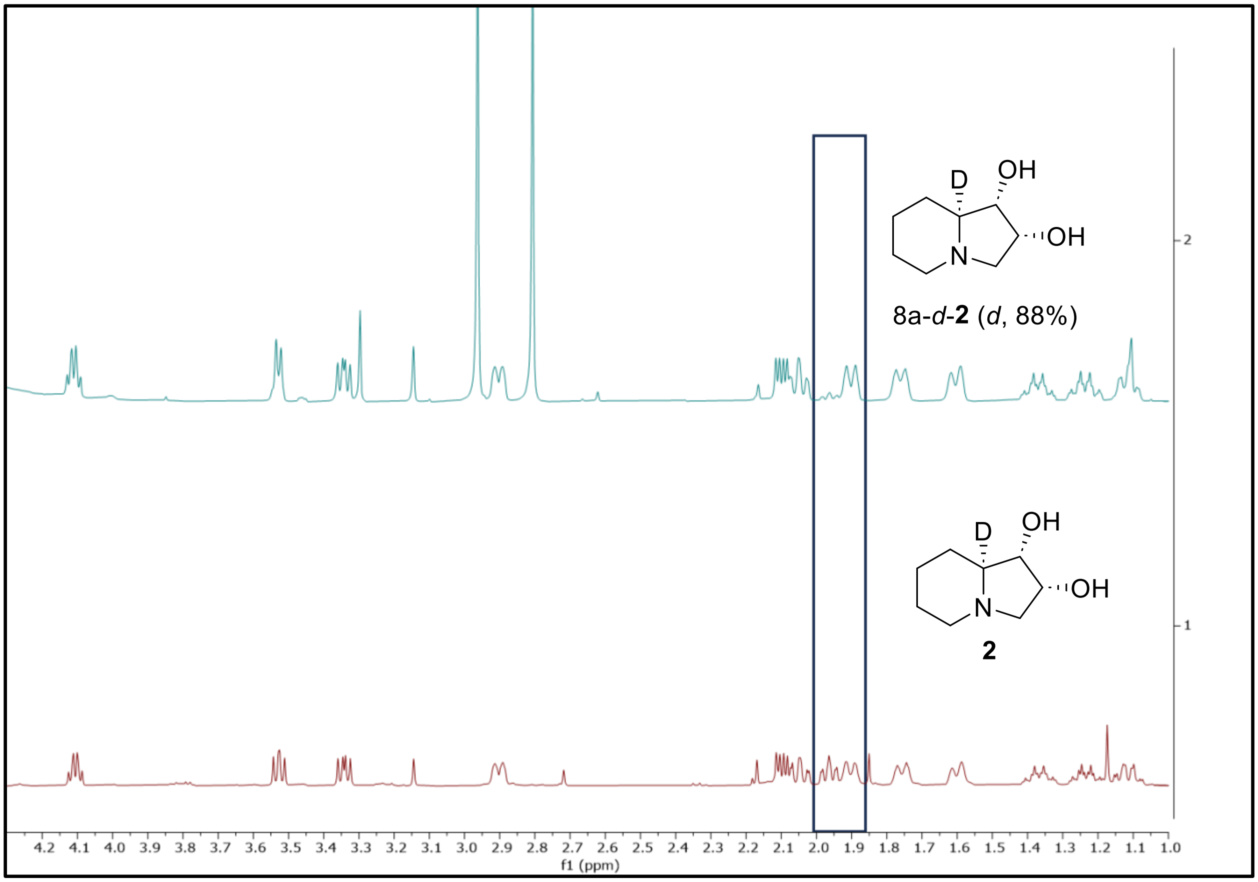

**Figure S80. The stack of $^{1}H$ NMR (500 MHz) of 8a-*d*-2 (*d*, 88%, upper) and 2 (bottom) in $D_2O$.**

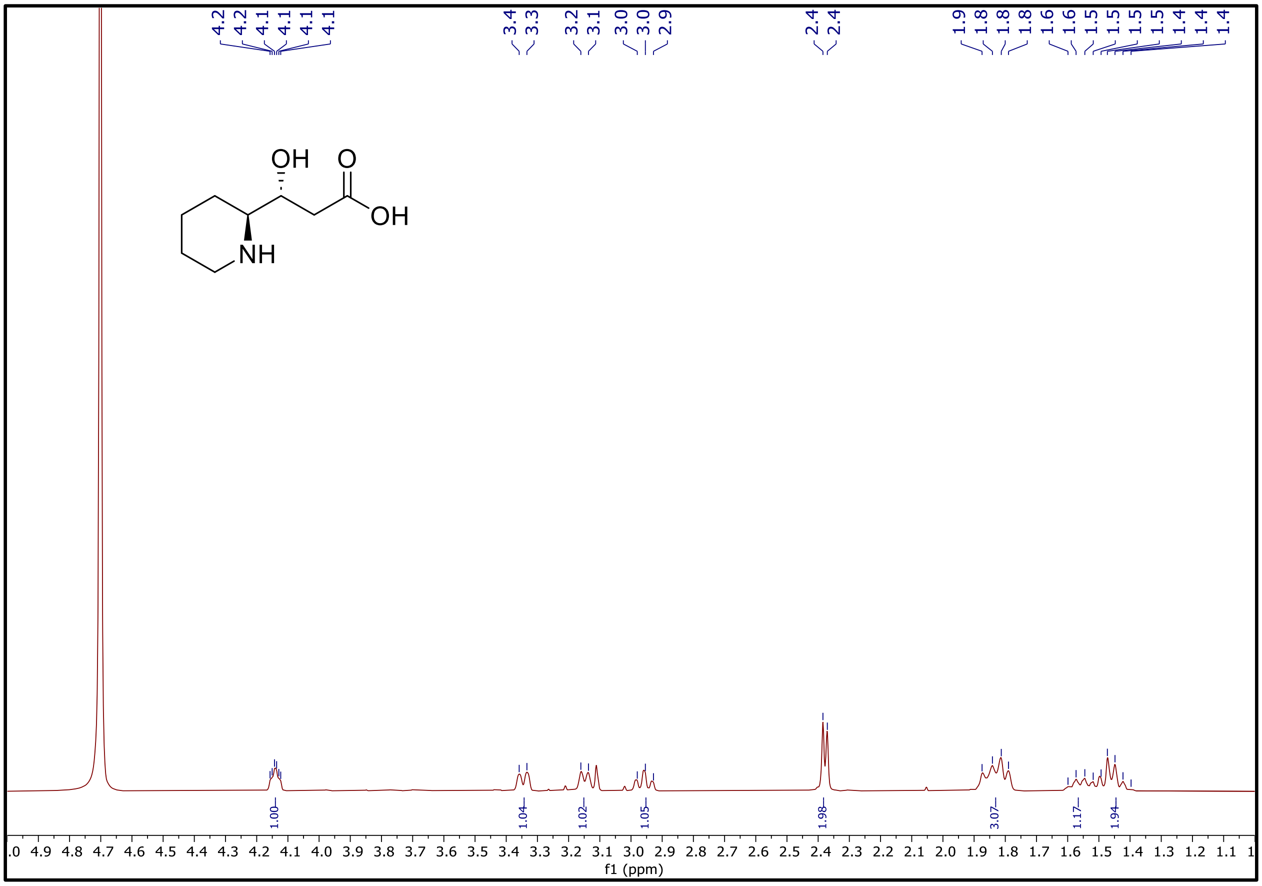

**Figure S81. [1]H NMR (500 MHz) of 3’ in $D_2O$.**

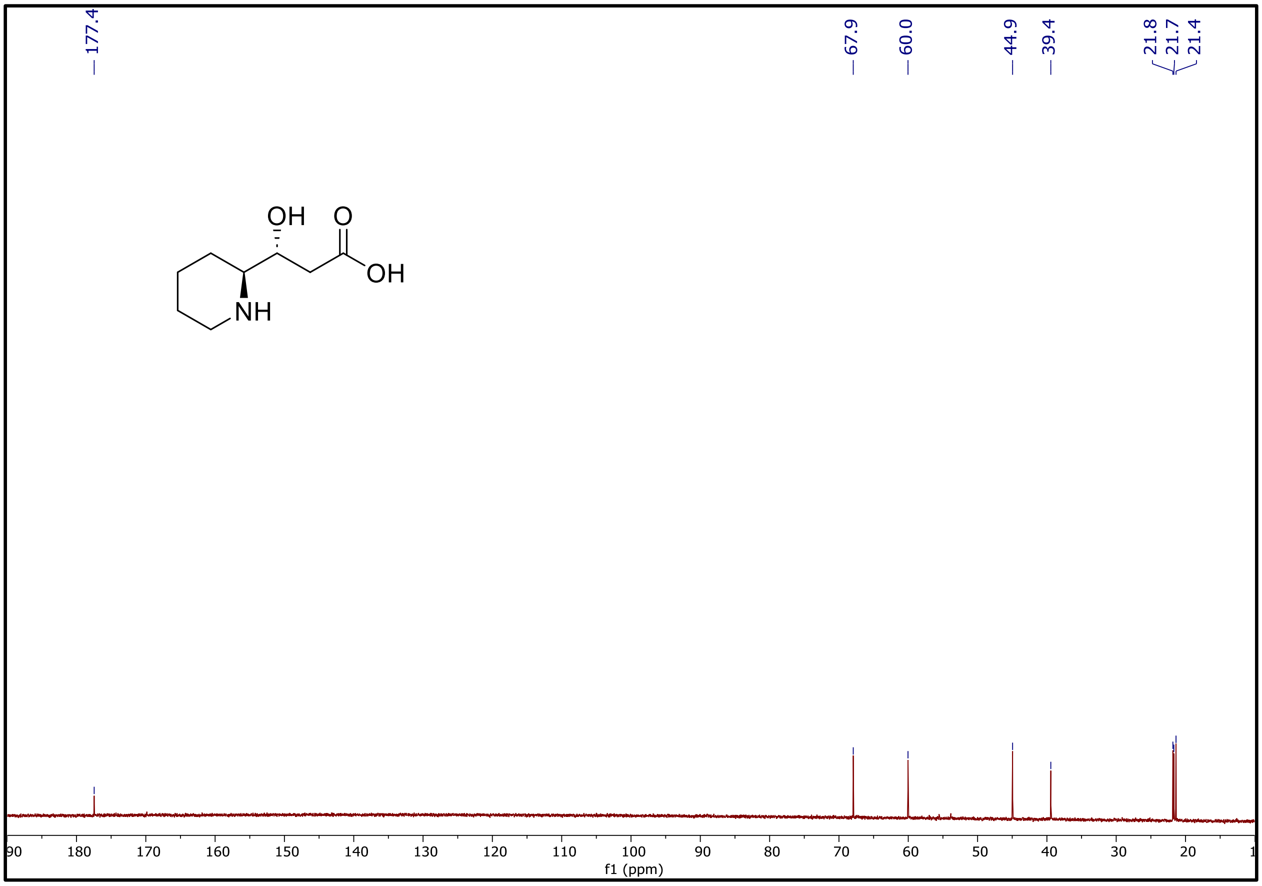

**Figure S82. $^{13}$C NMR (126 MHz) of 3’ in $D_2O$.**

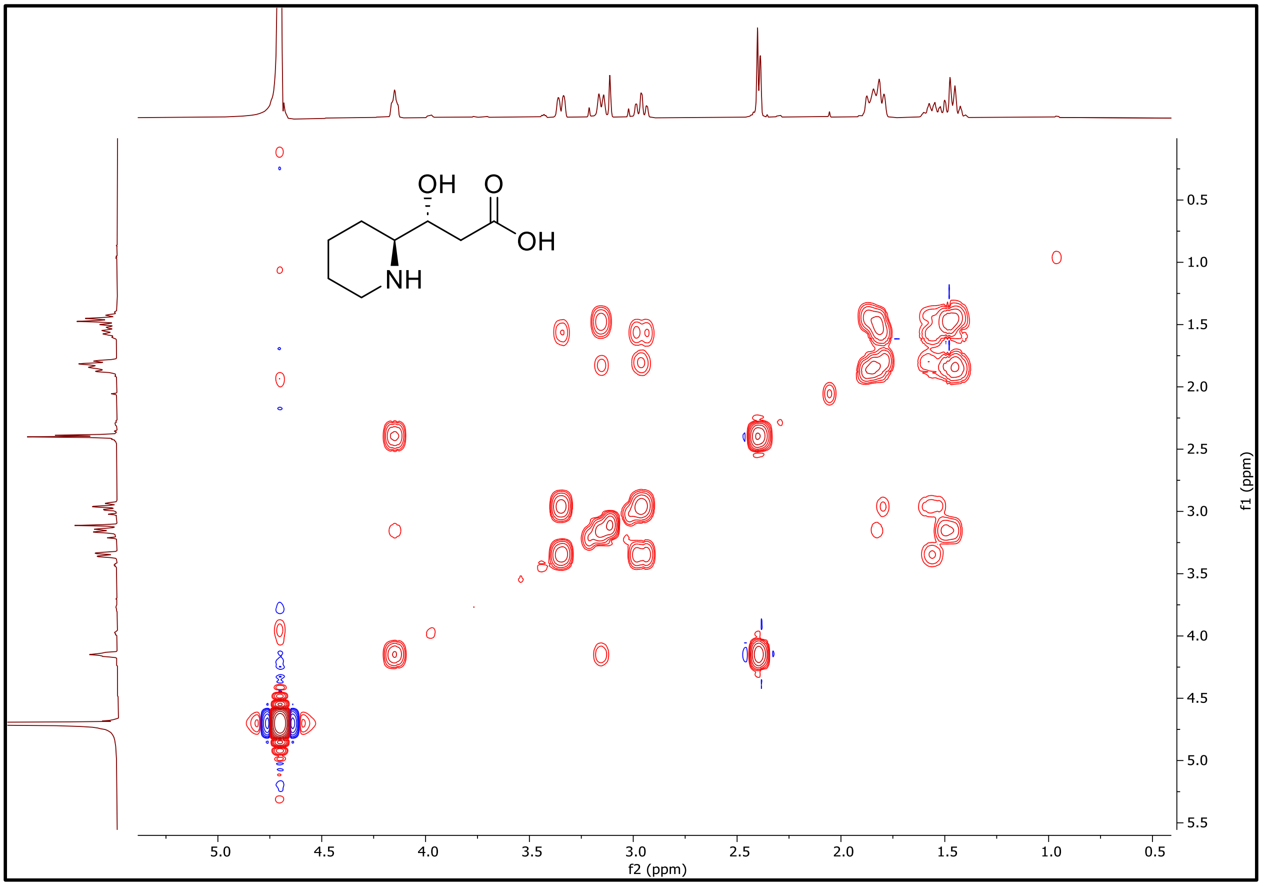

**Figure S83. $^{1}H$-$^{1}H$ Cosy NMR (500 MHz) of 3' in $D_2O$.**

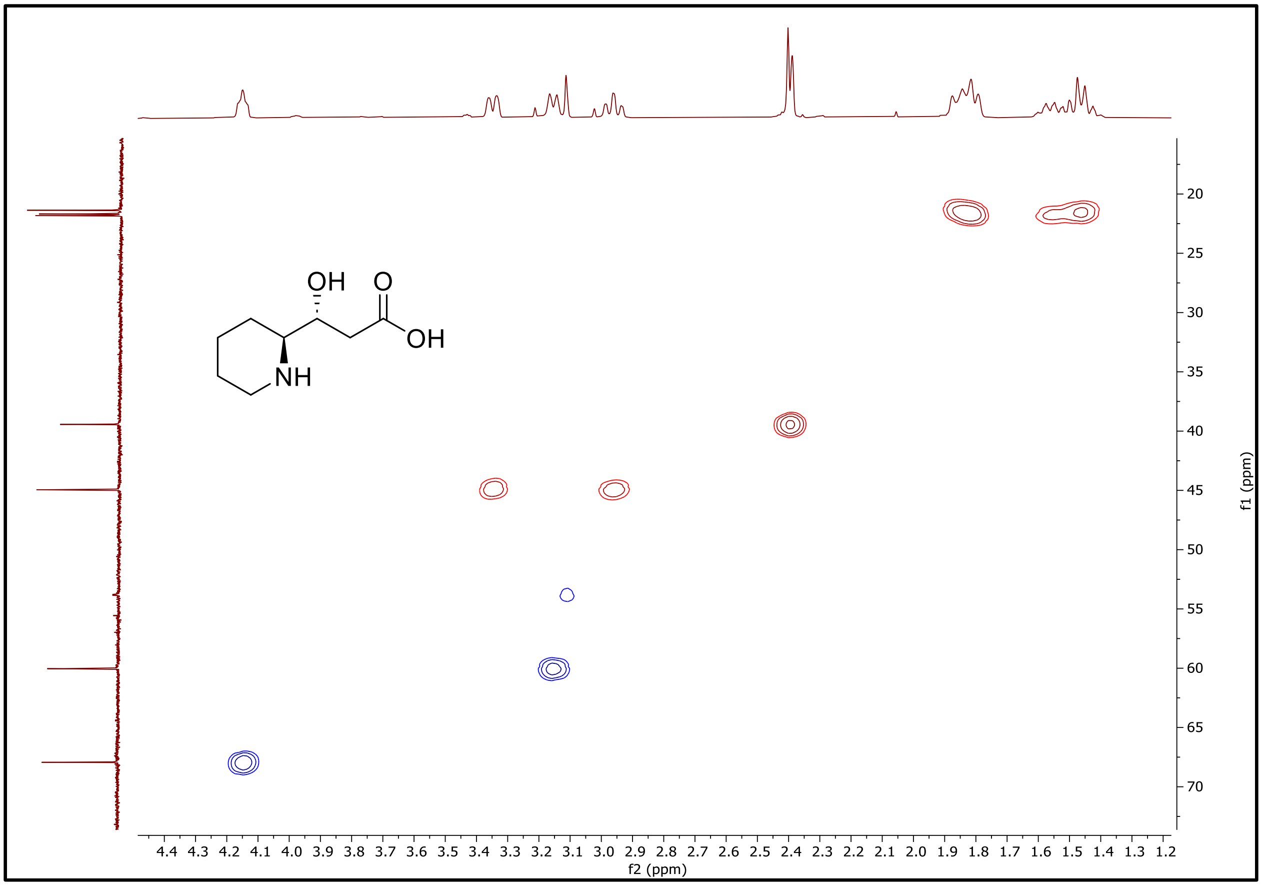

**Figure S84.** $^{1}H$-$^{13}C$ **HSQC NMR (500 MHz) of 3' in $D_2O$.**

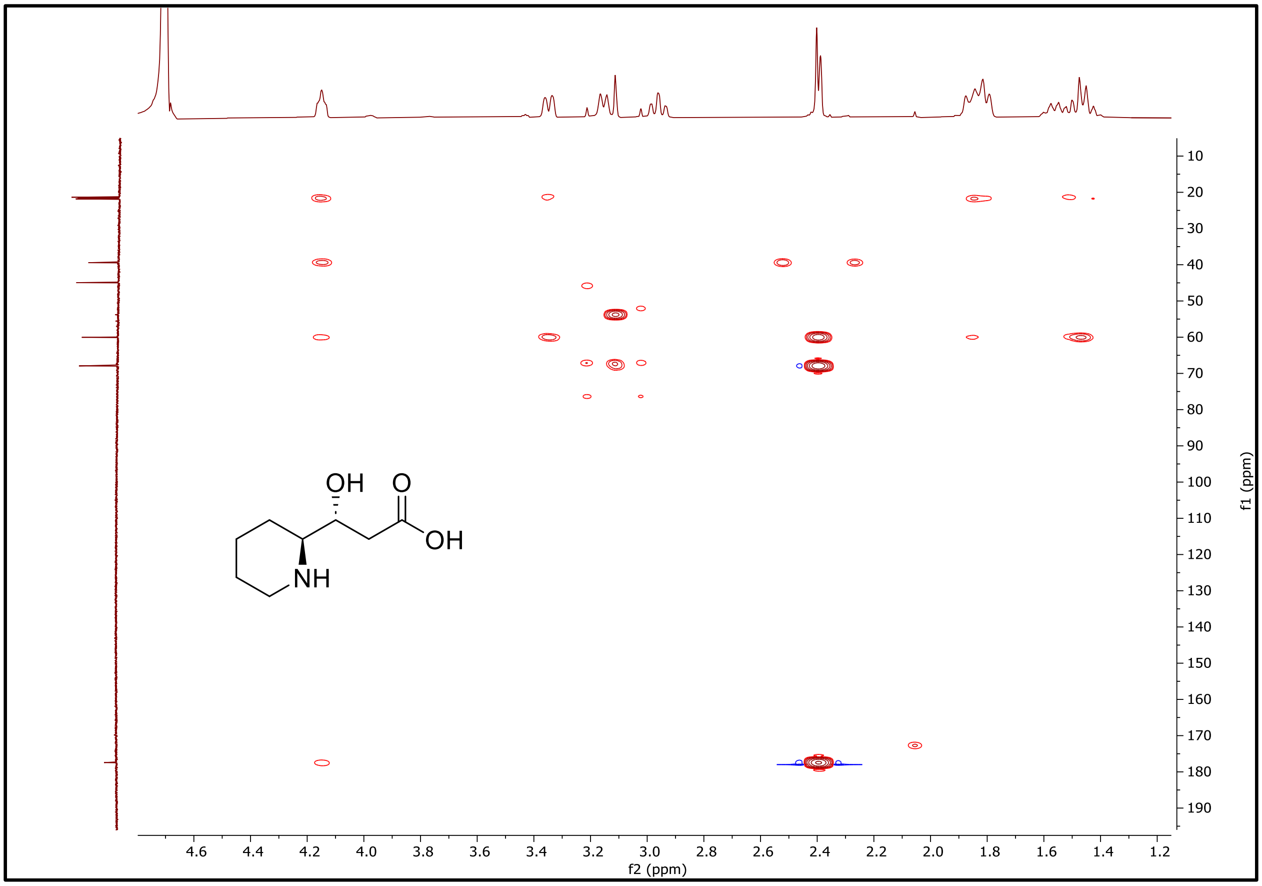

**Figure S85.** $^1H$-$^{13}C$ **HMBC NMR (500 MHz) of 3' in** $D_2O$**.**

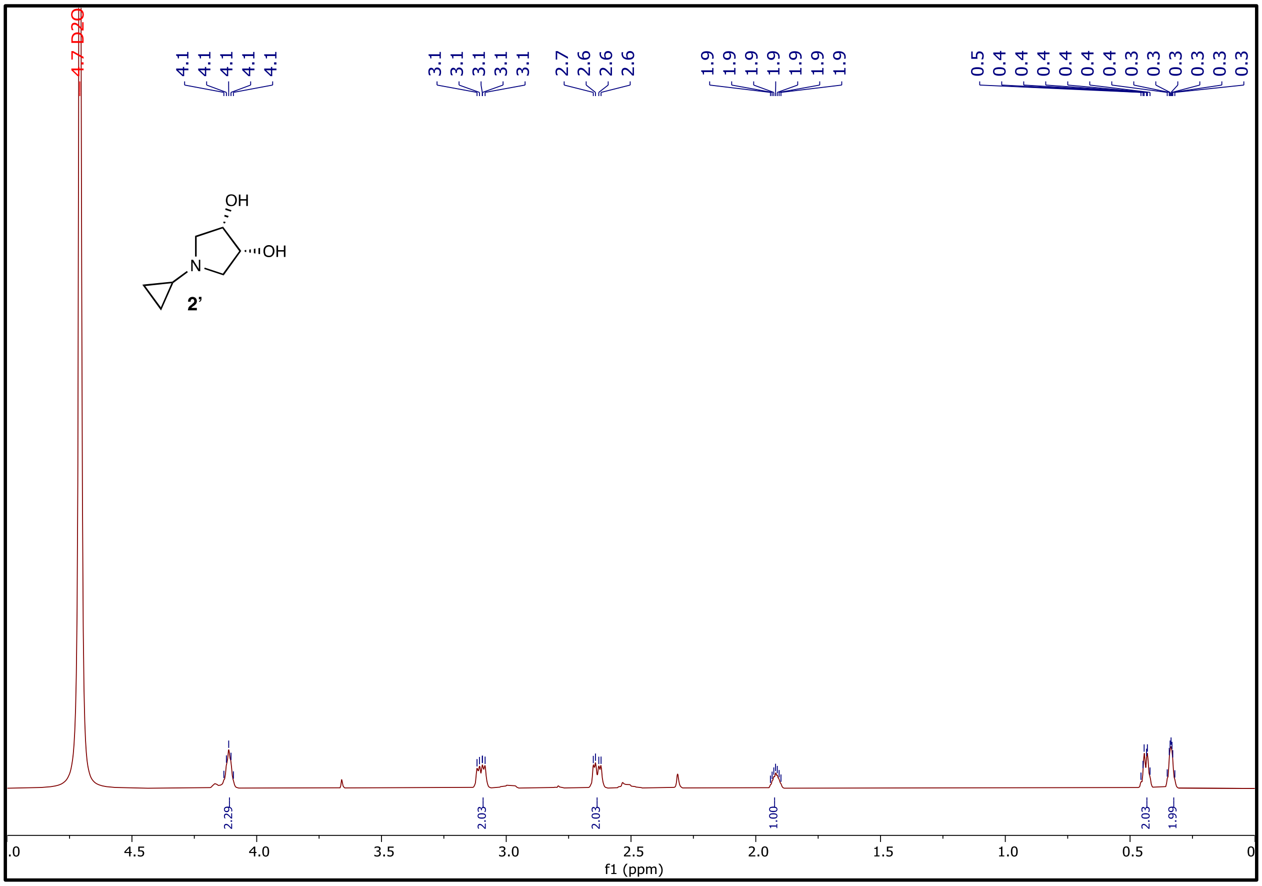

**Figure S86. $^1$H NMR (500 MHz) of 2’ in $D_2O$.**

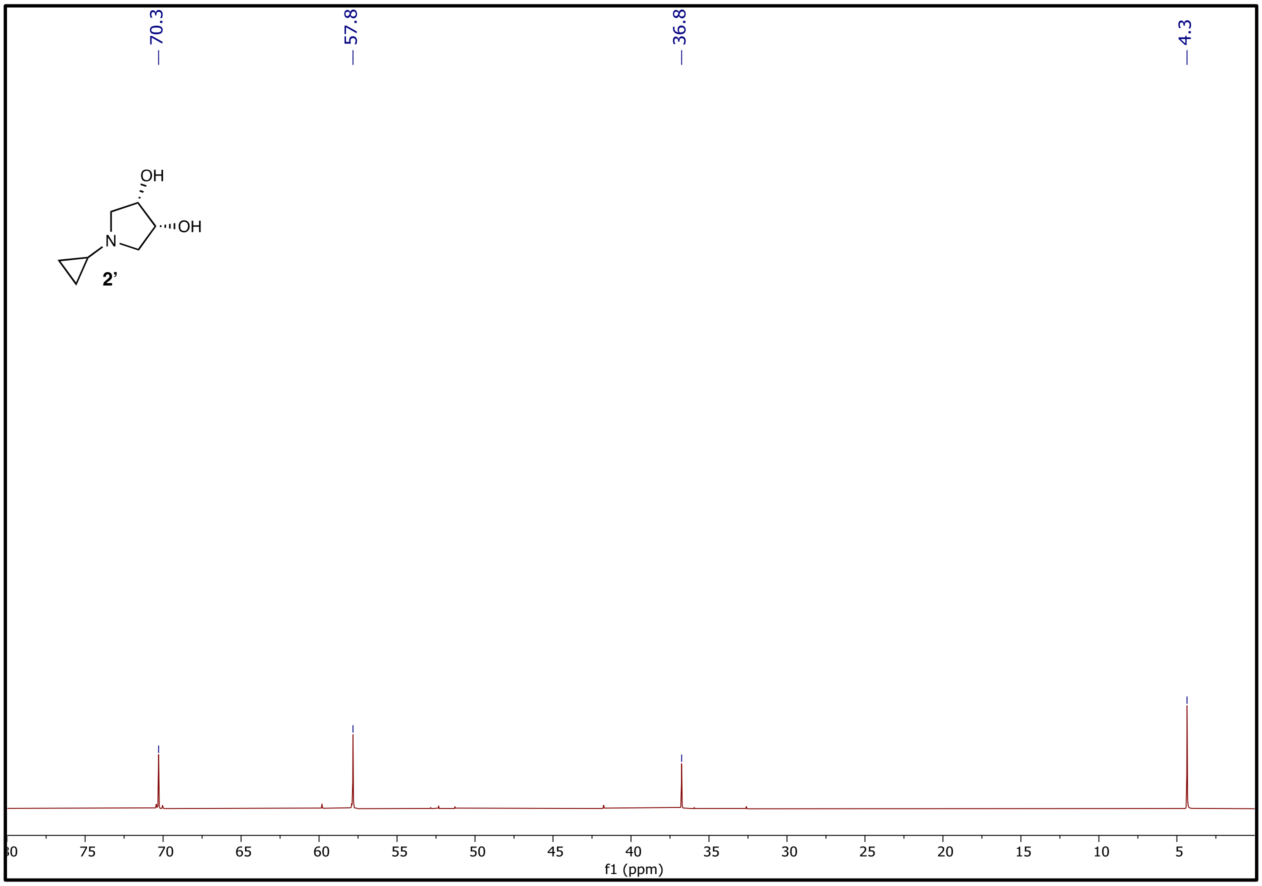

**Figure S87. $^{13}$C NMR (126 MHz) of 2' in $D_2O$.**
